# Chemical tools to Detect and Inhibit IgA1 Proteases in *Haemophilus influenzae*

**DOI:** 10.64898/2026.03.17.712269

**Authors:** Vani Verma, Pooja S. Thomas, Martina Lancieri, Jasper Van den Bos, Adrian Fabisiak, Sarah Peeters, Marie-Stéphanie Aschtgen, Edmund Loh, Ingrid De Meester, Hans De Winter, Pieter Van der Veken, Michaela Prothiwa

## Abstract

Non-encapsulated (“non-typeable”) *Haemophilus influenzae* is a major cause of mucosal infections such as otitis media, conjunctivitis, and exacerbations of chronic obstructive pulmonary disease. Rising antibiotic resistance has increased interest in anti-virulence strategies that reduce pathogenicity without exerting selective pressure for resistance. A key virulence factor of *H. influenzae* is immunoglobulin A1 protease (IgA1P), a secreted serine protease that promotes colonization by cleaving human IgA1 at the hinge region and enabling immune evasion. Despite its therapeutic promise, progress has been limited by the lack of chemical tools to probe IgA1P activity in its native biological environment. Here, we report the first activity-based probes that enable direct detection of active IgA1Ps in complex samples and *H. influenzae* clinical isolates. Competitive activity-based screening using these probes identified novel IgA1P inhibitors, and structure-activity relationship guided optimization yielded a potent lead IgA1P inhibitor. This inhibitor preserves IgA1 at the bacterial surface, demonstrating effective inhibition of IgA1P activity in its native context and chemical disruption of an IgA1P-dependent immune-evasion mechanism. Together, these chemical tools establish a versatile platform for elucidating IgA1Ps role in *H. influenzae* colonization and virulence, and for validating IgA1Ps as diagnostic markers and antivirulence targets.

## Results and Discussion

Immunoglobulin A1 (IgA1) is the dominant antibody on mucosal surfaces of the respiratory tract and a key mediator of immune exclusion. It provides a first line of defense against bacterial colonization by coating bacteria, promoting agglutination, and limiting epithelial attachment[1-3]. Several important human bacterial pathogens, including *Haemophilus influenzae, Streptococcus pneumoniae* and pathogenic *Neisseria*, overcome this barrier by secreting IgA1 proteases (IgA1Ps)[4-7]. These enzymes selectively cleave IgA1, thereby disable its protective functions[8]. Since IgA1Ps directly compromise a central component of mucosal immunity, they are widely recognized as important virulence factors and attractive targets for anti-virulence strategies[9].

A defining feature of IgA1Ps is their extraordinary substrate specificity. These enzymes cleave strictly C-terminal to proline residues within the proline-rich IgA1 hinge region, thereby separating the antigen-binding Fab domains from the Fc region and functionally inactivating the antibody (**Figure 1A and B**)[3, 10-11]. As a result, IgA1P-producing bacteria become coated with IgA1-derived Fab fragments that lack Fc-mediated effector functions, enabling evasion of Fc receptor recognition and blocking access of intact antibodies of other isotypes[8, 12-13]. This highly specific reaction has emerged through convergent evolution in diverse bacterial pathogens, with mechanistically distinct serine-, metallo-, and cysteine-type proteases targeting the same IgA1 hinge motif[8]. Among these enzymes, serine-type IgA1Ps offer a particularly attractive chemical target profile. As chymotrypsin-like serine hydrolases bearing a canonical Ser–His–Asp catalytic triad[14], they are intrinsically amenable to covalent inhibition. At the same time, their strict requirement for proline at the P1 position is rare among human serine proteases and offers an unusual opportunity for selectivity[15].

**Figure 1.**
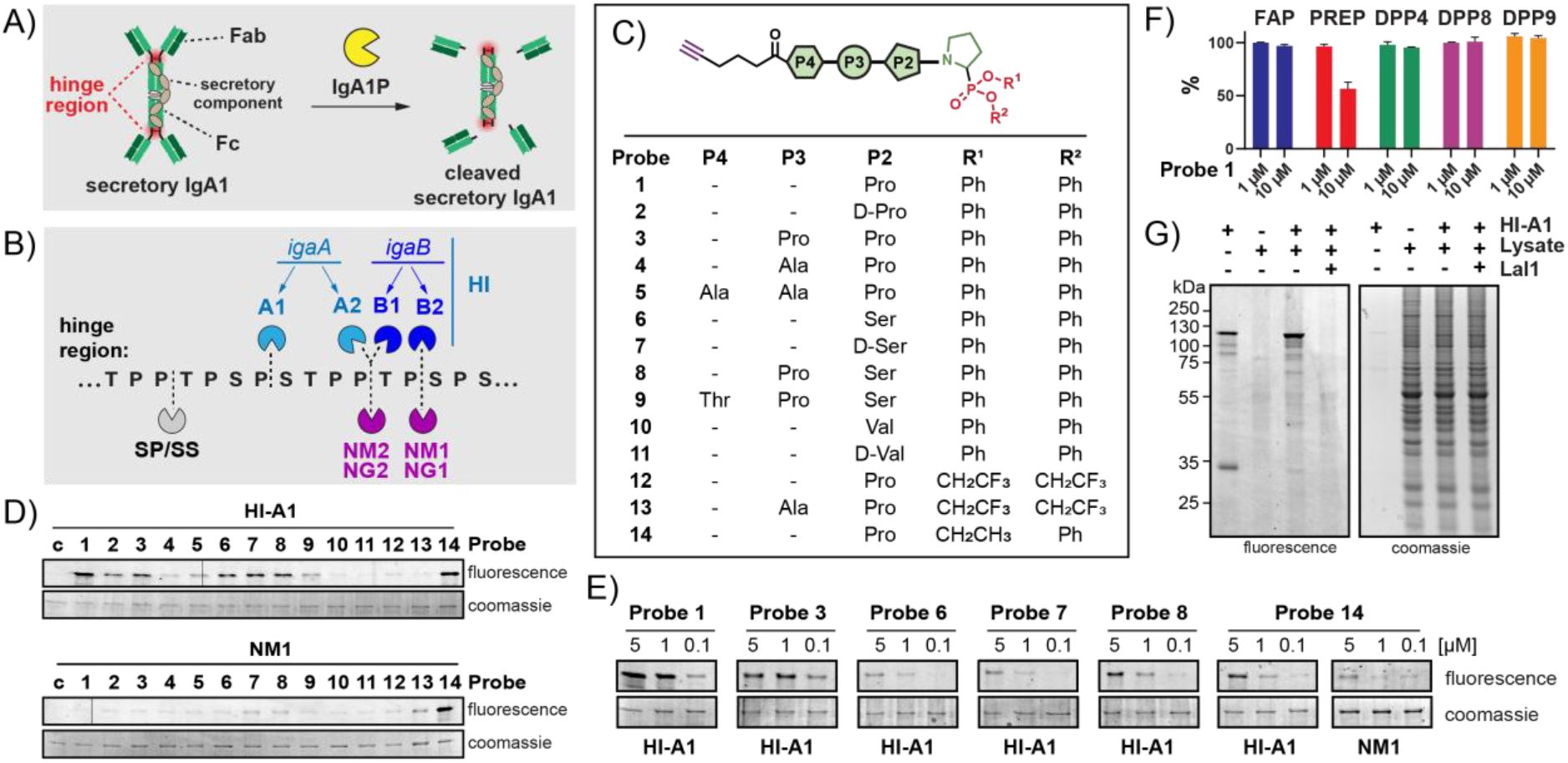
Labelling of recombinant serine-type IgA1Ps. A) Cleavage of IgA1 by IgA1P. B) Hinge region of IgA1 with location of peptide bonds cleaved by IgA1Ps. HI-A1, -A2,- B1 and -B2 is according to nomenclature by Murphy et al.[16, 20] (SS = *Streptococcus sanguinis* IgA1P, SP = *Streptococcus pneumoniae* IgA1P, HI = *Haemophilus influenzae* IgA1P, NM = *Neisseria meningitidis* IgA1P, NG = *Neisseria Gonorrhoeae* IgA1P). C) Probe library. D) Screening of probe library at 10 µM with HI-A1 and NM1 wild-type IgA1Ps (20 µg/mL). The vertical lines are imaging artifacts and do not represent gel splicing. E) Dose-down of probes with HI-A1 and NM1 wild-type IgA1Ps (20 µg/mL). F) Selectivity profiling of probe 1 (1 and 10 µM) against FAP, PREP, DPPs 4, -8, -9. Activity is reported as percentage of residual enzyme activity relative to the untreated control (v_i_/v_0_ × 100), where v_i_ and v_0_ denote the initial velocities in the presence and absence of probe, respectively. Data represent mean ± SD (n = 3). G) Selective and activity-dependent labeling of recombinant HI-A1 (20 µg/mL) by probe 1 (1 µM) in the background of mouse brain lysate (1 mg/mL). Lal1 = Lalistat 1.

In this context, IgA1Ps of *Haemophilus influenzae* (*H. influenzae*) are of high interest. *H. influenzae* is an exclusively human pathogen responsible for a broad spectrum of acute and chronic diseases, including meningitis, pneumonia, bacteremia, otitis media, and persistent respiratory infections in patients with chronic obstructive pulmonary disease (COPD)[16-17]. Although vaccination has dramatically reduced disease caused by *H. influenzae* type b strains (Hib), antibiotic-resistant non-typeable *H. influenzae* (NTHi) continues to represent a major clinical burden[18]. Pathogenic *H. influenzae* strains secrete serine-type IgA1Ps with substantially higher activity than their non-pathogenic relatives[19]. Virtually all clinical isolates encode the chromosomal *igaA* gene, while a substantial subset additionally carries the horizontally acquired *igaB* gene, giving rise to A-type and B-type IgA1Ps with distinct cleavage patterns (**Figure 1B**)[20-21]. These enzymes contribute differently to infection, with A-type IgA1Ps linked to epithelial invasion and B-type enzymes to intracellular survival[20, 22]. Despite their clear association with pathogenicity, how IgA1Ps function in native, infection-relevant environments remains poorly understood, in large part due to the lack of suitable experimental tools.

Progress in IgA1P research has been limited by two fundamental challenges: (i) the lack of animal models expressing human IgA1, and (ii) the absence of efficient chemical tools that enable detection or inhibition of IgA1Ps in complex samples. Several IgA1P inhibitors have been reported[23-28]; however, their limited potency or selectivity has largely restricted their use to purified enzyme systems. Likewise, existing approaches to monitor IgA1P activity rely on autoradiographic or Western blot-based detection of IgA1 cleavage, which are labor-intensive, expensive and poorly suited for broad application[29-31]. A FRET-substrate has also been described[32]; however, application of such substrates in complex mixtures can be challenging due to competing proteolytic activities. Consequently, direct, activity-dependent analysis of IgA1Ps *in situ* remains difficult.

Here, we address this challenge by developing activity-based probes (ABPs) that provide direct chemical access to *H. influenzae* IgA1Ps. These probes covalently label only the active form of the enzyme, enabling selective detection and visualization of IgA1Ps in tissue proteomes and clinical isolates. Using these probes in competitive profiling experiments, we identified and optimized IgA1P inhibitors that block IgA1 cleavage in native settings.

To develop ABPs for IgA1Ps, we exploited the conserved requirement for cleavage C-terminal to proline. We designed a focused probe library featuring a fixed P1 proline and systematic variation in P2–P4, guided by reported IgA1P inhibitors[25, 32], IgA1 hinge sequences, and selected non-natural residues to probe S-subsite preferences (**Figure 1C, Table S1**). Phosphonate warheads were chosen for covalent inhibition due to their high selectivity toward serine nucleophiles[33]. Specifically, diphenyl[34-36], phenyl/ethyl[37-38], and trifluoroethyl phosphonate[39] electrophiles were incorporated to span a range of phosphonate reactivities. Each probe was equipped with an N-terminal alkyne handle to allow in-gel analysis by rhodamine fluorophore installation via Cu-catalyzed azide–alkyne cycloaddition (CuAAC). Probes were synthesized using standard peptide coupling followed by late-stage phosphonate installation (**Scheme S1-3**). The prolyl-phosphonate warhead (H-Pro^P^(OPh)_2_) was prepared from freshly generated 1-pyrrolidine trimer and diphenylphosphite[40-41]. Alkyne-containing peptides were assembled via standard solid-phase peptide synthesis and subsequently coupled to H-Pro^P^(OPh)_2_ using HATU and DIPEA to afford **probes 1-11. Probes 12** and **13** were obtained from **probes 1** and **4**, respectively, through transesterification[42]. Lastly, **probe 14** was generated via transesterification followed by selective mono-dealkylation[43] and a final Steglich-type esterification.

For probe evaluation, we recombinantly expressed and purified the *H. influenzae* A1-type IgA1P (HI-A1), which represents the major and conserved IgA1P across clinical strains[14, 16]. The closely related serine-type homolog Neisseria meningitidis type 1 IgA1P (NM1) was included to benchmark probe performance across species and was likewise recombinantly produced and purified[44]. Individual probes were incubated with each recombinant enzyme for 2 hours, followed by CuAAC attachment of a rhodamine dye, SDS-polyacrylamide gel electrophoresis (SDS-PAGE), and in-gel fluorescence imaging. Seven of the fourteen probes labelled HI-A1 (**Figure 1D**). The strongest signals were obtained with **probe 1**, featuring a Pro– Pro recognition motif and a diphenyl phosphonate warhead, and **probe 14**, the corresponding mixed phenyl/ethyl analogue. In contrast, probes containing a P2 valine residue (**probes 10** and **11**) or a trifluoroethyl phosphonate electrophile (**probes 12** and **13**) showed no detectable labelling, indicating a sterically constrained S3 pocket and insufficient warhead reactivity. Strikingly, despite the high sequence similarity between HI-A1 and NM1, only **probe 14** labelled NM1.

Next, we performed dose–response experiments to rank probe performance. **Probe 1** emerged as the most potent candidate for HI-A1, producing the strongest signal and robust labelling at concentrations as low as 100 nM (**Figure 1E**). **Probe 1** also inhibited IgA1 cleavage in a concentration-dependent manner, confirming functional engagement of HI-A1 (**Figure S1**). None of the probes labeled a catalytically inactive HI-A1 S288A mutant, confirming active-site-directed covalent engagement (**Figure S2**). Although **probe 14** showed good binding in the initial screen against both NM1 and HI-A1, its performance was reduced in dose-response experiments for both enzymes, and it was excluded from further studies due to chemical instability (**Figure S3**). Based on potency and selectivity for HI-A1, **probe 1** was selected as the lead ABP for subsequent experiments.

Selectivity is a key parameter for assessing the applicability of activity-based probes. Therefore, the activity of **probe 1** was evaluated against recombinant human serine proteases with post-proline substrate specificity, including prolyl oligopeptidase (PREP), fibroblast activation protein (FAP), and the dipeptidyl peptidases (DPPs) DPP4, DPP8, and DPP9. In fluorogenic substrate assays, **probe 1** did not inhibit FAP or any DPP family member at concentrations of 1 or 10 µM, while PREP showed partial inhibition at a concentration of 10 µM (**Figure 1F**). However, PREP is localized intracellularly[45], whereas IgA1Ps act primarily in the extracellular space; therefore, this off-target interaction at high concentrations is unlikely to interfere with most IgA1P-directed applications. Consistent with this observation, **probe 1** also did not label the abundant chymotrypsin-like serine proteases human trypsin or chymotrypsin (**Figure S4**).

Probe selectivity was further evaluated in mouse lung and brain lysates, which represent complex host proteomes relevant to tissues commonly affected during *H. influenzae* disease. Both lysates contained abundant active serine hydrolases, as confirmed using a broad-spectrum serine hydrolase activity-based probe (**Figure S5**). When spiked with recombinant HI-A1, **probe 1** selectively labelled HI-A1 (**Figure 1G** and **S6A**). Labelling was abolished by preincubation with the IgA1P inhibitor Lalistat 1[26] (Lal1), confirming activity-dependent engagement. No additional bands were detected across either proteome, except for a faint signal observed in lung lysates at high probe concentrations (**Figure S6A**). This signal was eliminated by a PREP-selective inhibitor, identifying PREP as the only detectable off-target under these conditions (**Figure S6B**). PREP activity is higher in lung than brain lysates[46], explaining the weak lung-specific signal and indicating that potential PREP engagement in lung tissue could be mitigated by co-application of a PREP inhibitor in ex vivo or in vivo settings, if required.

Having established probe selectivity across purified IgA1Ps and complex host proteomes, we next investigated whether **probe 1** can directly report endogenous IgA1P activity in native bacterial samples. To this end, we assembled a compact panel of diverse *H. influenzae* clinical isolates, which provide direct access to secreted IgA1Ps in their physiological context (**Figure 2A, Table S2**). As a functional reference, we first assessed cleavage of IgA1 across the isolate panel. Five of six isolates showed clear IgA1P activity, enabling classification into A- and B-type enzymes (**Figure 2B**). Isolates #1, #2, and #5 displayed A-type cleavage, matching recombinant HI-A1, whereas isolates #3 and #4 showed B-type cleavage, matching recombinant NM1, a Neisseria-derived progenitor of B-type IgA1Ps in *H. influenzae*[47]. For direct detection, a small set of fluorescent derivatives of **probe 1** was synthesized (**Scheme S4**), and sulfo-Cy5–labeled variant (**probe 15**) was selected for further studies (**Figures 2C** and **S7**). Strikingly, **probe 15** was able to label native IgA1Ps directly in culture supernatants, yielding a dominant band at ∼120 kDa in all IgA1-cleaving isolates, while the inactive isolate #6 showed no signal (**Figure 2D**). Labelling was activity-dependent, as it was suppressed by the IgA1P inhibitor Lal1. Residual labeling in isolate #3 is consistent with the reduced sensitivity of native B-type IgA1Ps to Lal1[26]. A probe concentration series revealed strain-dependent differences in native IgA1P activity across isolates (**Figure 2E**). Isolate #3 showed the highest activity, with labelling at 100 nM probe concentration, whereas isolates #1 and #4 required 1 µM and isolates #2 and #5 required 5 µM for detection. These data demonstrate that native IgA1P output can be measured with high sensitivity using a direct, activity-based readout. Although probe design was guided by the A-type IgA1P HI-A1, **probe 15** also labelled native B-type IgA1Ps. This indicates that the probe engages conserved catalytic features shared across *H. influenzae* IgA1Ps. As the presence of igaB has been linked to elevated protease activity and is regarded as an independent virulence determinant[21], the ability to capture B-type IgA1Ps is particularly important for functional studies in native samples.

**Figure 2.**
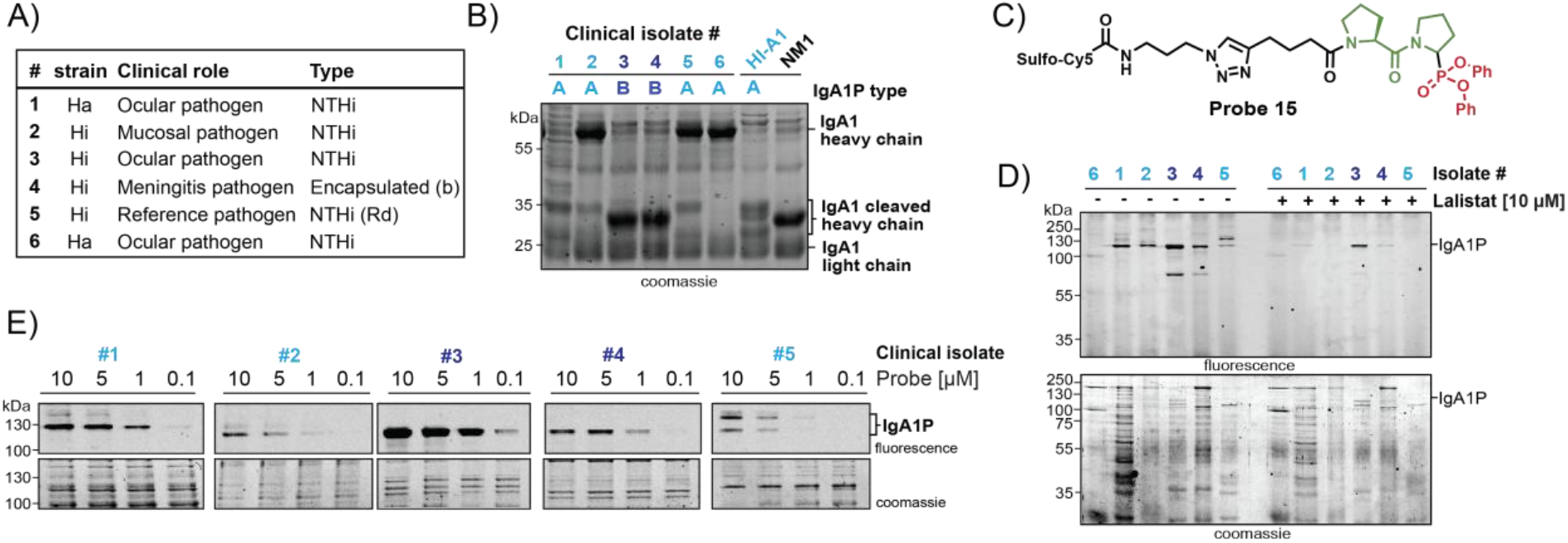
Activity-based profiling of endogenous IgA1Ps in *H. influenzae* clinical isolates. A) Panel of *H. influenzae* clinical isolates used in this study, indicating isolate number and anatomical origin. Hi = *H. influenzae*; Ha = *H. influenzae* biotype aegyptius B) IgA1 hinge-cleavage patterns generated by culture supernatants of the indicated isolates, revealing functional IgA1P activity. C) Chemical structure of fluorescent activity-based probe 15. D) Direct detection of endogenous IgA1Ps in unprocessed culture supernatants of H. influenzae clinical isolates by probe 15 (25 µM). E) Probe 15 concentration series in selected clinical isolates, revealing isolate-dependent differences in IgA1P activity and enabling activity-based ranking of IgA1P secretion.

A central advantage of activity-based probes is that they enable both detection and inhibition of enzyme activity. We therefore explored probe-guided strategies to identify IgA1P inhibitors. Despite their relevance as anti-virulence targets, IgA1Ps have proven difficult to inhibit: even high-throughput screening efforts have produced only moderately potent inhibitors, likely due to their highly specific substrate recognition[23, 26]. To address this challenge, we leveraged an in-house library comprising a focused set of 147 compounds designed for post-proline-cleaving proteases (**Figures S8–S15**). This library predominantly comprised of non-covalent and covalent peptidic inhibitors bearing proline or proline analogues in the P1 position, incorporating either nitrile-or phosphonate-based warheads. **Probe 15** was applied in competitive screening format as a direct readout of active-site engagement on recombinant HI-A1 (**Figures S16A**). The screening yielded only a limited number of hits with micromolar potency, further emphasizing the challenging nature of IgA1Ps as drug targets (**Figures S16B** and **C**). The most potent hit, **UAMC-936**, was resynthesized (**UAMC-936-1**) and showed comparable inhibitory profile on HI-A1 with complete inhibition at 50 µM and partial inhibition at 10 µM (**Figure 3B** and **S16C**). Notably, UAMC-936 is a close structural analogue of the most potent ABP (**probe 1**), thereby validating the probe design and underscoring the highly restricted substrate specificity of IgA1Ps.

**Figure 3.**
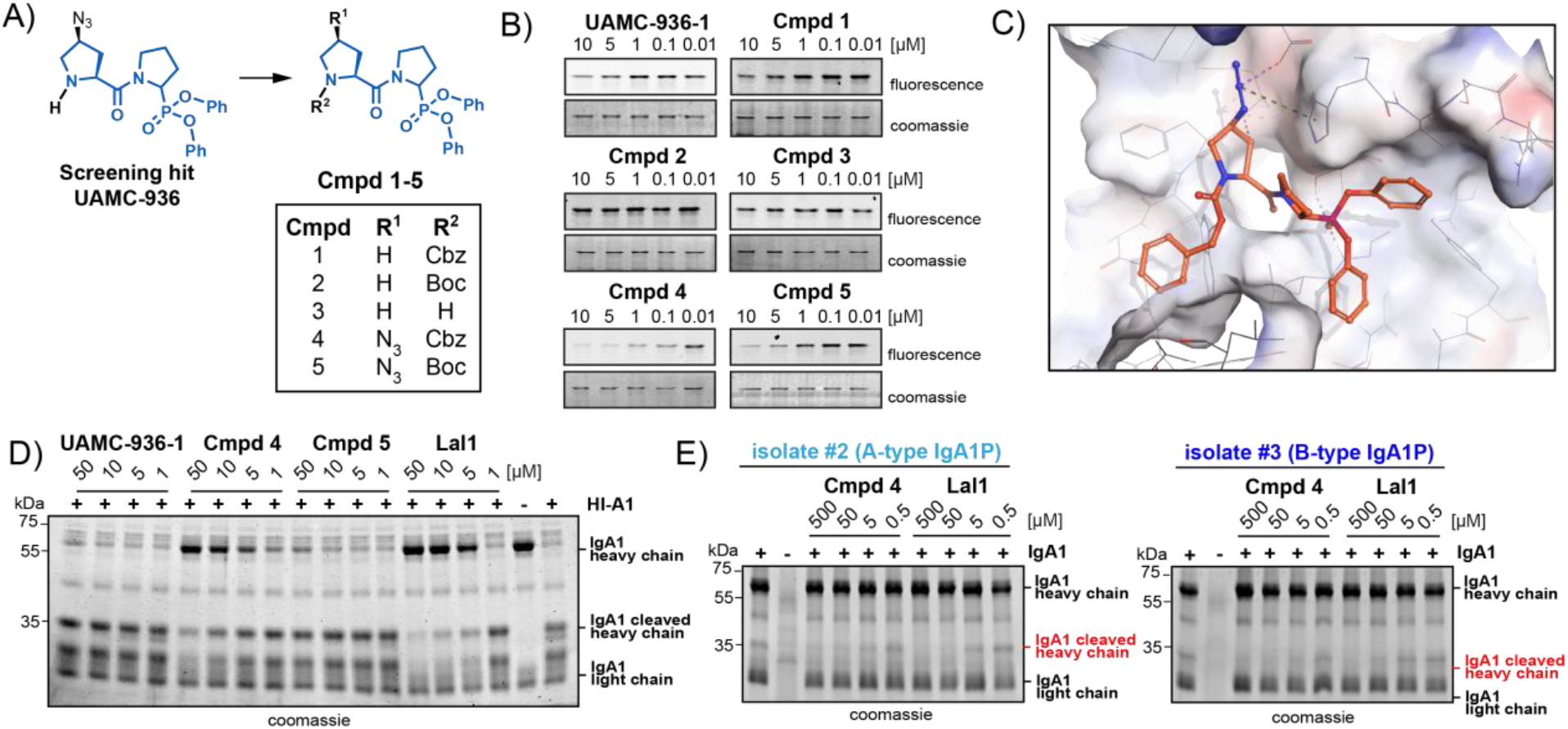
Identification of IgA1P inhibitors. A) Synthesized focused inhibitor library. B) Concentration-dependent inhibition of recombinant HI-A1 (20 µg/mL) by the resynthesized screening hit UAMC-936-1 and inhibitors, assessed by competition with probe 15. C) Molecular docking of compound 4 within the active site of HI-IgA1P. D) Inhibition of IgA1 cleavage by recombinant HI-A1 (20 µg/mL). E) Inhibition of IgA1 cleavage in *H. influenzae* culture supernatants. Cmpd = Compound; Lal1 = Lalistat 1; Cbz = benzyloxycarbonyl; Boc = tert-butoxycarbonyl.

Next, we sought to improve inhibitor affinity towards HI-A1, five analogues of **UAMC-936** were synthesized (**Figure 3A**). As IgA1Ps act as endopeptidases, modifications were introduced at the P2-N position. Cbz- and Boc-protected variants, as well as free amine analogues, were prepared with and without an azide functionality (**compounds 1–5**). In dose-down labelling assays with recombinant HI-A1 (**Figure 3B**), compound 4 (Cbz-protected analogue of UAMC-936) emerged as the most potent inhibitor, showing significant inhibition at concentrations as low as 100 nM. The azide functionality proved critical for enhanced potency, as evidenced by the comparison of compounds 1 and 4, whereas Boc-protected and free amine variants (**compounds 2, 3**, and **5**) showed markedly reduced inhibition. These findings indicate a preference for neutral, bulky substituent at the P2-N position, while positively charged amines are poorly accommodated. Docking of compound 4 within the HI-IgA1P active site is consistent with these observations (**Figure 3C** and **S17**). Functional inhibition was confirmed in an HI-A1 IgA1 cleavage assay (**Figure 3D**). Compound 4 strongly suppressed IgA1 cleavage at concentrations as low as 5 µM, showing potency comparable to the reference inhibitor Lal1 (IC_50_ = 4.7 µM for HI-A1[26]).

To determine whether native IgA1Ps of different types are also targeted, **compound 4** was evaluated in *H. influenzae* clinical isolates expressing either A-type (isolate #2) or B-type (isolate #3) enzymes. Consistent with IgA1 cleavage assays using recombinant HI-A1, A-type IgA1P activity was significantly inhibited by both **compound 4** and Lal1 at 5 µM, as indicated by the disappearance of the cleaved IgA1 heavy-chain fragment (**Figure 3E**). In contrast, B-type IgA1P activity was completely suppressed by compound 4 at 5 µM, whereas Lal1 required a 100-fold higher concentration (500 µM) to achieve full inhibition. These data demonstrate that **compound 4** is markedly more potent against native B-type IgA1Ps.

Having identified **compound 4** as a potent inhibitor of *H. influenzae* IgA1Ps, we next tested whether this inhibition produces a functional effect on immune evasion in intact bacteria. IgA1Ps promote immune evasion by cleaving surface-bound IgA1, thereby preventing bacterial agglutination and mucosal immune protection[8, 10, 48]. Blocking this activity should therefore preserve surface-bound IgA1. To test this, we selected a *H. influenzae* clinical isolate with high IgA1P activity (isolate #3, B-type). Whole bacterial cultures were pre-incubated with increasing concentrations of **compound 4** and subsequently exposed to human secretory IgA1 to allow antibody coating (**Figure 4A**). Cells were then stained with anti-IgA1-FITC and analyzed by flow cytometry. Untreated bacteria showed low fluorescence, consistent with efficient IgA1P-mediated removal of surface-bound IgA1 (**Figure 4B**). In contrast, treatment with **compound 4** resulted in a clear, concentration-dependent increase in surface-associated IgA1. Percentage of FITC-positive bacteria increased by 26% at 10 µM and by 44% at 100 µM relative to untreated cells. These data demonstrate that chemical inhibition of IgA1Ps restores IgA1 on the bacterial surface under conditions of high protease activity. Importantly, **compound 4** showed no antibacterial activity up to 250 µg mL^-1^ (**Figure S18**), supporting its potential as an anti-virulence lead and a chemical tool to study IgA1P-dependent immune evasion in native bacterial contexts.

**Figure 4.**
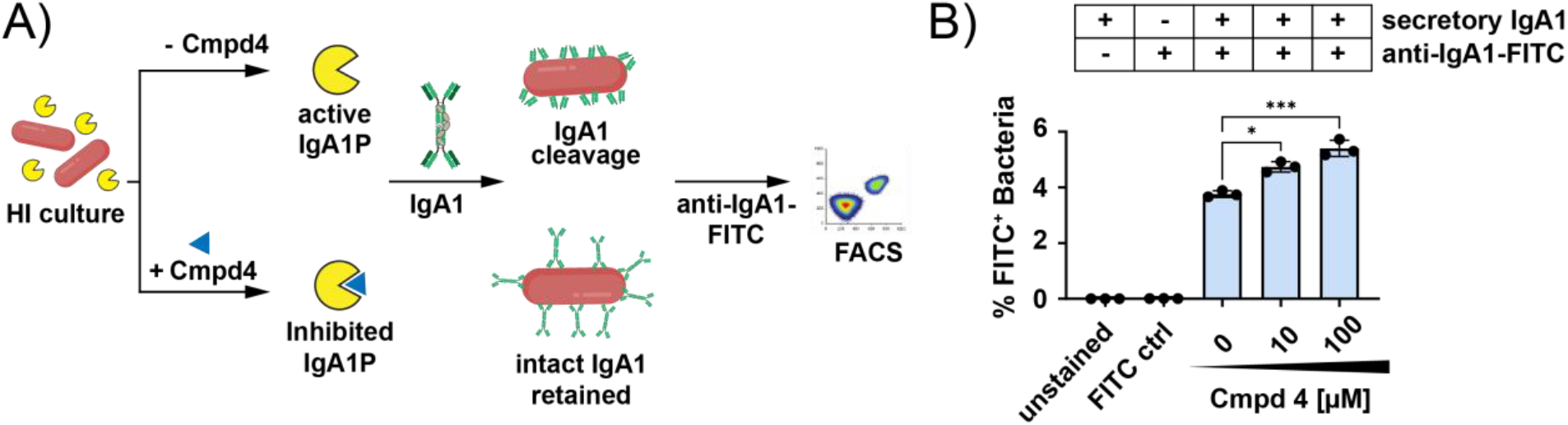
Modulation of IgA1P-dependent virulence-associated activities. A) Clinical isolate #3 cultures were pre-incubated with Cmpd 4 or vehicle and subsequently treated with secretory IgA1, which binds to the bacterial surface. Remaining surface-bound IgA1 was detected using an anti-IgA1–FITC antibody and quantified by fluorescence-activated cell sorting (FACS). B) %FITC-positive bacteria quantified by FACS, following treatment of isolate #3 with Cmpd 4 (10 and 100 µM), alongside unstained, FITC ctrl, and DMSO negative control samples. Data represent mean ± SD (n = 3 technical replicates). Statistical analysis was performed using ordinary one-way ANOVA with Dunnett’s post hoc test versus the DMSO control. *p < 0.05; ***p < 0.001

In summary, this work establishes the first activity-based probes that enable selective detection of IgA1Ps. By covalently targeting the active site of the major *Haemophilus influenzae* IgA1P, these probes provide access to IgA1P activity in complex proteomes and clinical isolates. The probe platform further enabled competitive profiling that guided the optimization of a small-molecule inhibitor that blocks IgA1 cleavage on live bacteria without affecting growth, providing a chemical strategy to interfere with IgA1P-mediated immune evasion. The modular probe scaffold further offers the opportunity to incorporating alternative tags for enrichment-based workflows, quantitative proteomics, and comparative analyses of IgA1P activity across strains, infection models, and patient-derived samples[49-51]. Moreover, selective labelling of active IgA1Ps may facilitate investigation of emerging host substrates beyond IgA1[44, 52-53]. Subtle warhead modification can tune probe selectivity across IgA1P classes, as shown for NM1, highlighting the potential to extend this platform to IgA1Ps from other clinically relevant pathogens. Together, these probes and inhibitors provide powerful chemical tools to investigate the biological roles of this virulence-associated protease family while opening new opportunities for diagnostic development and antivirulence drug discovery.

## Acknowledgements

We thank Prof. Koen Augustyns for his generous support. We are grateful to Prof. Mogens Kilian (Aarhus University) for selecting and donating the clinical strains, analyzing the genome of sequenced strains for IgA1P gene presence, and for critical reading of the manuscript. We thank Prof. Todd Holyoak (University of Waterloo) for providing the wild-type and mutant HI-A1 plasmids. We acknowledge Julie Darrigues and Vera Dermesrobian of the KU Leuven Flow and Mass Cytometry Core Facility (FACS Core KU Leuven) for expert assistance and guidance with flow cytometry data analysis, and Prof. Greetje Vande Velde and Lauren Michiels (KU Leuven) for the donation of mouse tissues. We thank Valeria Parravicini for technical support. This work was supported by the Bijzonder onderzoeksfonds (BOF), University Antwerp. V. V. was supported by a Fonds Wetenschappelijk Onderzoek (FWO) postdoctoral fellowship (12A5G25N), P. T. and S. P. by FWO Ph.D. fellowships (11A9V26N and 11PCL24N), and A. F. by a Marie Skłodowska-Curie Actions Postdoctoral Fellowship (MSCA-2023-PF-01-101155555).

