## Supplementary Information for "Chemical tools to Detect and Inhibit IgA1 Proteases in *Haemophilus influenzae*"

[c] Flow Cytometry and Cell Sorting Facility
University of Antwerp, Universiteitsplein 1, 2610 Antwerp, Belgium

[d] Department of Microbiology, Tumor and Cell Biology, Clinical Microbiology
Karolinska Institute, 171 77 Stockholm, Sweden

[e] Clinical Microbiology
Karolinska University Hospital, 171 76 Stockholm, Sweden

[f] Singapore Centre on Environmental Life Sciences Engineering
Nanyang Technological University, 637551, Singapore

+ equal contribution

### Table of Contents


| Section | Page |
|---|---|
| 1. Supplementary Tables | S3 |
| 2. Supplementary Figures | S5 |
| 3. Biochemical, Microbiological and Computational Methods | S19 |
| 4. Synthetic Schemes | S24 |
| 5. Synthesis of Probes and Compounds | S25 |
| 6. LC-MS Chromatograms | S37 |
| 7. NMR Spectra | S50 |
| 8. Analytical Data for Unpublished Compounds from the In-House Library | S54 |

### 1. Supplementary Tables

**Table S1. Peptide recognition motifs evaluated for IgA1P-directed activity-based probe (ABP) development.** Peptides were selected based on sequence features of the IgA1 hinge region, incorporation of non-natural amino acids, prior inhibitor reports, and selected residues outside the hinge region.

| Probe | Peptide sequence (N→ C) | Design rationale |
|---|---|---|
| 1 | Pro-Pro | Based on IgA1 hinge motif (type A2 and B1); expected IgA1P recognition[25] |
| 2 | (D)-Pro-Pro | D-Pro substitution to probe stereochemical tolerance |
| 3 | Pro-Pro-Pro | Extended hinge region mimic |
| 4 | Ala-Pro-Pro | Based on reported IgA1P inhibitors for *H. influenzae* and *N. gonorrhoeae* [24] |
| 5 | Ala-Ala-Pro-Pro | Based on reported IgA1P inhibitors for *H. influenzae* and *N. gonorrhoeae* [24] |
| 6 | Ser-Pro | Based on IgA1 hinge motif (type A1); expected IgA1P recognition[25] |
| 7 | (D)-Ser-Pro | D-Ser substitution to probe stereochemical tolerance |
| 8 | Pro-Ser-Pro | Based on IgA1 hinge motif (type A1); expected IgA1P recognition[25] |
| 9 | Thr-Pro-Ser-Pro | Based on IgA1 hinge motif (type A1); expected IgA1P recognition[25] |
| 10 | Val-Pro | Probing IgA1P recognition outside the hinge region |
| 11 | (D)-Val-Pro | Probing IgA1P recognition outside the hinge region |

***Note:*** The **Thr-Pro** motif (type B2) was not included in the recognition motif evaluation, as coupling between Alk-Thr(tBu)-OH and Pro-DPP consistently resulted in dehydration to form an alkene-containing by-product.

**Table S2. Diverse panel of *H. influenzae* clinical isolates differing in biotype, serotype, anatomical origin, and IgA1P allele.** The panel includes two isolates classified as *H. influenzae* biotype *aegyptius*, a divergent member of the *H. influenzae* species complex, to test probe performance across clinically relevant backgrounds.

| clinical isolate No. | Strain ID | Genus | Species | Alternative ID | Sero-type | Bio-type | Clinical origin | IgA1P genome location | Source |
|---|---|---|---|---|---|---|---|---|---|
| #1 | HK367 | *Haemophilus* | *aegyptius* | NCTC8502 | | III | Conjunctiva | *igaA* | NCTC |
| #2 | HK389 (type strain) | *Haemophilus* | *influenzae* | NCTC8143 | | II | Respiratory tract | *igaA* | NCTC |
| #3 | HK286 | *Haemophilus* | *influenzae* | Mordhost AB1890 | | III | Conjunctivitis | | SSI |
| #4 | HK177 | *Haemophilus* | *influenzae* | Lautrop AB1881 | b | I | Meningitis (Denmark) | | SSI |
| #5 | HK398 | *Haemophilus* | *influenzae* | CIP5424 Rd | Rd | IV | Reference strain | *igaA* | NCTC |
| #6 | HK369 | *Haemophilus* | *aegyptius* | NCTC8135 | | III | Conjunctiva | *igaA* | NCTC |

**Table S3**. **IgA1P expression constructs, vector backbones, and sources.**

| Plasmid ID | Properties | Vector | Provider |
|---|---|---|---|
| pET-24bH.inf IgA1P wt[14] | Protease domain of HI-A1 from *H. influenzae* strain Rd | pET-24a (+) | Todd Holyoak, University of Kansas Medical Centre, USA. |
| pET-24bH.inf IgA1P S288A mt [14] | S288A mutant of HI-A1 from *H. influenzae* strain Rd | pET-24a (+) | Todd Holyoak, University of Kansas Medical Centre, USA. |
| pNIC-CTHF NM IgA1P T1 wt[44] | Protease domain of NM1 from *Neisseria meningitidis* strain MC58 | pNIC-CTHF | Edmund Loh, Karolinska Institute, Sweden. |

#### 2. Supplementary Figures

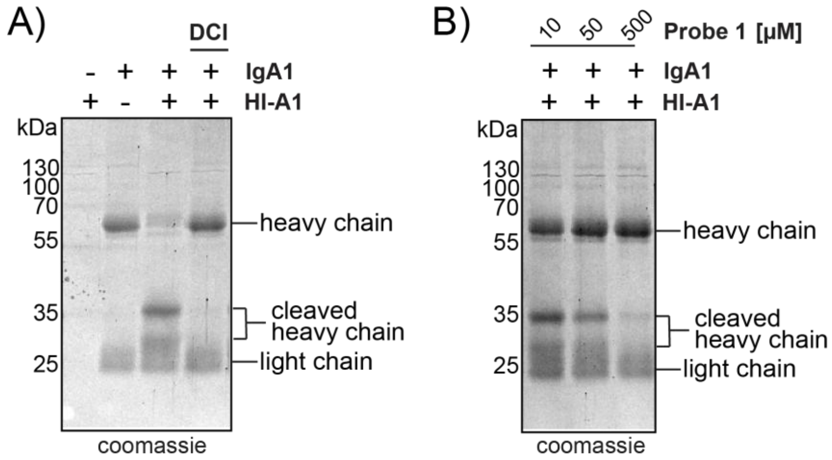


**Figure S1. Inhibition of IgA1P-mediated IgA1 cleavage.** (A) The serine protease inhibitor 3,4-dichloroisocoumarin (DCI, 10 µM) blocks IgA1 heavy-chain cleavage by HI-A1 wt, confirming serine protease–dependent activity. (B) Probe 1 inhibits IgA1 cleavage in a concentration-dependent manner (≥10 µM).

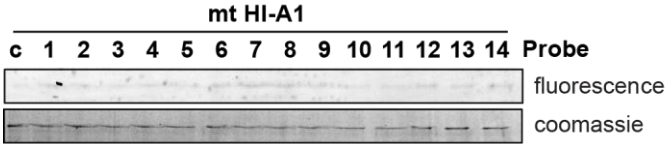


**Figure S2. Lack of probe labelling of the inactive HI-A1 S288A mutant.** None of the probes in the library (10 µM) labelled the HI-A1 S288A mutant, in which the catalytic Ser288 is replaced by alanine.

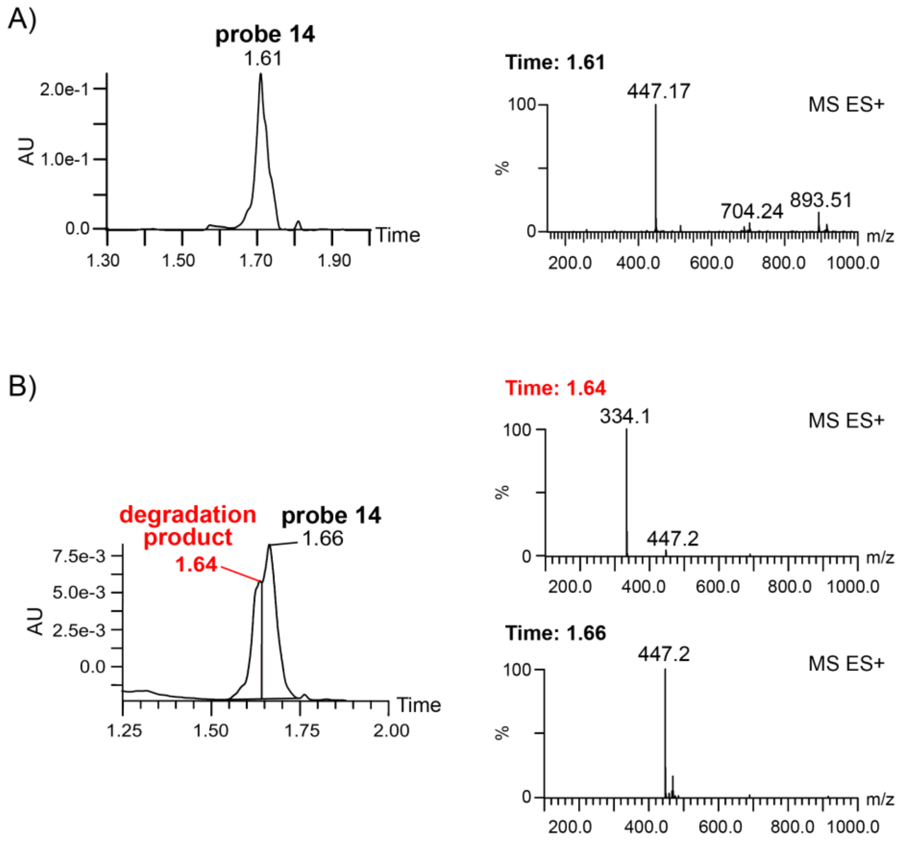


**Figure S3. Degradation of probe 14.** A) UPLC–MS analysis of 99% pure probe 14. B) UPLC–MS analysis of probe 14 after 24 h in DMSO. The solid material and the DMSO stock showed identical degradation profiles after 24 h; therefore, only the DMSO stock is shown. The observed instability explains the weak potency of probe 14 in Figure 1E.

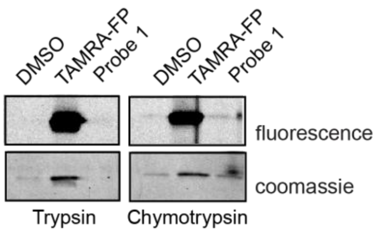


**Figure S4. Off-target reactivity of Probe 1 against serine proteases**. Human trypsin and chymotrypsin were incubated with Probe 1 (10 µM) or TAMRA–fluorophosphonate (FP–TAMRA, 10 µM). FP–TAMRA labelled both enzymes, whereas Probe 1 showed no detectable labelling. Reduced Coomassie signal in the Probe 1 and DMSO samples reflects protease autocleavage, while the stronger band in the FP–TAMRA sample indicates inhibition.

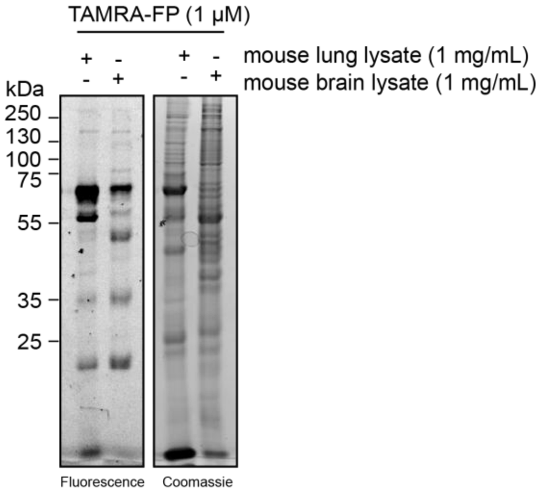


**Figure S5. Serine hydrolase activity in mouse tissue lysates.** Mouse brain and lung lysates were incubated with the broad-spectrum serine hydrolase probe FP–TAMRA. Fluorescence imaging reveals active hydrolases in the lysates, while Coomassie staining shows total protein loading.

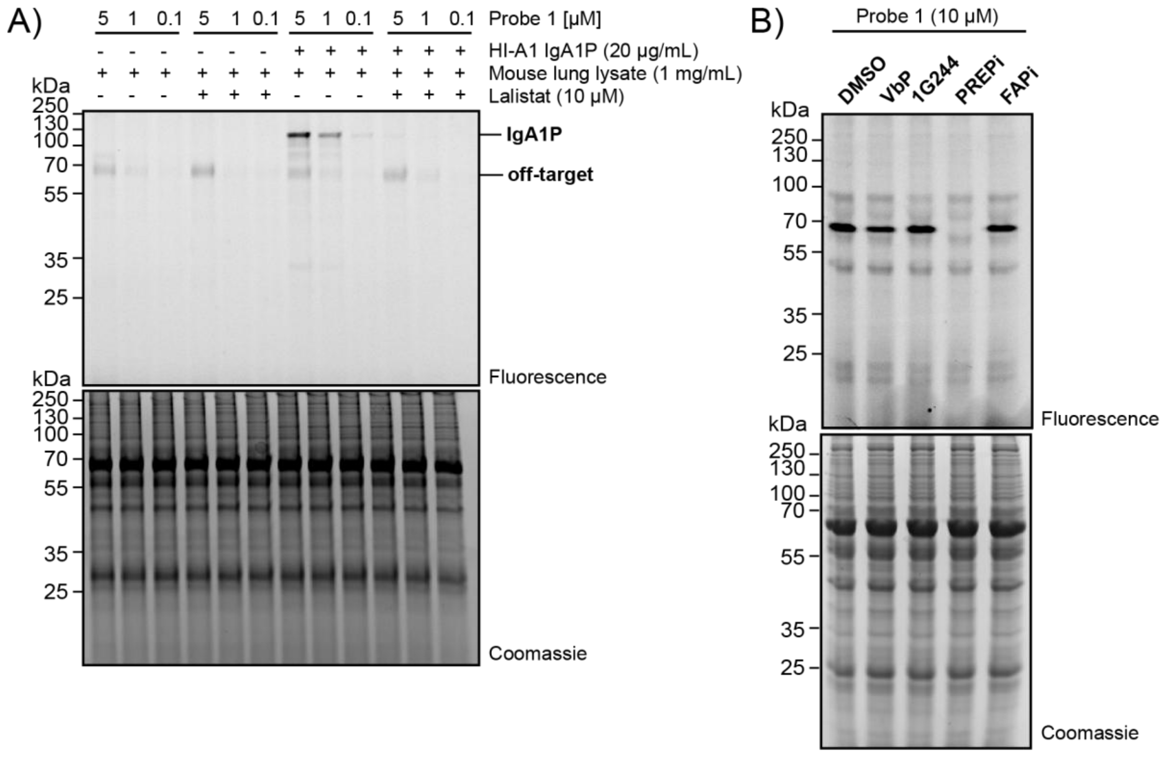


**Figure S6. Identification of a minor off-target of probe 1 in mouse lung lysates.** A) Labelling of recombinant HI-A1 by probe 1 in the presence of mouse lung lysate resulted in a dominant HI-A1 band and a single minor off-target signal at high probe concentrations. B) Preincubation of lung lysates with the pan-DPP protease inhibitor VbP (Val-boroPro)[54], the DPP8/9 selective inhibitor 1G244 (UAMC-618[55], the PREP-selective inhibitor PREPi (KYP2047)[56], or a FAP-selective inhibitor (FAPi, UAMC-1110)[57], revealed that only PREPi abolished the off-target signal, identifying the off-target as PREP.

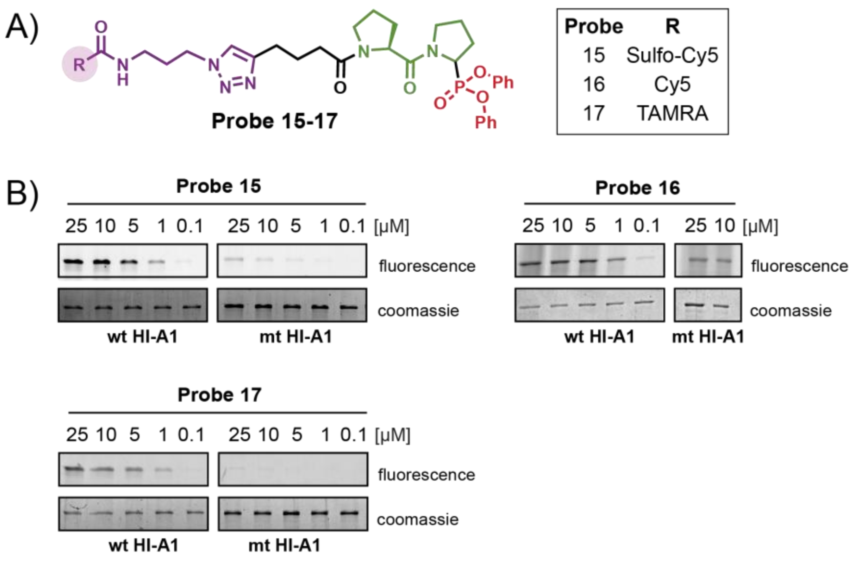


**Figure S7. Development of fluorescent probes.** A) Structure of fluorescent probes 15-17. B) Incubation of wt HI-A1 or mt HI-A1 with a concentration series of probes 15–17 showed similar labelling profiles. Among them, probe 15 gave the strongest signal at 5 and 1 µM, and was therefore selected for fluorescent probe labelling experiments.

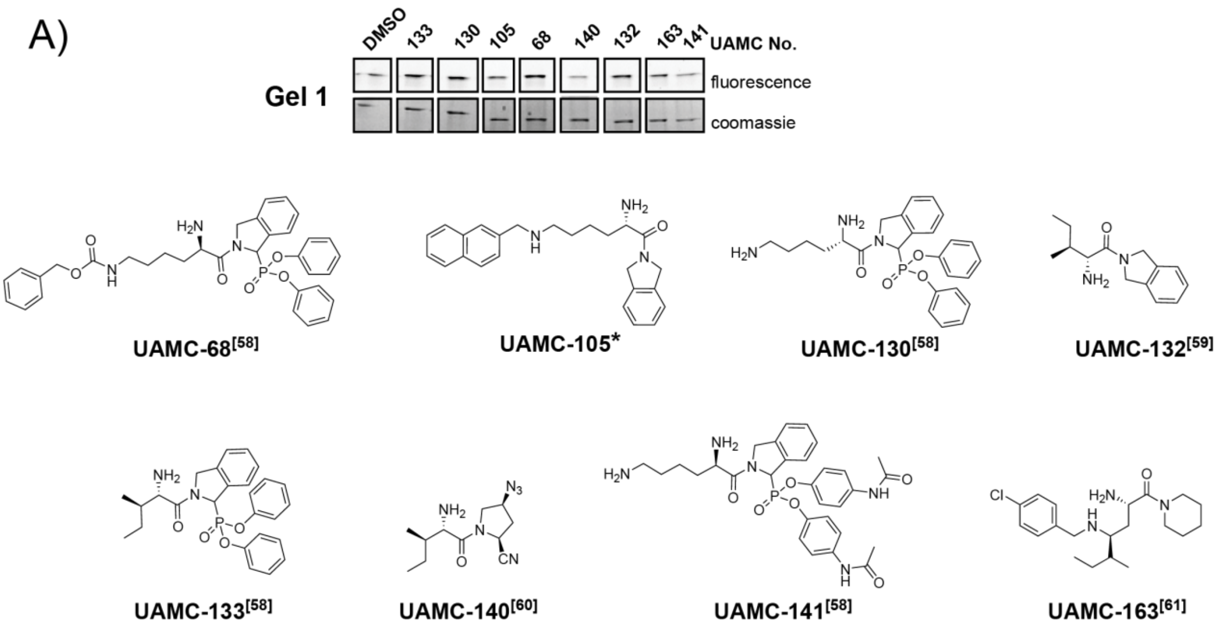


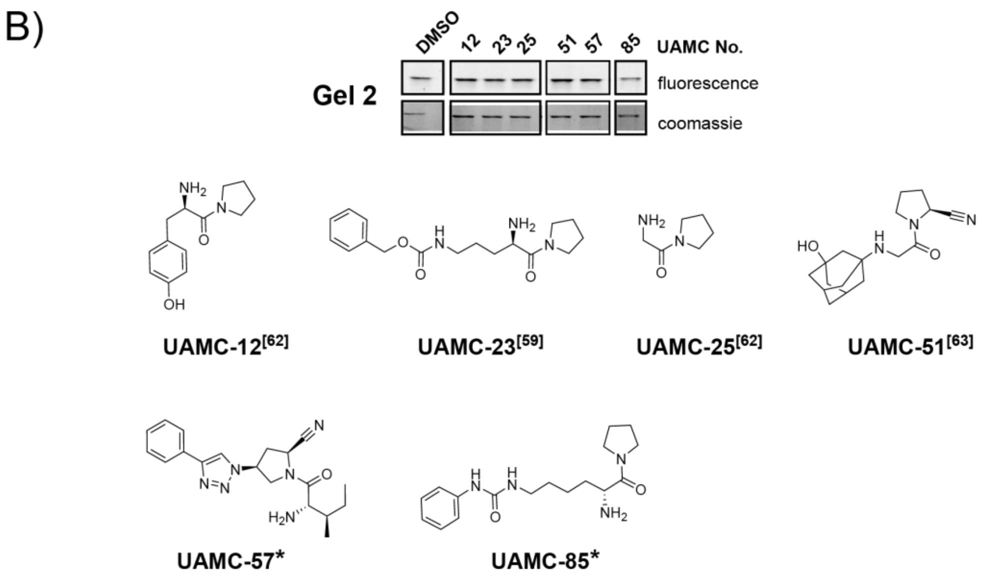


**Figure S8. Structures of the focused in-house compound library and initial screening against recombinant HI-A1.** Compounds (50 µM) were screened using Probe 15 (5 µM). Due to partial degradation of aged library members, compound purity was determined after screening, and only lanes corresponding to compounds with ≥70% purity are shown. Active compounds were defined as those causing >50% reduction of the probe-derived fluorescence signal relative to the DMSO control. No active compounds were identified in these gels. Unpublished library members (*) are included for completeness; their HRMS and HPLC characterization data are provided in Section 8. 68, 130, 133, 141[58], 132[59], 140[60], 163[61], 12[62], 23[59], 25[62], 51[63]

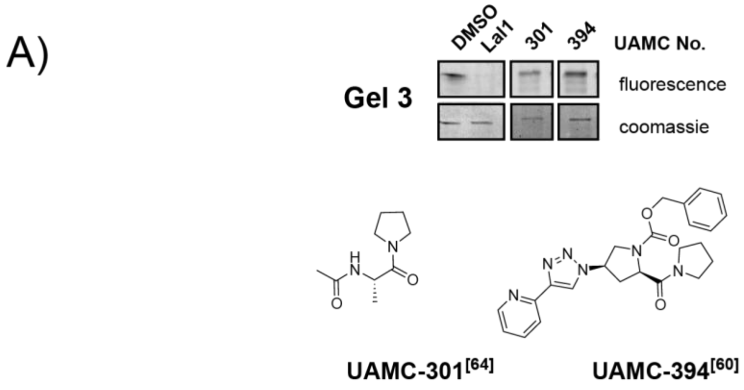


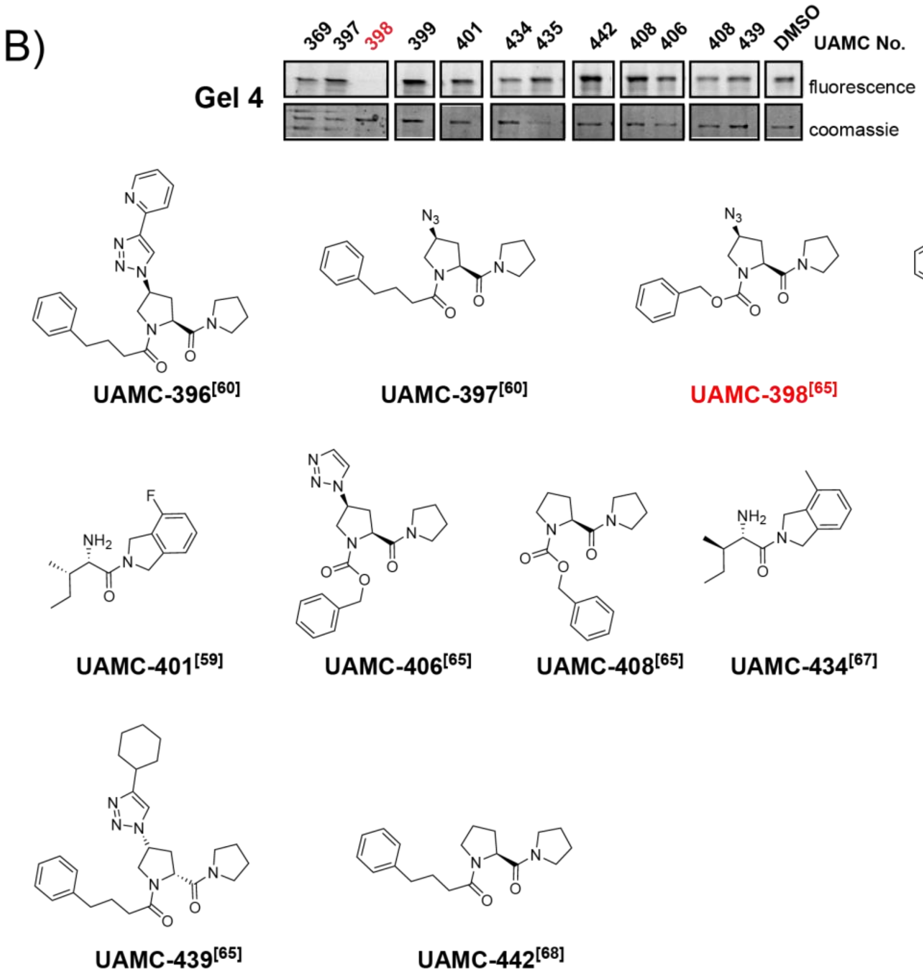


**Figure S9. Structures of the focused in-house compound library and initial screening against recombinant HI-A1.** Compounds (50 µM) were screened using Probe 15 (5 µM). Due to partial degradation of aged library members, compound purity was determined after screening, and only lanes corresponding to compounds with ≥70% purity are shown. Active compounds were defined as those causing >50% reduction of the probe-derived fluorescence signal relative to the DMSO control and are highlighted in red. Unpublished library members (*) are included for completeness; their HRMS and HPLC characterization data are provided in Section 8. Lal1 = Lalistat 1. 301[64], 394[60], 396, 397, [60], 398[65], 399[66], 401[59], 406[65], 408[65], 434[67], 439[65], 442[68]

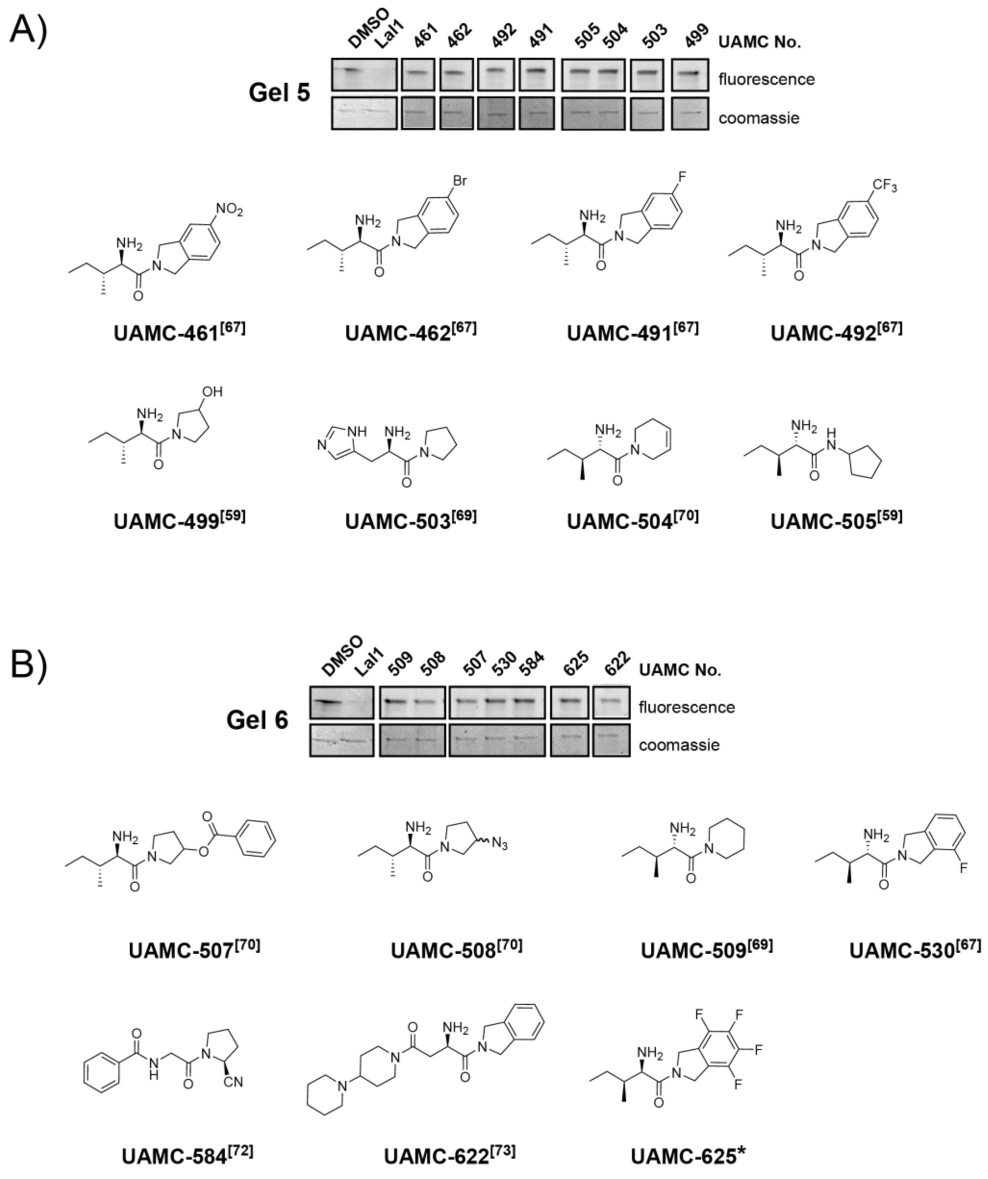


**Figure S10. Structures of the focused in-house compound library and initial screening against recombinant HI-A1.** Compounds (50 µM) were screened using Probe 15 (5 µM). Due to partial degradation of aged library members, compound purity was determined after screening, and only lanes corresponding to compounds with ≥70% purity are shown. Active compounds were defined as those causing >50% reduction of the probe-derived fluorescence signal relative to the DMSO control. No active compounds were identified in these gels. Unpublished library members (*) are included for completeness; their HRMS and HPLC characterization data are provided in Section 8. Lal1 = Lalistat 1.
461, 462, 491, 492[67], 499[59], 503[69], 504[70], 505[59], 25[71], 507, 508[70], 509[69], 530[67], 584[72], 622[73].

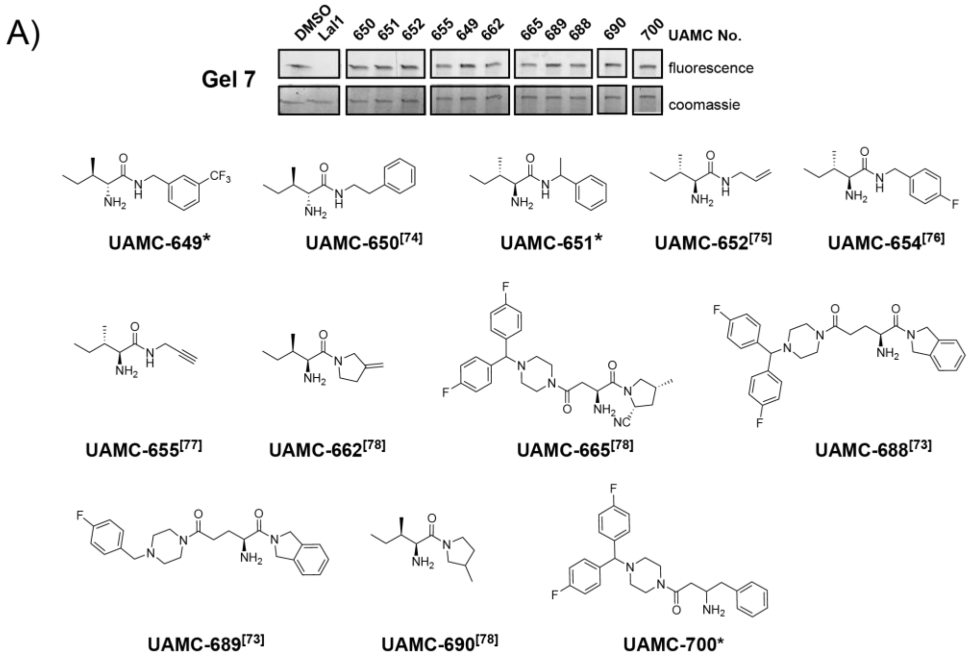


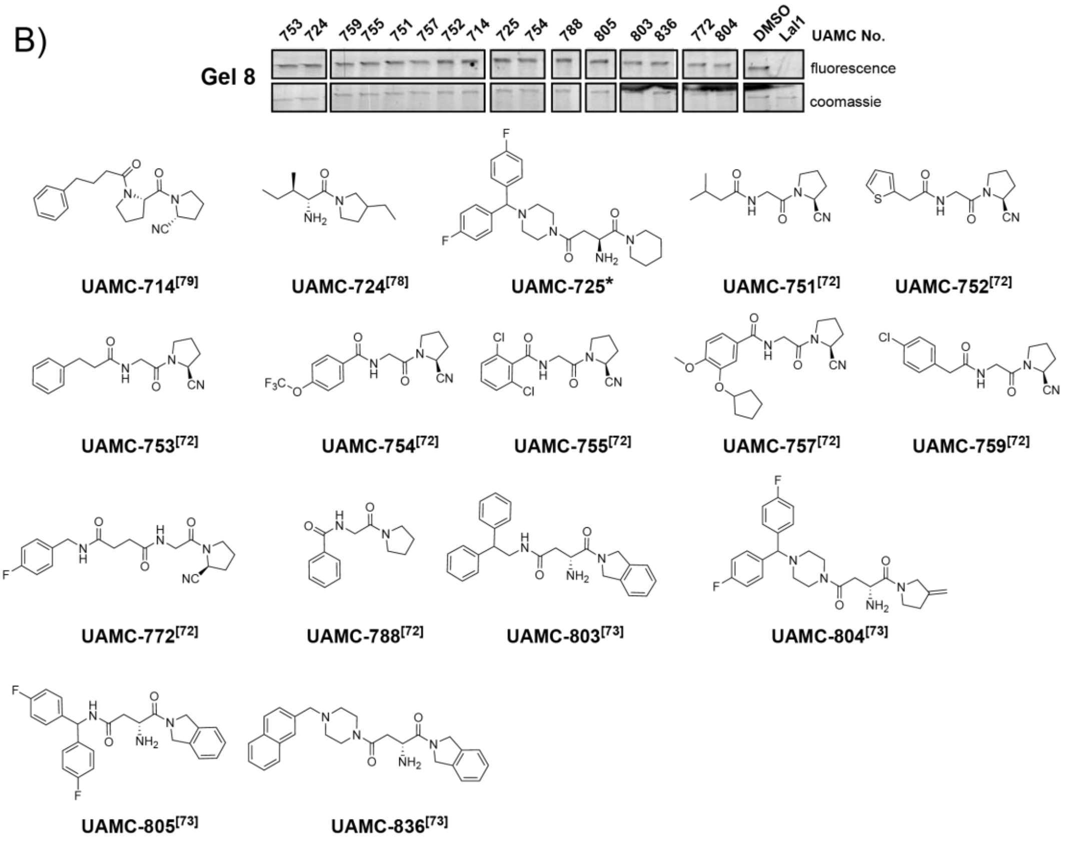


**Figure S11. Structures of the focused in-house compound library and initial screening against recombinant HI-A1.** Compounds (50 µM) were screened using Probe 15 (5 µM). Due to partial degradation of aged library members, compound purity was determined after screening, and only lanes corresponding to compounds with ≥70% purity are shown. Active compounds were defined as those causing >50% reduction of the probe-derived fluorescence signal relative to the DMSO control. No active compounds were identified in these gels. Unpublished library members (*) are included for completeness; their HRMS and HPLC characterization data are provided in Section 8. Lal1 = Lalistat 1. 650[74], 652[75], 654[76], 655[77], 662, 665, 690, 724[78], 688, 689[73], 714[79], 751, 752, 753, 754, 755, 757, 759, 772, 788[72], 803, 804, 805, 836[73]

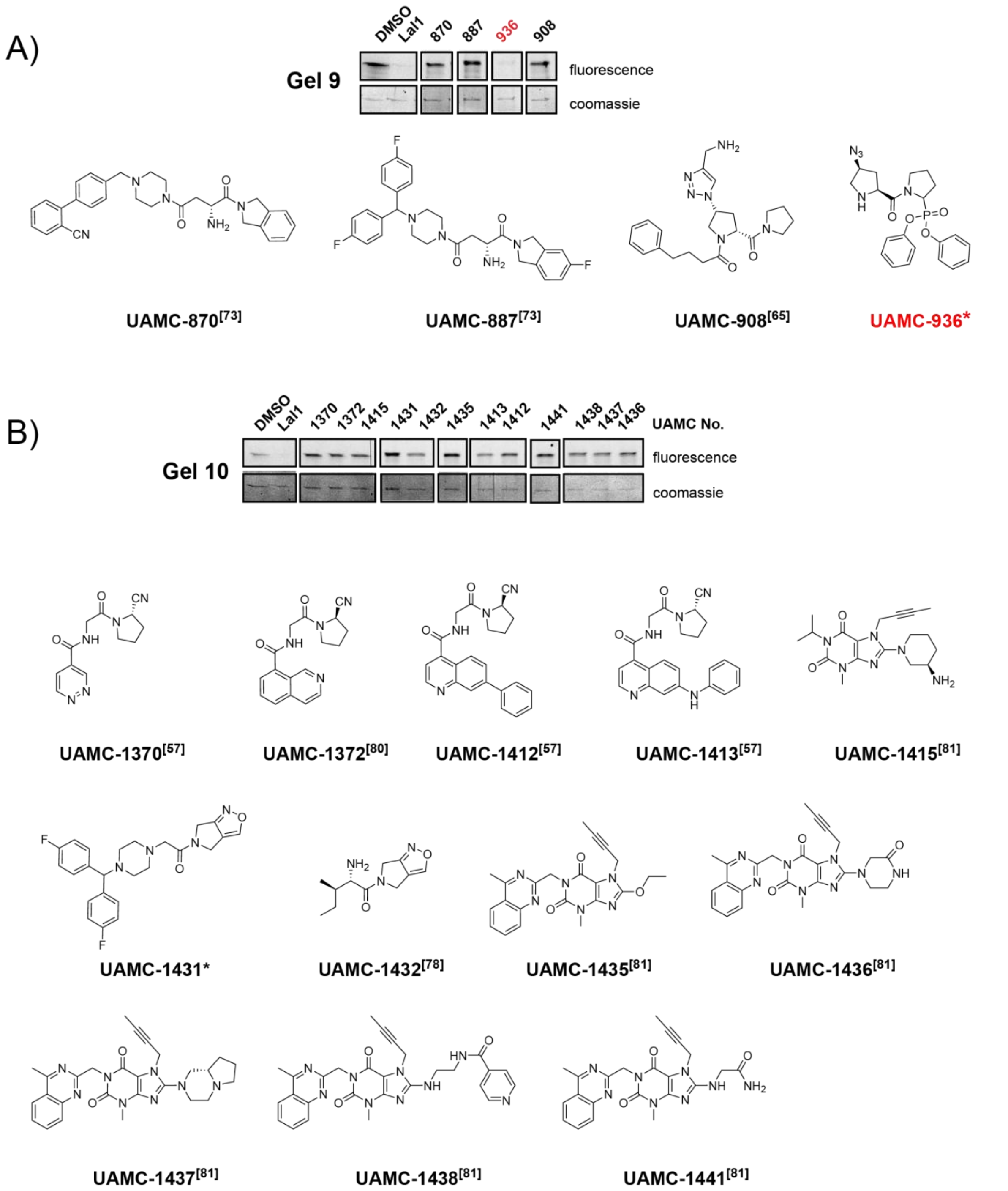


**Figure S12. Structures of the focused in-house compound library and initial screening against recombinant HI-A1.** Compounds (50 µM) were screened using Probe 15 (5 µM). Due to partial degradation of aged library members, compound purity was determined after screening, and only lanes corresponding to compounds with ≥70% purity are shown. Active compounds were defined as those causing >50% reduction of the probe-derived fluorescence signal relative to the DMSO control and are highlighted in red. Unpublished library members (*) are included for completeness; their HRMS and HPLC characterization data are provided in Section 8. Lal1 = Lalistat 1. 870, 887[73], 908[65], 1370, 1412, 1413[57], 1372[80], 1415, 1435, 1436, 1437, 1438, 1441[81], 1432[78]

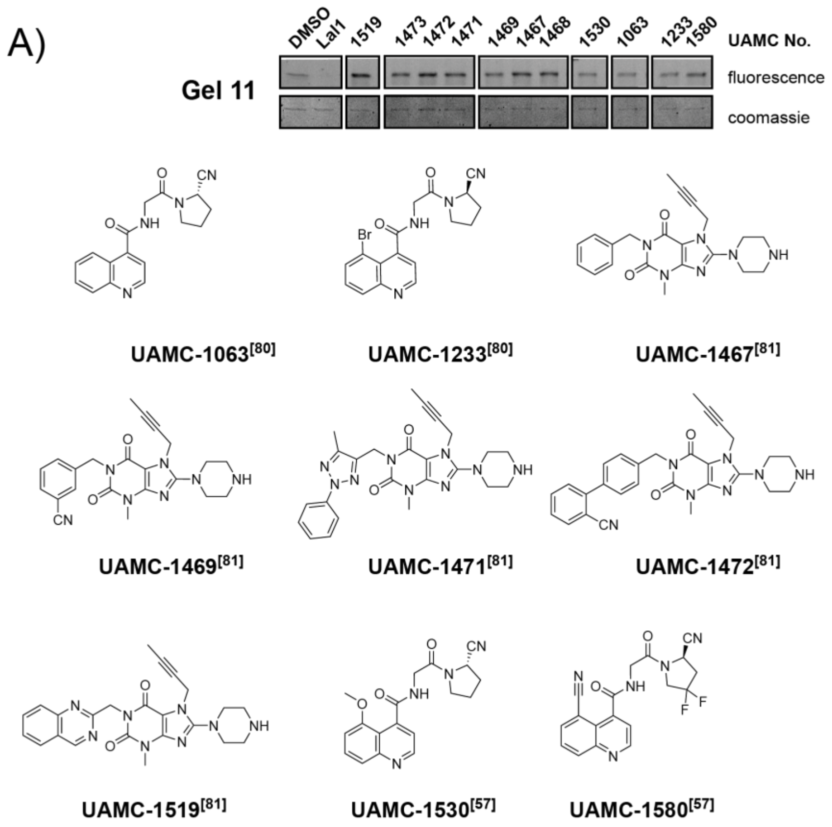


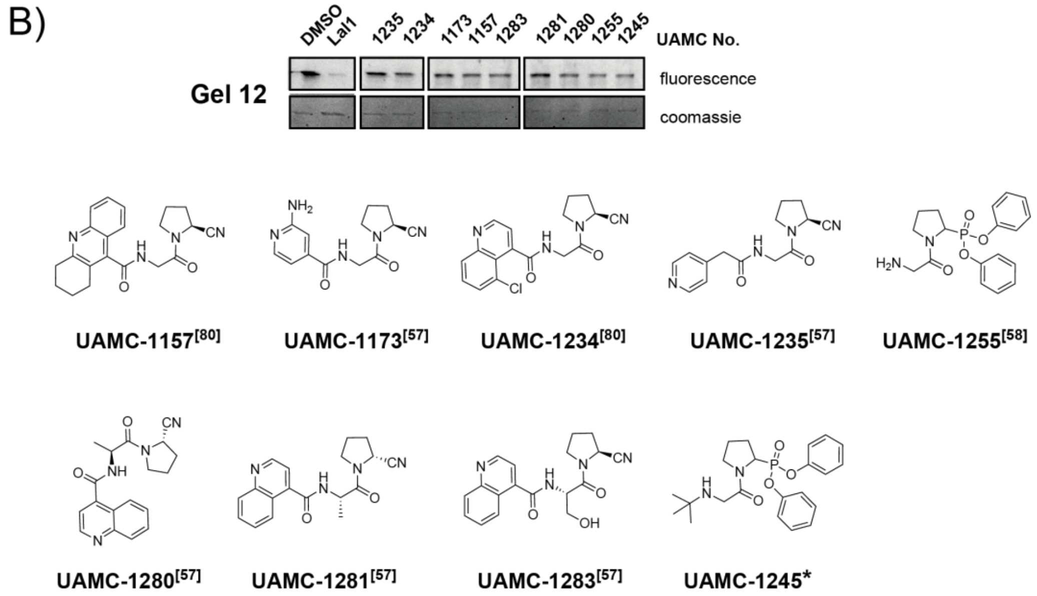


**Figure S13. Structures of the focused in-house compound library and initial screening against recombinant Hl-A1.** Compounds (50 µM) were screened using Probe 15 (5 µM). Due to partial degradation of aged library members, compound purity was determined after screening, and only lanes corresponding to compounds with ≥70% purity are shown. Active compounds were defined as those causing >50% reduction of the probe-derived fluorescence signal relative to the DMSO control. No active compounds were identified in these gels. Unpublished library members (*) are included for completeness; their HRMS and HPLC characterization data are provided in Section 8. Lal1 = Lalistat 1. 1063, 1233[80], 1467, 1468, 1469, 1471, 1472, 1473, 1519[81], 1530, 1580[57], 1157[80], 1173[57], 1234[80], 1255[58], 1235, 1280, 1281, 1283[57]

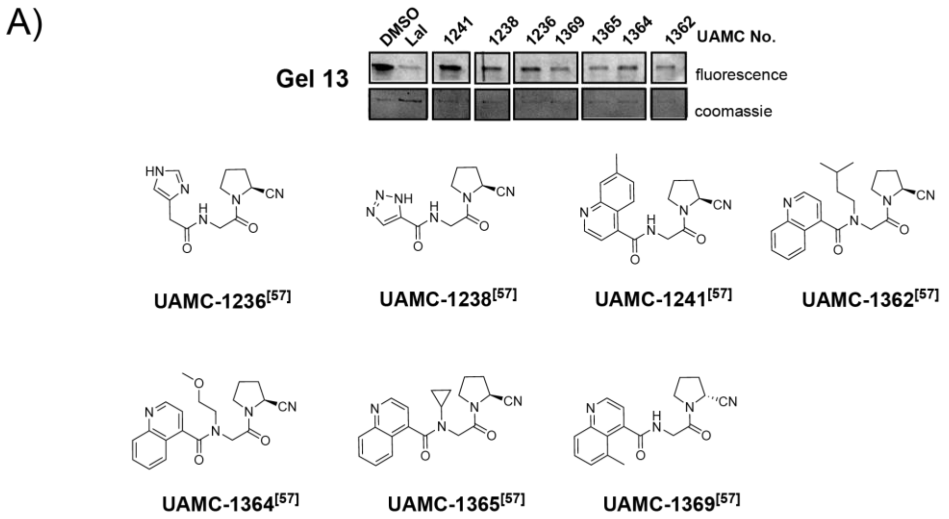


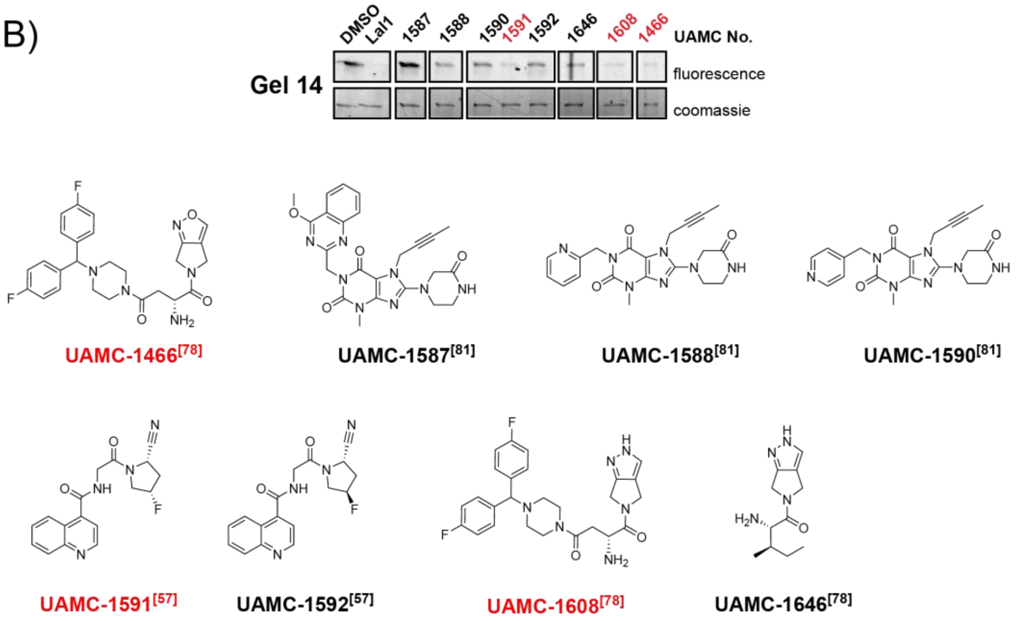


**Figure S14. Structures of the focused in-house compound library and initial screening against recombinant HI-A1.** Compounds (50 µM) were screened using Probe 15 (5 µM). Due to partial degradation of aged library members, compound purity was determined after screening, and only lanes corresponding to compounds with ≥70% purity are shown. Active compounds were defined as those causing >50% reduction of the probe-derived fluorescence signal relative to the DMSO control. No active compounds were identified in these gels. Unpublished library members (*) are included for completeness; their HRMS and HPLC characterization data are provided in Section 8. Lal1 = Lalistat 1. 1236, 1238, 1241, 1362, 1364, 1365, 1369[57], 1466[78], 1587, 1588, 1590[81], 1591, 1592[57], 1608, 1646[78]

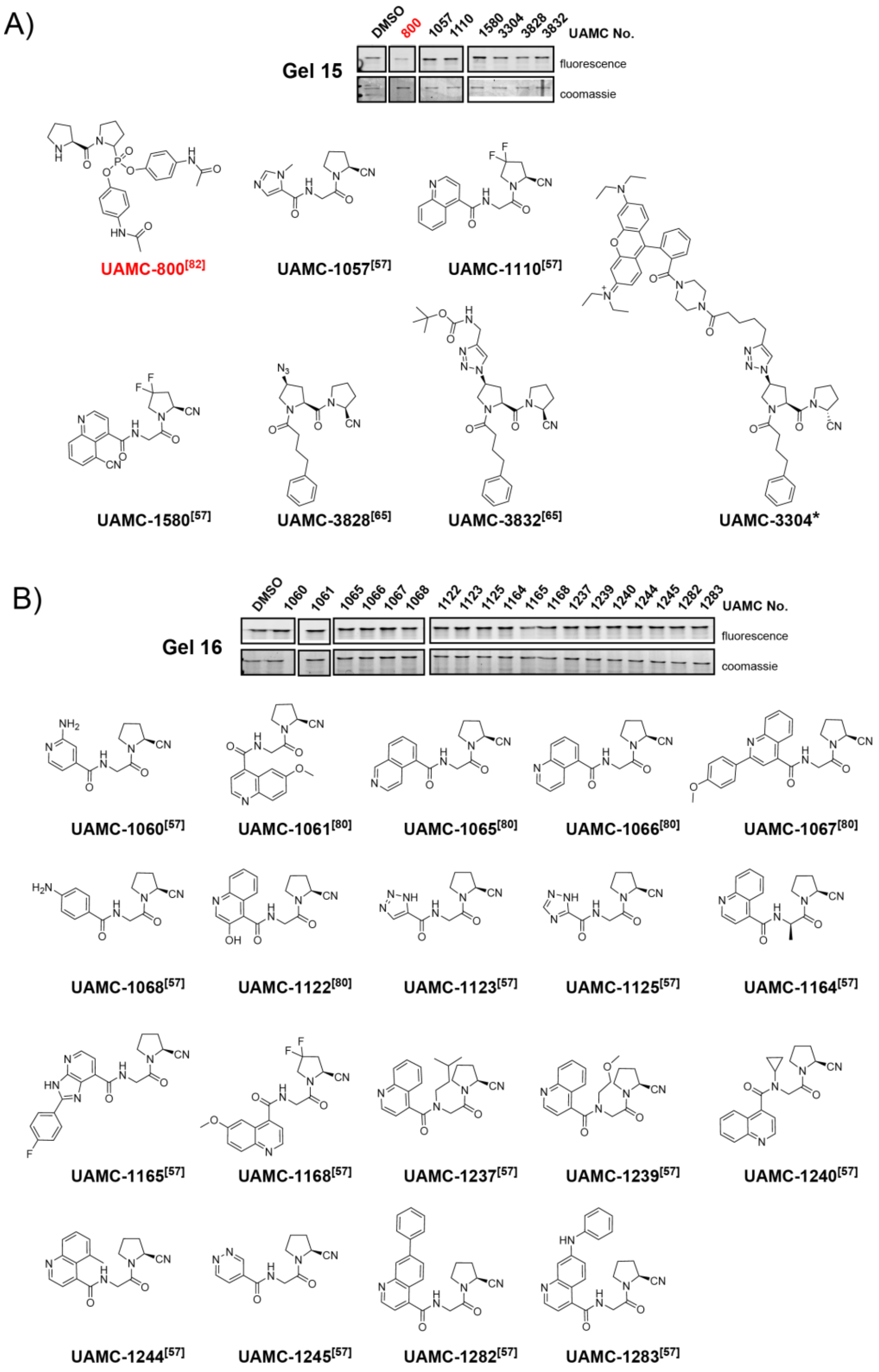


**Figure S15. Structures of the focused in-house compound library and initial screening against recombinant HI-A1.** Compounds (50 µM) were screened using Probe 15 (5 µM). Due to partial degradation of aged library members, compound purity was determined after screening, and only lanes corresponding to compounds with ≥70% purity are shown. Active compounds were defined as those causing >50% reduction of the probe-derived fluorescence signal relative to the DMSO control. No active compounds were identified in these gels. Unpublished library members (*) are included for completeness; their HRMS and HPLC characterization data are provided in Section 8. Lal1 = Lalistat 1. 800[82], 1057, 1110, 1580[57], 3828, 3832[65], 1060[57], 1061, 1065, 1066, 1067, 1122[80], 1068[57], 1123, 1125, 1164, 1165, 1168, 1237, 1239, 1240, 1244, 1245, 1282, 1283[57]

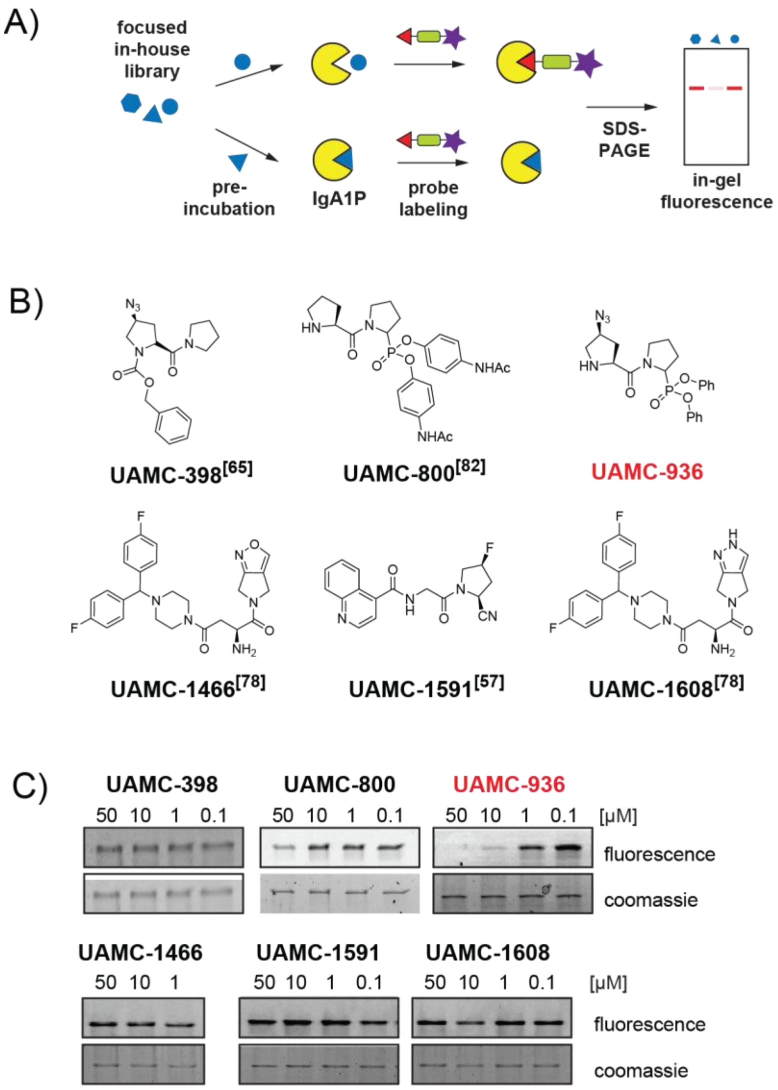


**Figure S16. Competitive activity-based protein profiling (cABPP) of a focused in-house library.** (A) Schematic of the gel-based competitive screening assay. Recombinant HI-A1 was preincubated with individual library members, followed by addition of probe 15. Covalent probe labelling was analyzed by in-gel fluorescence imaging. Reduced fluorescence intensity indicates competitive inhibition of probe engagement. (B) Initial screening of a focused in-house library at 50 µM (Figures S8-S15) resulted in the identification of inhibitors that reduce probe labelling of HI-A1. (C) Dose-response analysis of the primary hits in competition with probe 15 revealed UAMC-936 as the only compound that substantially inhibited probe labelling. 398[65], 800[82], 936, 1466[78], 1591[57], 1608[78]

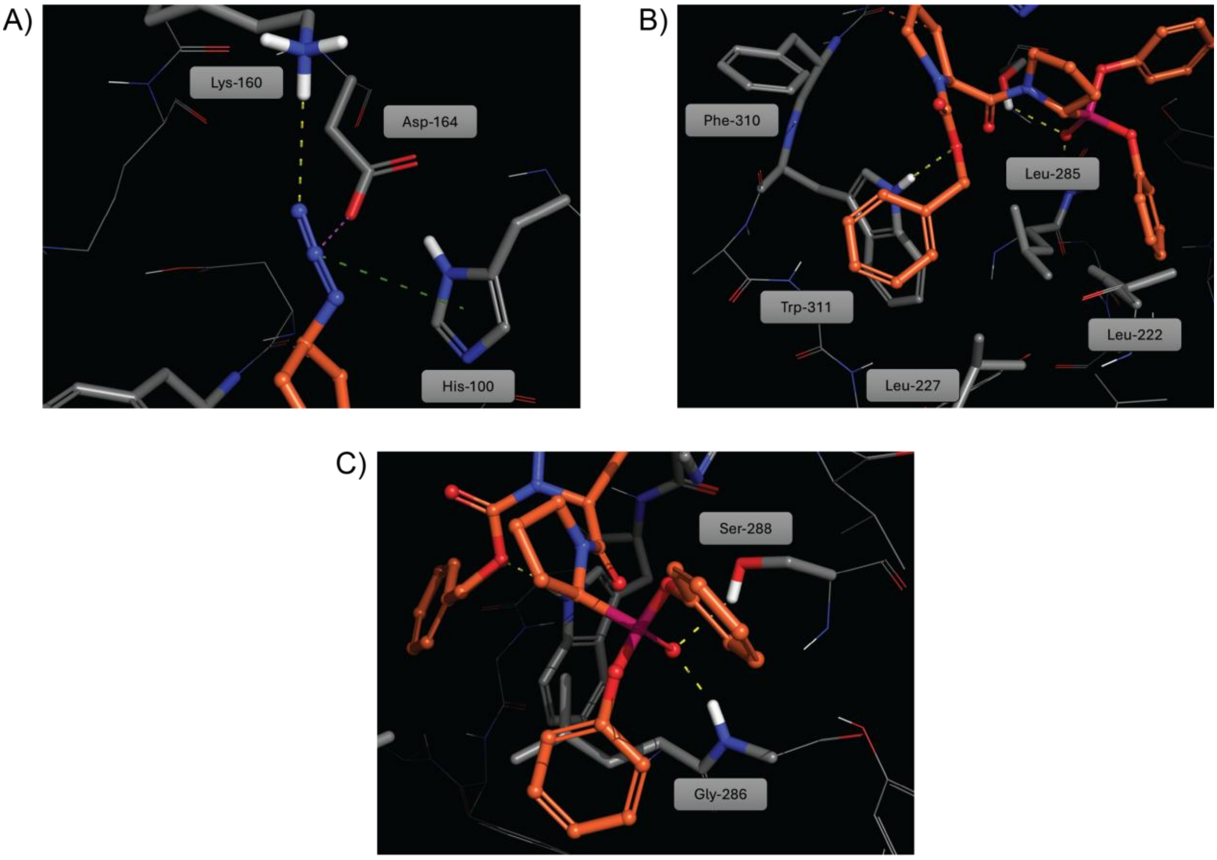


**Figure S17. Zoomed-in representation of the molecular docking of compound 4 in the active site of HI-A1.** A) The azide functionality accepts a hydrogen bond from Lys-160 and is involved in an ionic interaction with Asp-164 and a charge-pi interaction with His-100. B) Benzyloxycarbonyl (-Cbz) group is flanked by hydrophobic residues Phe-310, Trp-311, Leu-285 and Leu-222, and forms a hydrogen bond interaction with Trp-311. C) Phosphonate group is interacting with the catalytic Ser-288 and the backbone NH of Gly-286.

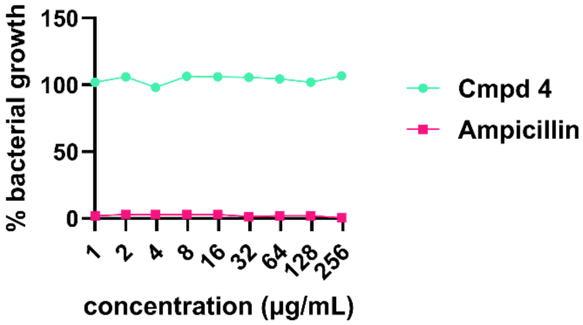


**Figure S18.** Minimum inhibitory concentration (MIC) curve of compound 4 against *H. influenzae* clinical isolate #3 (growth medium: supplemented brain heart infusion broth).

### 3. Biochemical, Microbiological and Computational Methods

#### A. IgA1P Expression and Purification

Competent *Escherichia coli* BL21 (DE3) Rosetta™ cells (Novagen) were thawed on ice, and plasmid DNA (Table S3) was added at 1:10 (v/v) DNA-to-cell ratio. The cells were incubated on ice for 30 mins followed by heat shock at 42 °C for 30s and returned on ice for 2 min. Super Optimal broth with Catabolite repression (SOC) medium (500 µL; New Englands BioLabs) was added, and cells were incubated at 37 °C with shaking for 1h. The transformation mixture was plated on Lysogeny Broth (LB) agar supplemented with the appropriate antibiotics and incubated overnight at 37 °C to obtain transformant colonies. A single colony was used to inoculate selective LB medium, and the resulting overnight culture was mixed with glycerol for cryogenic storage at −80 °C.

Recombinant protein expression and purification were performed following the protocol defined by Spoerry *et al.,* with minor modifications[44]. Transformed *E.coli* BL21 (DE3) Rosetta cells™ (New England Biolabs) were grown in Terrific Broth (Thermo Scientific™) supplemented with 50 µg/mL Kanamycin sulfate (Merck) at 37 °C with shaking until the culture reached an $OD_{600}$ of 1.8-2. The temperature was reduced to 18 °C, and protein expression was induced with 0.5 mM IPTG for 16 h while shaking. Cells were harvested by centrifugation, and the resulting cell pellet was resuspended in lysis buffer (100mM 4-(2-hydroxyethyl)-1-piperazineethanesulfonic acid (HEPES), 500mM NaCl, 10mM imidazole, 10% glycerol, 0.5 mM tris(2-carboxyethyl)phosphine (TCEP), pH 8). Cells were lysed by sonication on ice. The lysate was further clarified by centrifugation followed by filtration through a 0.45 µm membrane. IgA1P purification was performed at 4 °C on ÄKTA start™ system using UNICORN start software. Lysates were loaded onto a 1 mL HisTrap HP column (Cytiva) at a flow rate of 1 mL/min. The column was equilibrated according to manufacturer's instructions. His-tagged proteins were washed with 10 column volumes of wash buffer (20 mM HEPES, 500 mM NaCl, 10 mM imidazole, 10% (v/v) glycerol, 0.5 mM TCEP, pH 7.5) and eluted with elution buffer (20 mM HEPES, 500 mM NaCl, 500 mM imidazole, 10% (v/v) glycerol, 0.5 mM TCEP, pH 7.5) at a flow rate of 1 mL/min. Fractions corresponding to the elution peak were collected and analyzed by SDS-PAGE (12.5%). Fractions containing the protease were pooled for further use. The protein samples were concentrated using a 5 mL HiTrap™ desalting column packed with Sephadex G-25 resin at a flow rate of 5 mL/min. The column was first equilibrated with storage buffer (20 mM HEPES, 300 mM NaCl, 10% (v/v) glycerol, 0.5 mM TCEP, pH 7.5), after which the pooled protein sample was loaded onto the column, and protein was subsequently eluted in storage buffer. Protein concentration was determined at 280 nm using a NanoDrop™ spectrophotometer, yielding values of 1.4 mg/mL for HI-A1 wt, 1.1 mg/mL for HI-A1 mt and 0.481 mg/mL for NM1 wt. Samples were aliquoted and stored at -80 °C.

#### B. (Competitive) IgA1 Cleavage Assay

Recombinant IgA1Ps (20 µg/mL) were incubated with IgA from human serum (0.25 mg/mL; Sigma-Aldrich) at 37 °C for 2 h in IgA1P reaction buffer (25mM HEPES, pH 7.5). Serum IgA consists predominantly of IgA1 (~90%)[83]. For competitive assays, inhibitors at the indicated concentrations were pre-incubated with the supernatant for 30 min at RT prior to IgA addition. Reactions were quenched with 5x SDS loading buffer, and the cleavage products were resolved by SDS-PAGE and visualized using FastGene™ Q-stain.

#### C. ABP Labelling of Recombinant Enzymes and Tissue Samples

**Labelling of recombinant IgA1Ps with alkyne-ABPs:** Recombinantly expressed IgA1Ps (20 µg/mL) were incubated with the indicated concentrations of alkyne-tagged ABPs 1-14, or DMSO (vehicle control) for 2h at 37 °C in IgA1P reaction buffer. Post incubation, copper-catalyzed azide alkyne click chemistry (CuAAC) was performed in 25 µL total reaction volumes, containing 5-TAMRA-PEG3-azide (25 µM; Carl Roth), THPTA (50 µM; BaseClick GmbH), $CuSO_4$ (1 µM; Sigma-Aldrich) and Ascorbic acid (1 µM; Thermo Scientific™). Reactions were vortexed briefly, and incubated at 1 h, RT in the dark. Reactions were quenched with 2x SDS loading buffer, and the samples were analyzed using SDS-PAGE with in-gel rhodamine fluorescence imaging followed by total protein staining with FastGene Q-stain.

**Labelling of recombinant IgA1Ps with fluorescent ABPs:** Recombinantly expressed IgA1Ps (20 µg/mL) were incubated with the indicated concentrations fluorescent ABPs 15-17 in IgA1P reaction buffer. Samples were vortexed briefly, and incubated at 1 h, 37 °C in the dark. Reactions were quenched with 50 µL 2x SDS loading buffer, and the samples were analyzed using SDS-PAGE with in-gel fluorescence imaging followed by total protein staining with FastGene Q-stain.

**Competitive ABP labelling of recombinant IgA1Ps:** Recombinantly expressed IgA1Ps (20 ug/mL) were pre-incubated with the indicated concentrations of inhibitors for 1h at RT in IgA1P reaction buffer. ABPs were then added at the indicated concentrations and samples were incubated for 2 h at 37 °C. Reactions were quenched with 50 µL 2x SDS loading buffer. Samples were analyzed using 2x SDS-PAGE, with in-gel fluorescence imaging followed by total protein staining with FastGene Q-stain.

**(Competitive) ABP labelling in mouse tissue lysates:** Mouse tissue samples were provided by Prof. Greetje Vande Velde (KU Leuven). All animal procedures were approved by the Ethical Committee for Animal Experimentation of KU Leuven and conducted in accordance with institutional guidelines. A surplus C57BL/6 mouse from the KU Leuven animal facility was euthanised by intraperitoneal injection of pentobarbital (Dolethal, overdose). Lung and brain were collected and homogenised in PBS buffer (~ 1mL per organ) using a Tissue Master homogeniser (OMNI International). Lysates (1 mg/mL) were incubated with indicated concentrations of fluorescent ABPs for 2h at 37 °C. Reactions were quenched with 2× SDS loading buffer and samples were analyzed by SDS–PAGE followed by in-gel fluorescence imaging and total protein staining with FastGene Q-stain. For competitive profiling, lysates were pre-incubated with inhibitors at the indicated concentrations for 30 min at RT prior to ABP labelling. For labelling of IgA1Ps in a proteome background, lysates were spiked with recombinant IgA1P (20 µg/mL).

**ABP labelling of human trypsin and chymotrypsin:** Trypsin from human pancreas (T6424, Sigma-Aldrich, 10 µg/mL) and chymotrypsin from human pancreas (230900, Sigma-Aldrich, 10 µg/mL) were incubated with the indicated concentrations of ABPs or DMSO (vehicle control) for 2 h at 37 °C. After incubation, a click reaction was performed for Probe 1 and DMSO samples for 1h in the dark. All samples were quenched with 2x SDS loading buffer and were analyzed using SDS-PAGE with in-gel fluorescence imaging followed by total protein staining with FastGene Q-stain.

#### D. Residual Enzyme Activity Assays

**Residual activity of DPP's:** DPP4 was purified from human seminal plasma as described previously[84]. Gateway entry clones for human DPP8 (Accession no. DQ891733) and DPP9 (Accession no. DQ892325) were obtained from Dharmacon, and the recombinant protein expression in Sf9 insect cells were done using N-terminal BaculoDirect kit (Life Technologies). Both the proteases were purified using immobilized $Ni^{2+}$-chelating chromatography (GE healthcare, Diegem, Belgium), followed by anion-exchange chromatography using a 1 mL Mono Q (GE healthcare, Diegem, Belgium). Enzyme concentrations used in the final assay mixtures were 0.39 nM (DPP4), 2.25 nM (DPP8) and 1.53 nM (DPP9). Enzyme activity was measured using Ala-Pro-p-nitroanilide (Bachem) as the substrate at 25 µM (DPP4), 300 µM (DPP8) or 150 µM (DPP9) at pH 7.4 (0.05 M HEPES-NaOH buffer with 0.1 % Tween-20, 0.1 mg/mL BSA and 150 mM NaCl). Probe 1 was applied at 1 and 10 µM final concentrations, and pre-incubated with the enzyme for 15 minutes at 37 °C before the reactions were initiated with substrate addition. pNA release was measured kinetically at 405 nm for at least 15 minutes at 37 °C. Measurements were done on the Infinite 200 (Tecan Group Ltd.) and the Magellan software was used to process the data.

**Residual activity of FAP and PREP:** Recombinant human PREP was expressed in BL21(DE3) cells and purified using immobilized Co-chelating chromatography (GE healthcare), followed by anion-exchange chromatography on a 1 ml Mono Q column (GE healthcare). Gateway entry clones for human FAP (Accession number DQ891423; Dharmacon) and the human secretion signal was replaced with the HoneyBee mellitin secretion signal. FAP expression in Sf9 cells were performed using C-terminal BaculoDirect kit (Life Technologies). FAP was purified from the insect cell supernatant using immobilized Ni-chelating chromatography (GE healthcare, Diegem, Belgium), followed by anion-exchange chromatography using a 1 mL HiTrap Q and size exclusion chromatography using the Superdex 200 column (GE healthcare, Diegem, Belgium). FAP enzyme activity was measured using Z-Gly-Pro-7-amino-4-methylcoumarine (Bachem) at 50 µM in 0.05 M Tris-HCl buffer (pH 8) containing 1 mg/mL BSA and 140 mM NaCl, while PREP screening used N-succinyl-Gly-Pro-7-amino-4-methylcoumarine (AMC; Bachem) at 250 µM in 0.1 M K-phosphate buffer (pH 7.8) containing 1 mM EDTA, 1 mM DTT and 0.1 mg/mL BSA. For both enzymes, Probe 1 was tested at 1 and 10 µM final concentrations, and pre-incubated with the enzyme for 15 minutes at 37 °C before the reactions were initiated with substrate addition. AMC release was measured kinetically at $\lambda_{ex}$= 380 nm and $\lambda_{em}$= 465 nm for at least 15 minutes at 37 °C. Measurements were done on the Infinite 200 (Tecan Group Ltd.) and the Magellan software was used to process the data.

#### E. Microbiology Experiments

**Cultivation of *H. influenzae* Clinical Isolates:** Brain Heart Infusion (BHI) medium was prepared by dissolving BHI powder (Oxoid CM1135; 37g/L) in Milli-Q® water followed by sterilization at 121 °C for 1h. The medium was supplemented with X factor Hemin (Thermo Scientific™ 11499093) and V factor β-nicotinamide adenine dinucleotide (NAD, Sigma-Aldrich N7381), each at 10 µg/mL, to obtain supplemented BHI (sBHI). Hemin stock (10 mg/mL) was prepared by dissolving hemin in a minimal volume of sterile 0.1M NaOH and bringing it to final volume with sterile water to ensure solubility. *H. influenzae* clinical isolates (Table S2) were plated onto Haemophilus Test medium (HTM) agar plates (Thermo Scientific™) and incubated at 37 °C with 5% $CO_2$ for 16-18h. Colonies were inoculated into sBHI (15 mL) and grown overnight at 37 °C with 5% $CO_2$. Cultures (OD600 = 1–2) were harvested by centrifugation (4500 rpm, RT, 15 min), and the supernatants were collected and concentrated four-fold using 50 kDa Amicon filters (4000 × g, 4°C, 10 min).

**(Competitive) IgA1 cleavage in *H. influenzae* clinical isolates:** *H. influenzae* culture supernatant was incubated with IgA from human serum (0.4 mg/mL; Sigma-Aldrich) for 2 h at 37 °C. For competitive assays, inhibitors at the indicated concentrations were pre-incubated with the supernatant for 30 min at RT prior to IgA addition. Reactions were quenched by adding 5x SDS loading buffer and analyzed by SDS-PAGE followed by protein staining with FastGene™ Q-stain.

**ABP labelling of endogenous IgA1Ps**: *H. influenzae* culture supernatants were incubated with probe 15 for 2 h at 37 °C in the dark. For competitive assays, inhibitors at the indicated concentrations were pre-incubated with the supernatant for 30 min at RT before probe addition. Reactions were quenched with 2x SDS loading buffer and analyzed by SDS–PAGE with in-gel fluorescence imaging followed by protein staining (FastGene Q-stain).

**Minimum Inhibitory Concentration (MIC) assay**: MICs of compound 4 were determined against *H. influenzae* clinical isolate #3 using the broth microdilution method in sterile 96-well plates in three technical replicates to assess any growth-inhibitory effects. Overnight bacterial cultures were prepared as described above, grown till OD600 of ~1, diluted 1:50 in fresh sBHI, and further cultured to mid-log phase (OD600 = 0.4-0.5) prior to inoculum preparation. Cultures were then diluted to $1 \times 10^6$ CFU/mL, to obtain a final inoculum of $5 \times 10^5$ CFU/mL per well after 1:1 mixing with compound solutions. Two-fold serial dilutions (2x working concentrations) of compound 4 were prepared in sBHI starting from 1 mg/mL and yielding concentrations of 500, 250, 125, 62.5, 31.3, 15.6, 7.8, 3.9 and 2.0 ug/mL. Subsequently, 100 µL of each 2x compound dilution was added to wells in technical replicates, followed by 100 µL of the bacterial suspension. Ampicillin was included as an antibiotic control at the same concentration range. Growth controls (100 µL sBHI + 100 µL bacterial suspension) and sterility controls (200 µL sBHI) were also included. Plates were incubated at 37 °C with 5% $CO_2$ for 18h, and bacterial growth was monitored both visually and by measuring OD600 values.[85-86] Raw OD600 values from triplicate wells were averaged for each concentration. Background absorbance was corrected by subtracting the mean OD of blank wells containing BHI medium only. Percent growth was calculated relative to the untreated growth control according to the formula: % growth = (Corrected OD sample / Corrected OD growth control) × 100. Data were plotted against antimicrobial concentration (µg/mL) on a $\log_2$-scaled x-axis. All analyses were performed using GraphPad Prism software version 11.

#### F. Flow cytometry

*H. influenzae* clinical isolate #3 was grown overnight at 37 °C under 5% $CO_2$ in sBHI medium. Cultures were grown to an OD600 of ~1, corresponding to about $2 \times 10^7$ CFU/mL, as determined previously through serial dilution plating assay. Cells were harvested (3500 rpm, 10 mins, RT) and the culture supernatant was concentrated approximately four-fold using 50 kDa Amicon filters (4000 × g, 4°C, 10 min) before resuspending the pellet to a final volume of 1 mL. For analysis, 200 µL of the suspension, corresponding to $5 \times 10^6$ bacteria, was pre-incubated with compound 4 at 10 and 100 µM for 30 min at RT. Samples were then incubated with colostrum-derived secretory IgA (50 µg/mL; IgA from human colostrum, I2636, Sigma Aldrich), which contains antibodies recognizing *H. influenzae* antigens[87], for 30 min at 37 °C. After two washes with PBS containing 1% BSA, bound IgA was detected using FITC-labeled goat anti-human IgA1 (A18782, Invitrogen; 1:500 dilution). FITC antibody incubation was performed in PBS containing 0.1% BSA for 45 min at 4 °C in the dark. Cells were washed and fixed with 2% paraformaldehyde (PFA) in PBS for 30 min at RT, and fluorescence intensity was measured using an Aurora flow cytometer (Cytek Biosciences).

Flow cytometry data were analyzed using FlowJo software (version 10.10.0 BD Biosciences). FITC-positive bacteria were gated relative to the unstained and FITC background controls. The percentage of FITC positive bacteria (% FITC$^+$) was calculated as the frequency of FITC positive events within the bacterial singlet population. Statistical analyses were performed on GraphPad Prism software version 11. Differences between the conditions were determined using ordinary one-way ANOVA, followed by Dunnett's multiple comparison test with the negative or un-treated control as the reference. Data represent 3 technical replicates per condition. A p value of ≤ 0.05 was considered a statistically significant value.

##### G. Docking studies

Molecular docking of compound 4 in the HI-A1 active site (pdb_00003h09) was done with Glide[88]. Standard parameters were applied and the active site was defined by centering on the catalytic Ser-288. No constraints were used. Visualization was done with Maestro (Schrödinger Release 2025-4: Maestro, Schrödinger, LLC, New York, NY, 2025.) and PyMol (The PyMOL Molecular Graphics System, Version 2.5.7, Schrödinger, LLC.).

#### 4. Synthetic Schemes

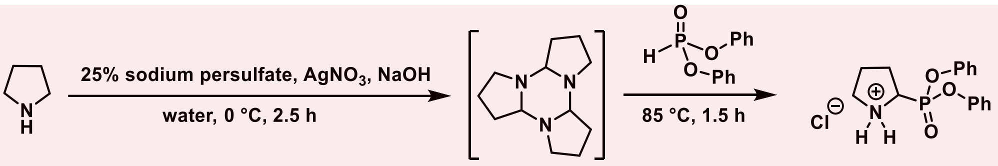


**Scheme S1.** Warhead synthesis.

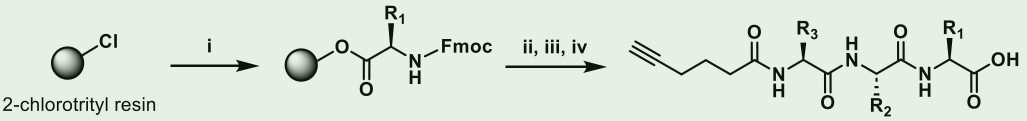


**Scheme S2. Synthesis of peptide linkers.** (i) Fmoc-amino acid, DIPEA, DCM, o/n; (ii) Repeat cycles: (1) 20% piperidine/DMF, 30 min. + 15 min.; (2) Fmoc-amino acid, HATU, DIPEA, DMF; (iii) 5-hexynoic acid, HOBt, DIC, DMF; (iv) 95:5 TFA:DCM or 1:99 TFA:DCM.

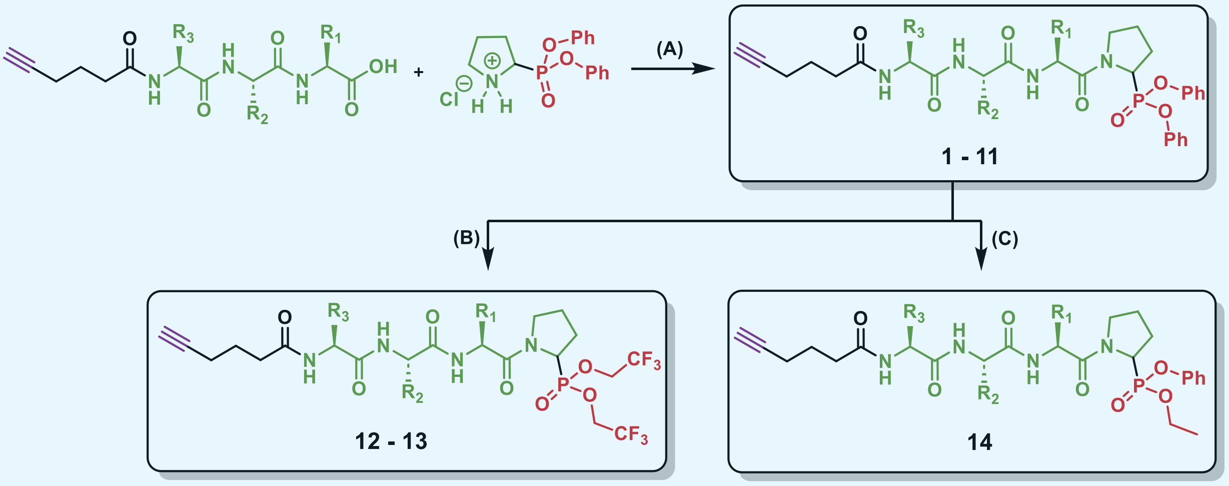


**Scheme S3. Synthesis of the activity-based probes.** (A) HATU, DIPEA, DMF, r.t., o/n. (B) KF, 18-crown6, 2,2,2-trifluoroethanol, 80 °C – 15 min. to r.t. 16 h. (C) (i) KF, 18-crown6, ethanol, 80 °C – 15 min. to r.t. 16 h; (ii) LiBr, 2-butanone, reflux, 18 h; (iii) Phenol, PyBOP, DIPEA, 4Å MS, DMF, r.t., 48 h.

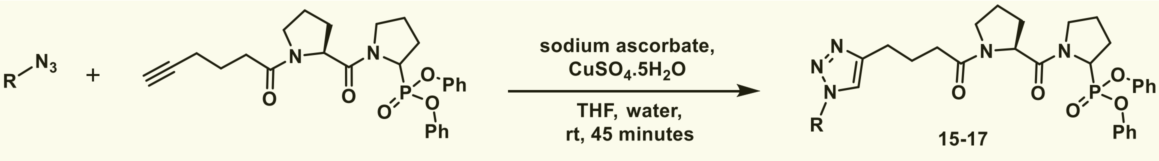


**Scheme S4. Fluorophore attachment using click-chemistry** (R = TAMRA, Cy5, Sulfo-Cy5).

### 5. Synthesis of Probes and Compounds

**Materials and methods:** All starting materials, reagents, and solvents were used as received without further purification. For liquid-phase synthesis, glassware and Teflon-coated stir bars were pre-dried at 110 °C for at least 4 hours prior to use. Liquid-phase reactions were carried out under an inert nitrogen atmosphere using balloons and dry solvents. Solid-phase peptide synthesis (SPPS) was performed under ambient conditions in syringes fitted with polypropylene filters. Flash chromatography was conducted on a Biotage Isolera One purification system equipped with an internal variable dual-wavelength diode array detector (200–400 nm). Reverse-phase purifications employed Biotage® Sfär Bio C4 D cartridges. Samples were applied to the columns by dry loading, using self-packed sample cartridges with Celite 545 as the loading medium.

**High-resolution mass spectrometry (HRMS) sample preparation:** The final compounds were dissolved in methanol to obtain a concentration of $10^{-2}$ M. This solution was further diluted with a 90:10 (v/v) methanol–water mixture to reach a final concentration of $10^{-5}$ M.

**Instrumentation:** NMR spectra were recorded on a Bruker Avance III Nanobay spectrometer equipped with an Ultrashield magnet, operating at 400 MHz for $^1$H nuclei. Spectra were processed and analyzed using MestReNova. Chemical shifts ($\delta$) are reported in parts per million (ppm) relative to the residual solvent peak, and coupling constants (J) are in hertz (Hz). The signal multiplicities are designated as m = multiplet. Ultra-performance liquid chromatography (UPLC) was used to determine the product purity. Analyses were performed on a Waters ACQUITY UPLC H-Class system equipped with a TUV detector coupled to a Waters QDa single-quadrupole mass spectrometer. Separations were carried out on a Waters ACQUITY UPLC BEH C18 column (1.7 µm, 2.1 mm × 50 mm). The mobile phase consisted of these solvents: $H_2O$ (solvent A), ACN (solvent B), 0.1% FA in $H_2O$ (solvent C), and 0.1% FA in ACN (solvent D). The gradient elution was programmed as follows: the column was first equilibrated with 95% solvent C and 5% solvent D for 0.15 min, followed by a linear increase of solvent D to 100% over 1.75 min. This composition was held constant for 0.25 min before returning to the initial conditions (95% C/5% D) for 0.75 min at a flow rate of 0.7 mL/min. Mass spectra were recorded over an *m/z* range of 100–1000; for compounds 15–17, the range was extended to 100–1300. UV detection was performed at 254 nm. Preparative high-performance liquid chromatography (PREP-HPLC) was performed on a Shimadzu LC-20 Prominence system equipped with an LC-20AD solvent delivery module and an SPD-20A UV–Vis detector. Compounds **11-13** were purified using a Phenomenex Luna C18(2) column (10 µm, 100 Å, 250 x 10 mm) with a mobile phase of acetonitrile + 0.1% TFA and water + 0.1% TFA. High-resolution mass spectra were acquired using a Synapt HDMS Q-TOF instrument (Waters, Manchester, UK). The MS was calibrated prior to use with a 0.1% $H_3PO_4$ solution. Spectra were lock mass corrected with the known mass per charge ratio (m/z) of the nearest $H_3PO_4$ cluster ion, obtained from a separate injection of a reference sample containing 0.1% phosphoric acid. Each sample (5 µL, $10^{-5}$ M) was injected via an Acquity UPLC system (Waters, Manchester, UK) using a mobile phase consisting of 90:10 (v/v) MeOH/$H_2O$ containing 0.1% formic acid, at a flow rate of 40 µL/min. Ionization was performed using a standard electrospray ionization (ESI) source, with sample injections spaced at 2-minute intervals. Analytes were detected as protonated and/or sodiated molecular ions.

#### A. General procedures

**General procedure 1 – solid phase peptide synthesis:**

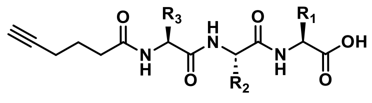

Peptides were synthesized on chlorotrityl resin as the solid support using Fmoc-protected amino acids. For preparing the resin containing trityl residues for loading, the resin (1.0 eq.) was swollen in DCM for 30 minutes in a cartridge for SPPS. After draining the DCM, the resin was loaded by agitating a mixture of the first Fmoc-amino acid (3.0 eq.) and DIPEA (6.0 eq.) with the resin in DCM overnight. After draining the loading mixture, the resin was washed with DCM (3x). Unreacted sites on the resin were capped using DCM/MeOH/DIPEA mixture (8.5:1:0.5, v/v/v) for 45 minutes. The resin was then washed sequentially with DCM (5x) and DMF (5x).

Fmoc-group on the N-terminus was removed with 2 treatments of 20% piperidine in DMF for 30 minutes and 15 minutes, respectively, followed by washing with DMF (5x). Subsequent couplings were carried out by pre-activating the incoming Fmoc-amino acid (3.0 eq.) with HATU (3.0 eq.), and DIPEA (6.0 eq.) in DMF for 15 minutes. The resulting solution was added to the resin and agitated overnight. When coupling was completed the cartridge was drained, the resin washed with DMF (3x) and the next deprotection and coupling was performed. This was repeated until all desired amino acids were coupled.

After completion of the desired sequence, 5-hexynoic acid (3.0 eq.), HOBt (3.0 eq.), and DIC (3.0 eq.) were added to the resin to introduce the terminal alkyne tag and agitated for 3 hours. The resin was washed with DMF (3x) followed by DCM (3x). Peptides were cleaved from the resin using TFA in DCM (1% TFA in DCM, 2 minutes, 10x for protected peptides; 50% TFA in DCM, 2 hours, 1x for unprotected peptides). The combined filtrate was concentrated under reduced pressure, and the crude peptide was purified by reverse phase column chromatography (5-60%ACN/water), followed by lyophilization to afford the desired peptides as colorless oils or solids (depending upon the product).

**General procedure 2 – ABP synthesis:**

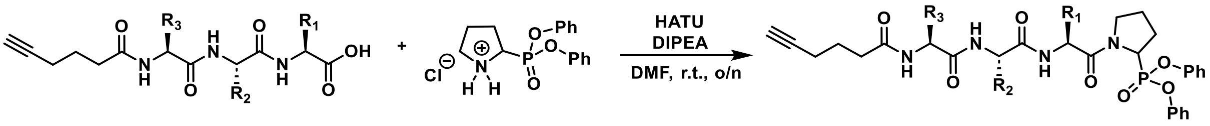


Under an inert atmosphere, the peptide (1.0 eq.) was dissolved in DMF (1.5 mL) followed by the addition of HATU (1.0 eq.) and DIPEA (5.0 eq.). The reaction mixture was stirred at room temperature for 30 minutes. Subsequently, 2-(diphenoxyphosphoryl)pyrrolidin-1-ium chloride (1.0 eq.) was added, and the reaction mixture was stirred overnight at room temperature. Reaction completion was monitored by UPLC-MS, the solvent was removed under reduced pressure, and the crude product was purified by reverse phase column chromatography to afford the desired compound.

**General procedure 3 – *t*-butyl group deprotection:**

The protected-compounds obtained from the peptide and the warhead coupling were subjected to 50% TFA/DCM for 1 hour followed by concentration under reduced pressure. The crude product was purified by reverse phase column chromatography to afford the desired compound.

**General procedure 4 – transesterification of ABPs:**

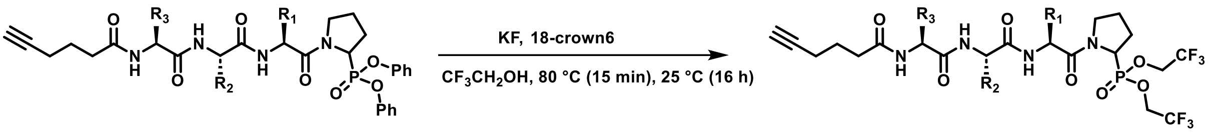


The diphenyl phosphonate-based activity-based probe (1 eq.) was dissolved in 2,2,2-trifluoroethanol. Potassium fluoride (10 eq.) and 18-crown-6 (0.068 eq.) were added, and the reaction mixture was stirred at 80 °C for 15 minutes, then allowed to stir at room temperature for 16 hours. The mixture was concentrated under reduced pressure, and the crude residue was purified by reverse phase column chromatography to afford the desired compound.

**General procedure 5 – synthesis of fluorescent probes:**

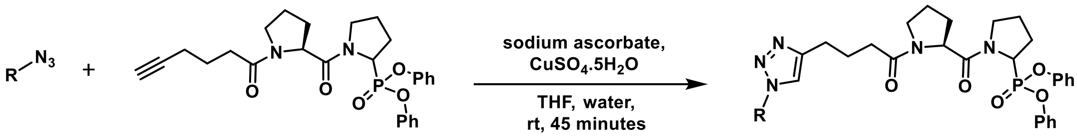


Diphenyl (1-(hex-5-ynoyl-L-prolyl)pyrrolidin-2-yl)phosphonate (1.0 eq.) and the desired fluorophore (1.3 eq.) were dissolved in THF/$H_2O$ (3:2 v/v). Sodium (R)-2-((S)-1,2-dihydroxyethyl)-4-hydroxy-5-oxo-2,5-dihydrofuran-3-olate (2.6 eq.) was added, followed by the addition of copper(II) sulfate pentahydrate (1.3 eq.). The reaction mixture was stirred at room temperature for 45 minutes. The solvent was removed under reduced pressure, and crude mixture was purified by reverse phase column chromatography to afford the fluorescent probe.

#### B. Synthesis

##### Synthesis of 2-(diphenoxyphosphoryl)pyrrolidin-1-ium chloride

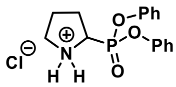

In a round bottom flask, pyrrolidine (12.2 mL, 150 mmol, 1.0 eq.) was dissolved in water (143 mL), sodium hydroxide (12 g, 300 mmol, 2.0 eq.) and silver nitrate (128 mg, 0.75 mmol, 0.005 eq.) was added. The reaction mixture was cooled to 0 °C and a 25% aqueous solution of sodium persulfate (35.7 g, 150 mmol, 1.0 eq.) was added dropwise while maintaining the temperature at ~0 °C. Upon completion of the addition, the reaction was stirred at the same temperature for 2.5 hours. The reaction mixture was then saturated with sodium chloride and extracted with DCM (4x). The combined organic layers were dried over sodium sulfate, filtered, and concentrated under reduced pressure at room temperature. The crude residue was dissolved in diethyl ether (15 mL), and the resulting precipitate was filtered off (2x). The filtrate was combined and concentrated under reduced pressure to afford the pyrroline trimer as dark orange oil. The crude pyrroline trimer was used directly in subsequent reaction without further purification.[40]

In a round bottom flask, pyrroline trimer (5.53 g, 26.67 mmol, 1.0 eq.) and diphenylphosphite (15.4 mL, 80.02 mmol, 3.0 eq.) were stirred at 85 °C for 1.5 hours. Upon reaction completion, dry diethyl ether (120 mL) was added to the viscous reaction mixture, resulting in the formation of a yellow precipitate. The precipitate was allowed to settle down, and the supernatant was transferred to another dry flask. This process was repeated twice with additional portions of dry diethyl ether (30 mL each). A solution of HCl in diethyl ether was then added to the combined filtrates, and the mixture was stirred at room temperature for 30 minutes. The excess solvent was decanted, and the resulting viscous residue was dried under reduced pressure. The material gradually turned into a foamy solid, which was recrystallized from acetone. The recrystallization was repeated twice to afford the titled product as a baby pink solid (8.28 g, 73% yield). The analytical data was consistent with those reported in the literature. **$^{1}$H NMR** ($D_2O$, 400 MHz): $\delta$ 7.43 – 7.38 (m, 4H), 7.33 – 7.29 (m, 2H), 7.18 – 7.13 (m, 4H), 4.44 – 4.37 (m, 1H), 3.58 – 3.47 (m, 2H), 2.60 – 2.52 (m, 1H), 2.41 – 2.31 (m, 1H), 2.27 – 2.15 (m, 2H).[41]

##### Synthesis of Alk-(L)-Pro-Pro-DPP (Probe 1)

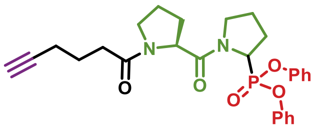


Using general procedure 2, hex-5-ynoyl-L-proline (153.0 mg, 0.730 mmol, 1.0 eq.), HATU (278.0 mg, 0.730 mmol, 1.0 eq.), DIPEA (0.6 mL, 3.650 mmol, 5.0 eq.), 2-(diphenoxyphosphoryl)pyrrolidin-1-ium chloride (249.0 mg, 0.730 mmol, 1.0 eq.).
Column chromatography conditions: 45% ACN/Water; Yield: 56% (203 mg, colorless oil).

**UPLC:** R*t* 1.81 minutes; (ESI+) m/z calc'd for $C_{27}H_{31}N_2O_5P$ $[M+H]^+$: 495.52; found: 495.10.
**HRMS:** m/z calc'd for $C_{27}H_{31}N_2O_5P$ $[M+H]^+$: 495,2043; found: 495,2055.

##### Synthesis of Alk-(D)-Pro-Pro-DPP (Probe 2)

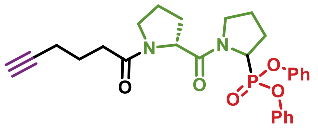


Using general procedure 2, hex-5-ynoyl-D-proline (21.0 mg, 0.100 mmol, 1.0 eq.), HATU (38.0 mg, 0.100 mmol, 1.0 eq.), DIPEA (0.09 mL, 0.500 mmol, 5.0 eq.), 2-(diphenoxyphosphoryl)pyrrolidin-1-ium chloride (34.0 mg, 0.100 mmol, 1.0 eq.).
Column chromatography conditions: 45% ACN/Water; Yield: 63% (31.0 mg, colorless oil).

**UPLC:** R*t* 1.80 minutes; (ESI+) m/z calc'd for $C_{27}H_{31}N_2O_5P$ $[M+H]^+$: 495.52; found: 495.14.
**HRMS:** m/z calc'd for $C_{27}H_{31}N_2O_5P$ $[M+H]^+$: 495,2043; found: 495,2055.

##### Synthesis of Alk-Pro-Pro-Pro-DPP (Probe 3)

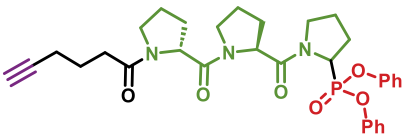

Using general procedure 2, hex-5-ynoyl-L-prolyl-L-proline (50.7 mg, 0.165 mmol, 1.0 eq.), HATU (63.0 mg, 0.165 mmol, 1.0 eq.), DIPEA (0.14 mL, 0.827 mmol, 5.0 eq.), 2-(diphenoxyphosphoryl)pyrrolidin-1-ium chloride (56 mg, 0.165 mmol, 1.0 eq.).
Column chromatography conditions: 45% ACN/Water; Yield: 52% (51.0 mg, colorless solid).

**UPLC:** R*t* 1.77 minutes; (ESI+) m/z calc'd for $C_{32}H_{38}N_3O_6P$ $[M+H]^+$: 592.64; found: 592.20.
**HRMS:** m/z calc'd for $C_{32}H_{38}N_3O_6P$ $[M+H]^+$: 592,2571; found: 592.2587.

##### Synthesis of Alk-Ala-Pro-Pro-DPP (Probe 4)

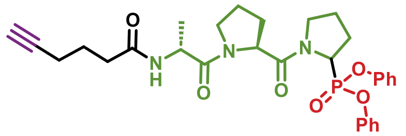


Using general procedure 2, hex-5-ynoyl-L-alanyl-L-proline (53.8 mg, 0.191 mmol, 1.0 eq.), HATU (73.0 mg, 0.191 mmol, 1.0 eq.), DIPEA (0.16 mL, 0.959 mmol, 5.0 eq.), 2-(diphenoxyphosphoryl)pyrrolidin-1-ium chloride (73.0 mg, 0.191 mmol, 1.0 eq.).
Column chromatography conditions: 50% ACN/Water; Yield: 57% (61.3 mg, colorless oil).

**UPLC:** R*t* 1.68 and 1.70 minutes; (ESI+) m/z calc'd for $C_{30}H_{36}N_3O_6P$ $[M+H]^+$: 566.60; found: 566.20 and 566.10.
**HRMS:** m/z calc'd for $C_{30}H_{36}N_3O_6P$ $[M+H]^+$: 566,2414; found: 566,2418.

##### Synthesis of Alk-Ala-Ala-Pro-Pro-DPP (Probe 5)

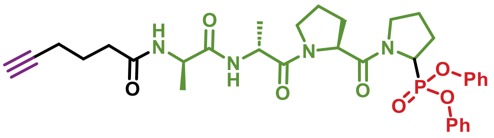


Using general procedure 2, hex-5-ynoyl-L-alanyl-L-alanyl-L-proline (41.9 mg, 0.119 mmol, 1.0 eq.), HATU (45.2 mg, 0.119 mmol, 1.0 eq.), DIPEA (0.10 mL, 0.595 mmol, 5.0 eq.), 2-(diphenoxyphosphoryl)pyrrolidin-1-ium chloride (40.6 mg, 0.119 mmol, 1.0 eq.).
Column chromatography conditions: 50% ACN/Water; Yield: 74% (56.3 mg, colorless oil).

**UPLC:** R*t* 1.67 and 1.68 minutes; (ESI+) m/z calc'd for $C_{33}H_{41}N_4O_7P$ $[M+H]^+$: 637.68; found: 637.24 and 637.09.
**HRMS:** m/z calc'd for $C_{33}H_{41}N_4O_7P$ $[M+H]^+$: 637.2786; found: 637.2805.

##### Synthesis of Alk-Ser-Pro-DPP (Probe 6)

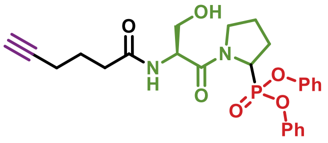


Using general procedure 2 and 3, O-(tert-butyl)-N-(hex-5-ynoyl)-L-serine (53.9 mg, 0.211 mmol, 1.0 eq.), HATU (80.2 mg, 0.211 mmol, 1.0 eq.), DIPEA (0.18 mL, 1.050 mmol, 5.0 eq.), 2-(diphenoxyphosphoryl)pyrrolidin-1-ium chloride (72.0 mg, 0.211 mmol, 1.0 eq.).
Column chromatography conditions for **t-butyl protected compound**: 50% ACN/Water; Yield: 12% (13.3 mg, colorless liquid).

Column chromatography conditions for **t-butyl deprotected compound**: 45% ACN/Water; Yield: 73% (8.5 mg, colorless solid).

**UPLC:** R*t* 1.63 minutes; (ESI+) m/z calc'd for $C_{25}H_{29}N_2O_6P$ $[M+H]^+$: 485.48; found: 485.10.
**HRMS:** m/z calc'd for $C_{25}H_{29}N_2O_6P$ $[M+H]^+$: 485.1836; found: 485.1845.

**Synthesis of Alk-(D)-Ser-Pro-DPP (Probe 7)**

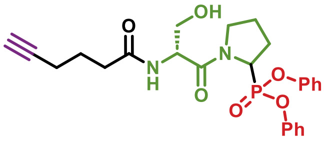


Using general procedure 2 and 3, O-(tert-butyl)-N-(hex-5-ynoyl)-D-serine (158.0 mg, 0.618 mmol, 1.0 eq.), HATU (235.0 mg, 0.618 mmol, 1.0 eq.), DIPEA (0.54 mL, 3.090 mmol, 5.0 eq.), 2-(diphenoxyphosphoryl)pyrrolidin-1-ium chloride (211.0 mg, 0.618 mmol, 1.0 eq.).
Column chromatography conditions for **t-butyl protected compound**: 50% ACN/Water; Yield: 12% (40.0 mg, colorless liquid).
Column chromatography conditions for **t-butyl deprotected compound**: 45% ACN/Water; Yield: 50% (18.0 mg, colorless solid).

**UPLC:** R*t* 1.65 minutes; (ESI+) m/z calc'd for $C_{25}H_{29}N_2O_6P$ $[M+H]^+$: 485.48; found: 485.10.
**HRMS:** m/z calc'd for $C_{25}H_{29}N_2O_6P$ $[M+H]^+$: 485.1836; found: 485.1845.

**Synthesis of Alk-Pro-Ser-Pro-DPP (Probe 8)**

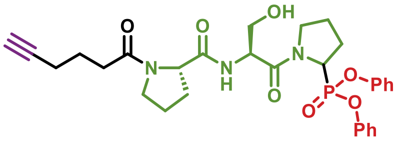


Using general procedure 2 and 3, O-(tert-butyl)-N-(hex-5-ynoyl-L-prolyl)-L-serine (215.8 mg, 0.612 mmol, 1.0 eq.), HATU (233.0 mg, 0.612 mmol, 1.0 eq.), DIPEA (0.53 mL, 3.061 mmol, 5.0 eq.), 2-(diphenoxyphosphoryl)pyrrolidin-1-ium chloride (209.0 mg, 0.612 mmol, 1.0 eq.).
Column chromatography conditions for **t-butyl protected compound**: 50% ACN/Water; Yield: 11% (43.0 mg, colorless liquid).
Column chromatography conditions for **t-butyl deprotected compound**: 35% ACN/Water; Yield: 40% (15.5 mg, colorless solid).

**UPLC:** R*t* 1.66 minutes; (ESI+) m/z calc'd for $C_{30}H_{36}N_3O_7P$ $[M+H]^+$: 582.60; found: 582.13.
**HRMS:** m/z calc'd for $C_{30}H_{36}N_3O_7P$ $[M+H]^+$: 582.2364; found: 582.2369.

**Synthesis of Alk-Thr-Pro-Ser-Pro-DPP (Probe 9)**

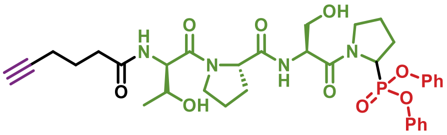


Using general procedure 2 and 3, O-(tert-butyl)-N-O-(tert-butyl)-N-(hex-5-ynoyl)-L-allothreonyl-L-prolyl-L-serine (259.7 mg, 0.509 mmol, 1.0 eq.), HATU (194.0 mg, 0.509 mmol, 1.0 eq.), DIPEA (0.44 mL, 2.540 mmol, 5.0 eq.), 2-(diphenoxyphosphoryl)pyrrolidin-1-ium chloride (174.0 mg, 0.509 mmol, 1.0 eq.).

Column chromatography conditions for **t-butyl protected compound**: 60% ACN/Water; Yield: 20% (83.0 mg, colorless liquid).
Column chromatography conditions for **t-butyl deprotected compound**: 25% ACN/Water; Yield: 15% (21.0 mg, colorless solid).

**UPLC:** R*t* 1.60 minutes; (ESI+) m/z calc'd for $C_{34}H_{43}N_4O_9P$ $[M+H]^+$: 683.71; found: 683.21.
**HRMS:** m/z calc'd for $C_{34}H_{43}N_4O_9P$ $[M+H]^+$: 683.2840; found: 683.2844.

##### Synthesis of Alk-Val-Pro-DPP (Probe 10)

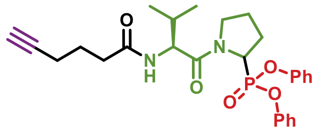


Using general procedure 2, hex-5-ynoyl-L-valine (55.0 mg, 0.260 mmol, 1.0 eq.), HATU (99.0 mg, 0.260 mmol, 1.0 eq.), DIPEA (0.22 mL, 1.303 mmol, 5.0 eq.), 2-(diphenoxyphosphoryl)pyrrolidin-1-ium chloride (88.6 mg, 0.260 mmol, 1.0 eq.).
Column chromatography conditions: 50% ACN/Water; Yield: 25% (32.0 mg, colorless oil).

**UPLC:** R*t* 1.86 and 1.88 minutes; (ESI+) m/z calc'd for $C_{27}H_{33}N_2O_5P$ $[M+H]^+$: 497.54; found: 497.10.
**HRMS:** m/z calc'd for $C_{27}H_{33}N_2O_5P$ $[M+H]^+$: 497.2200; found: 497.2213.

##### Synthesis of Alk-(D)-Val-Pro-DPP (Probe 11)

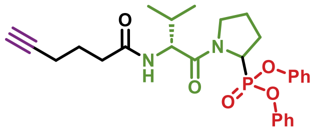


Using general procedure 2, hex-5-ynoyl-D-valine (30.0 mg, 0.142 mmol, 1.0 eq.), HATU (54.0 mg, 0.142 mmol, 1.0 eq.), DIPEA (0.12 mL, 0.710 mmol, 5.0 eq.), 2-(diphenoxyphosphoryl)pyrrolidin-1-ium chloride (48.0 mg, 0.142 mmol, 1.0 eq.).
Prep-HPLC column chromatography conditions: 78% ACN+0.1%TFA/Water+0.1%TFA; Yield: 14% (9.8 mg, colorless oil).

**UPLC:** R*t* 1.90 and 1.92 minutes; (ESI+) m/z calc'd for $C_{27}H_{33}N_2O_5P$ $[M+H]^+$: 497.54; found: 497.10 and 497.20.
**HRMS:** m/z calc'd for $C_{27}H_{33}N_2O_5P$ $[M+H]^+$: 497.2200; found: 497.2217.

##### Synthesis of Alk-Pro-Pro-TFEP (Probe 12)

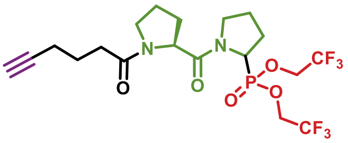


Using general procedure 4, Alk-Pro-Pro-DPP (173.0 mg, 0.349 mmol, 1.0 eq.), 2,2,2-trifluoroethanol (4 mL), potassium fluoride (203.0 mg, 3.498 mmol, 10.0 eq.), and 18-crown-6 (7 mg, 0.023 mmol, 0.068 eq.).
Prep-HPLC column chromatography conditions: 60% ACN + 0.1% TFA/Water + 0.1% TFA; Yield: 8% (15.0 mg, colorless oil).

**UPLC:** R*t* 1.74 minutes; (ESI+) m/z calc'd for $C_{19}H_{25}F_6N_2O_5P$ $[M+H]^+$: 507.38; found: 507.10.
**HRMS:** m/z calc'd for $C_{19}H_{25}F_6N_2O_5P$ $[M+H]^+$: 507.1478; found: 507.1487.

##### Synthesis of Alk-Ala-Pro-Pro-TFEP (Probe 13)

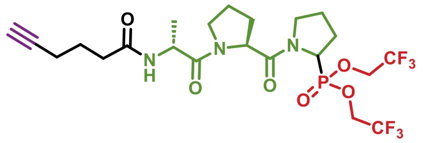

Using general procedure 4, Alk-Pro-Pro-DPP (55.0 mg, 0.093 mmol, 1.0 eq.), 2,2,2-trifluoroethanol (2 mL), potassium fluoride (56.3 mg, 0.93 mmol, 10.0 eq.), and 18-crown-6 (1.74 mg, 0.0063 mmol, 0.068 eq.).
Column chromatography conditions: 55% ACN + 0.1% TFA/Water + 0.1% TFA; Yield: 17% (9.5 mg, colorless oil).

**UPLC:** R*t* 1.66 minutes; (ESI+) m/z calc'd for $C_{22}H_{30}F_6N_3O_6P$ $[M+H]^+$: 578.46; found: 578.10.
**HRMS:** m/z calc'd for $C_{22}H_{30}F_6N_3O_6P$ $[M+H]^+$: 578.1849; found: 578.1849.

##### Synthesis of Alk-Pro-Pro-P(OEt)(OPh) (Probe 14) – 3 synthetic steps

Step1: Synthesis of Alk-Pro-Pro-P(OEt)$_2$

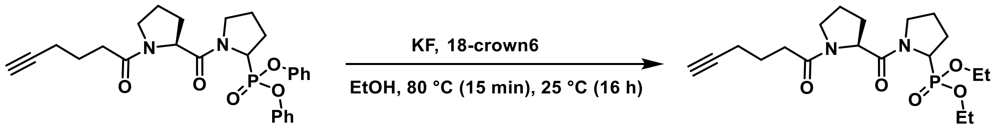


In a round bottom flask, diphenyl (1-(hex-5-ynoyl-L-prolyl)pyrrolidin-2-yl)phosphonate (150.0 mg, 0.303 mmol, 1.0 eq.) was dissolved in ethanol (3 mL). Potassium fluoride (176.0 mg, 3.033 mmol, 10.0 eq.) and 18-crown-6 (8.0 mg, 0.030 mmol, 0.1 eq.) were added, and the reaction mixture was stirred at 80 °C for 15 minutes and then at room temperature for 16 hours. The mixture was concentrated under reduced pressure, and the residue was purified by reverse phase column chromatography (15% ACN/water) to afford the title compound as a colorless oil (83.0 mg, 69%).

**UPLC:** R*t* 1.47 minutes; (ESI+) m/z calc'd for $C_{19}H_{31}N_2O_5P$ $[M+H]^+$: 399.43; found: 399.20.

Step 2: Synthesis of Alk-Pro-Pro-P(OEt)(OLi)

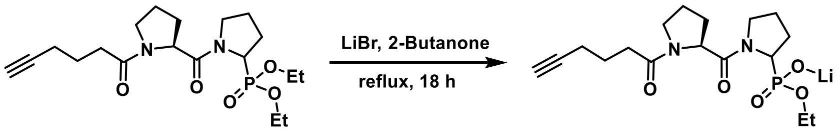


Lithium bromide (66.6 mg, 0.767 mmol, 2.5 eq.) was added to a solution of Alk-Pro-Pro-P(OEt)$_2$ (122.0 mg, 0.307 mmol, 1.0 eq.) in 2-butanone (3 mL), and the reaction mixture was stirred at 85 °C for 20 hours. A white precipitate formed and was removed by filtration. The solid was washed with 2-butanone (4 x 2 mL) and the combined filtrate was concentrated under reduced pressure to afford the title compound as a colorless solid (99 mg, 85%)

**UPLC:** R*t* 1.04 minutes; (ESI-) m/z calc'd for $C_{17}H_{27}N_2O_5P$ $[M-H]^-$: 369.38; found: 369.20.

Step 3: Synthesis of Alk-Pro-Pro-P(OEt)(OPh) **(Probe 14)**

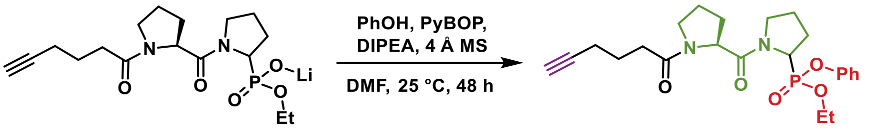


Under inert atmosphere, in a dry round bottom flask containing 4Å MS, Alk-Pro-Pro-P(OEt)(OLi) (78.0 mg, 0.208 mmol, 1.0 eq.), phenol (30.0 mg, 0.312 mmol, 1.5 eq.), and PyBOP (162.0 mg, 0.312 mmol, 1.5 eq.) were dissolved in DMF (2 mL). DIPEA (0.014 mL, 0.832 mmol, 4.0 eq.) was added and the reaction mixture was stirred for 48 hours. Upon completion, the reaction mixture was concentrated under reduced pressure and the crude mixture was subjected to reverse phase column chromatography to afford the title compound.

Column chromatography conditions: 37% ACN/Water; Yield: 5% (5.2 mg, colorless oil).

**UPLC:** R*t* 1.61 minutes; (ESI+) m/z calc'd for $C_{23}H_{31}N_2O_5P$ $[M+H]^+$: 447.48; found: 447.17.
**HRMS:** m/z calc'd for $C_{23}H_{31}N_2O_5P$ $[M+H]^+$: 447.2043; found: 447.2053.

##### Synthesis of SulfoCy5-Pro-Pro-DPP (Probe 15)

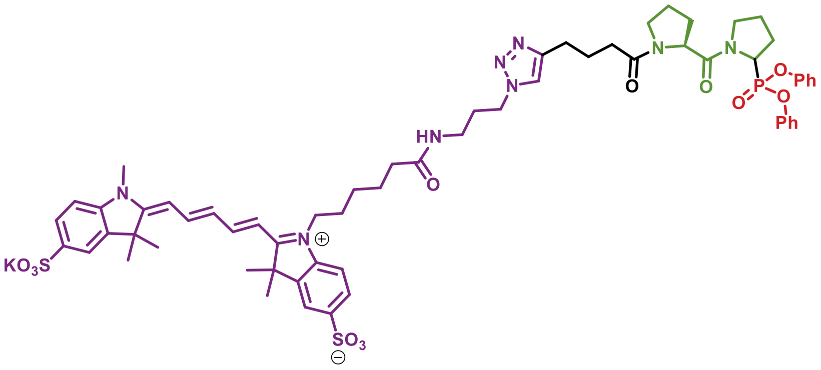

Using general procedure 5, diphenyl (1-(hex-5-ynoyl-L-prolyl)pyrrolidin-2-yl)phosphonate (12.4 mg, 25.2 μmol, 1 eq.), Sulfo-Cyanine 5 azide (25.0 mg, 32.7 μmol, 1.3 eq.), THF/$H_2O$ (1.5 mL/1 mL), sodium (R)-2-((S)-1,2-dihydroxyethyl)-4-hydroxy-5-oxo-2,5-dihydrofuran-3-olate (13.0 mg, 65.6 μmol, 2.6 eq.), and $CuSO_4.5H_2O$ (8.0 mg, 32.7 μmol, 1.3 eq.).
Column chromatography conditions: 25% ACN /Water; Yield: 71% (22.6 mg, dark blue solid).

**Note:** Both UPLC and HRMS analyses showed the ion corresponding to the free acid $C_{62}H_{75}N_8O_{12}PS_2$, rather than the potassium salt $C_{62}H_{74}KN_8O_{12}PS_2$
**UPLC:** R*t* 1.59 minutes; (ESI+) m/z calc'd for $C_{62}H_{75}N_8O_{12}PS_2$ $[M+2H]^{2+}$: 610.24; found: 610.26.
**HRMS:** m/z calc'd for $C_{62}H_{75}N_8O_{12}PS_2$ $[M+H]^+$: 1219.4756; found: 1219.4758.

##### Synthesis of Cy5-Pro-Pro-DPP (Probe 16)

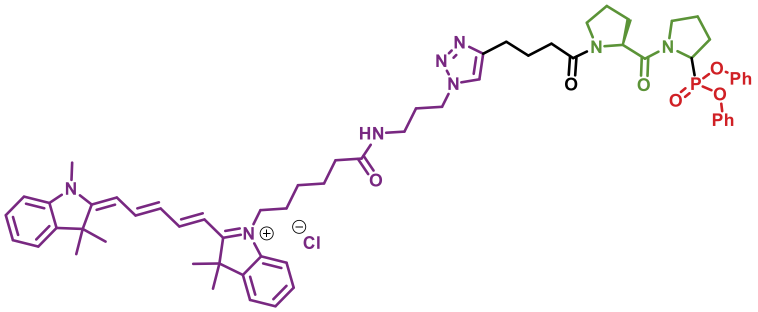

Using general procedure 5, diphenyl (1-(hex-5-ynoyl-L-prolyl)pyrrolidin-2-yl)phosphonate (19.0 mg, 38.3 μmol, 1 eq.), Cyanine 5 azide (30.0 mg, 49.8 μmol, 1.3 eq.), THF/$H_2O$ (1.5 mL/1 mL), sodium (R)-2-((S)-1,2-dihydroxyethyl)-4-hydroxy-5-oxo-2,5-dihydrofuran-3-olate (20.0 mg, 99.5 μmol, 2.6 eq.), and $CuSO_4 \cdot 5H_2O$ (12.4 mg, 49.8 μmol, 1.3 eq.).
Column chromatography conditions: 45% ACN /Water; Yield: 40% (16.8 mg, dark blue solid)

**UPLC:** R*t* 1.86 minutes; (ESI+) m/z calc'd for $C_{62}H_{76}N_8O_6P$ $[M+2H]^{2+}$: 530.28; found: 530.42.
**HRMS:** m/z calc'd for $C_{62}H_{76}N_8O_6P$ $[M+H]^+$: 1059.5620; found: 1059.5615.

##### Synthesis of TAMRA-Pro-Pro-DPP (Probe 17)

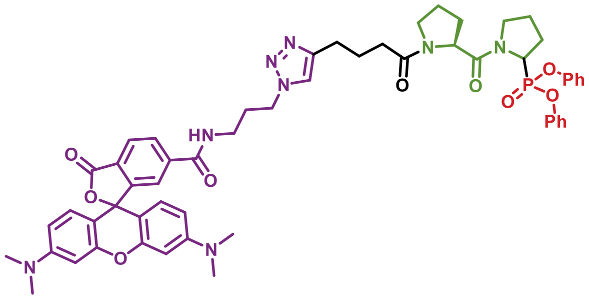

Using general procedure 5, diphenyl (1-(hex-5-ynoyl-L-prolyl)pyrrolidin-2-yl)phosphonate (75.0 mg, 0.151 mmol, 1 eq.), TAMRA azide, 5 isomer (101.0 mg, 0.197 mmol, 1.3 eq.), THF/$H_2O$ (3 mL/2 mL), sodium (R)-2-((S)-1,2-dihydroxyethyl)-4-hydroxy-5-oxo-2,5-dihydrofuran-3-olate (78.5 mg, 0.396 mmol, 2.6 eq.), and $CuSO_4 \cdot 5H_2O$ (49.4 mg, 0.198 mmol, 1.3 eq.).
Column chromatography conditions: 35% ACN /Water; Yield: 48% (73.0 mg, deep violet solid).

**UPLC:** R*t* 1.66 minutes; (ESI+) m/z calc'd for $C_{55}H_{59}N_8O_9P$ $[M+2H]^{2+}$: 504.55; found: 504.40.
**HRMS:** m/z calc'd for $C_{55}H_{59}N_8O_9P$ $[M+H]^+$: 1007.4215; found: 1007.4217.

##### Synthesis of Cbz-Pro-Pro-DPP (Compound 1)

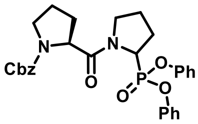

Using general procedure 2, (tert-butoxycarbonyl)proline (70.0 mg, 0.280 mmol, 1.0 eq.), HATU (106.4 mg, 0.280 mmol, 1.0 eq.), DIPEA (0.24 mL, 1.400 mmol, 5.0 eq.), 2-(diphenoxyphosphoryl)pyrrolidin-1-ium chloride (96.0 mg, 0.280 mmol, 1.0 eq.).
Column chromatography conditions: 57% ACN /Water; Yield: 71% (107.0 mg, colorless oil).

**UPLC:** R*t* 2.01 minutes; (ESI+) m/z calc'd for $C_{29}H_{31}N_2O_6P$ $[M+H]^+$: 535.54; found: 535.11.
**HRMS:** m/z calc'd for $C_{29}H_{31}N_2O_6P$ $[M+H]^+$: 535.1992; found: 535.2001.
**$^1$H NMR** ($CDCl_3$, 400 MHz): $\delta$ 7.29 – 6.95 (m, 15H), 5.34 – 4.36 (m, 4H), 3.93 – 3.27 (m, 4H), 2.53 – 1.59 (m, 8H).

##### Synthesis of Boc-Pro-Pro-DPP (Compound 2)

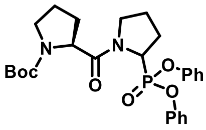

Using general procedure 2, (tert-butoxycarbonyl)proline (50.0 mg, 0.232 mmol, 1.0 eq.), HATU (88.0 mg, 0.232 mmol, 1.0 eq.), DIPEA (0.20 mL, 1.160 mmol, 5.0 eq.), 2-(diphenoxyphosphoryl)pyrrolidin-1-ium chloride (79.0 mg, 0.232 mmol, 1.0 eq.).
Column chromatography conditions: 50% ACN /Water; Yield: 64% (75.0 mg, colorless oil).

**UPLC:** R*t* 1.97 minutes; (ESI+) m/z calc'd for $C_{26}H_{33}N_2O_6P$ $[M+H]^+$: 501.53; found: 501.19.
**HRMS:** m/z calc'd for $C_{26}H_{33}N_2O_6P$ $[M+H]^+$: 501.2149; found: 501.2158.
**$^1$H NMR** ($CDCl_3$, 400 MHz): $\delta$ 7.28 – 6.92 (m, 10H), 5.41 – 4.34 (m, 2H), 3.88 – 3.26 (m, 4H), 2.51 – 2.34 (m, 2H), 2.18 – 1.65 (m, 6H), 1.38 – 1.24 (m, 9H).

**Synthesis of H-Pro-Pro-DPP (Compound 3)**

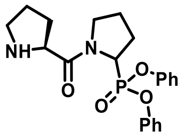


A mixture of Boc-Pro-Pro-DPP (40.0 mg, 0.079 mmol) and 4N HCl in dioxane (2 mL) was stirred at room temperature for 2 hours. The solvent was removed under reduced pressure and the resulting crude mixture was purified using reverse phase column chromatography to afford the title compound as a colorless oil.
Column chromatography conditions: 18% ACN/Water, Yield: 89% (28.3 mg, colorless oil).

**UPLC:** R*t* 1.29 minutes; (ESI+) m/z calc'd for $C_{21}H_{25}N_2O_4P$ $[M+H]^+$: 401.41; found: 401.06.
**HRMS:** m/z calc'd for $C_{21}H_{25}N_2O_4P$ $[M+H]^+$: 401.1625; found: 401.1632.
**$^1$H NMR** ($CDCl_3$, 400 MHz): $\delta$ 7.27 – 6.94 (m, 10H), 4.94 – 4.61 (m, 2H), 3.80 – 3.22 (m, 5H), 2.48 –1.59 (m, 8H).

**Synthesis of $N_3$-Pro(N-Cbz)-Pro-DPP (Compound 4)**

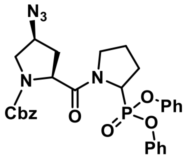


Using general procedure 2, (2S,4S)-4-azido-1-((benzyloxy)carbonyl)pyrrolidine-2-carboxylic acid (58.0 mg, 0.199 mmol, 1.0 eq.), HATU (75.6 mg, 0.199 mmol, 1.0 eq.), DIPEA (0.17 mL, 0.995 mmol, 5.0 eq.), 2-(diphenoxyphosphoryl)pyrrolidin-1-ium chloride (67.8 mg, 0.199 mmol, 1.0 eq.).
Column chromatography conditions: 49% ACN /Water; Yield: 70% (80.2 mg, colorless oil).

**UPLC:** R*t* 2.03 and 2.07 minutes; (ESI+) m/z calc'd for $C_{29}H_{30}N_5O_6P$ $[M+H]^+$: 576.56; found: 576.28.
**HRMS:** m/z calc'd for $C_{29}H_{30}N_5O_6P$ $[M+H]^+$: 576.2006; found: 576.2009.
**$^1$H NMR** ($CDCl_3$, 400 MHz): $\delta$ 7.31 – 6.92 (m, 15H), 5.22 – 4.28 (m, 5H), 3.96 – 3.82 (m, 2H), 3.56 –3.30 (m, 2H), 2.57 –1.63 (m, 6H).

**Synthesis of $N_3$-Pro(N-Boc)-Pro-DPP (Compound 5)**

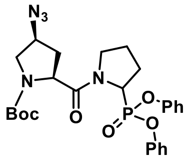


Using general procedure 2, (2S,4S)-4-Azido-1-(tert-butoxycarbonyl)pyrrolidine-2-carboxylic acid (100.0 mg, 0.344 mmol, 1.0 eq.), HATU (130.8 mg, 0.344 mmol, 1.0 eq.), DIPEA (0.30 mL, 1.720 mmol, 5.0 eq.), 2-(diphenoxyphosphoryl)pyrrolidin-1-ium chloride (117.4 mg, 0.344 mmol, 1.0 eq.).
Column chromatography conditions: 46% ACN /Water; Yield: 57% (106.0 mg, colorless oil).

**UPLC:** R*t* 2.01 and 2.05 minutes; (ESI+) m/z calc'd for $C_{29}H_{32}N_5O_6P$ $[M+H]^+$: 542.54; found: 542.31.
**HRMS:** m/z calc'd for $C_{29}H_{32}N_5O_6P$ $[M+H]^+$: 542.2163; found: 542.2167.
**$^1$H NMR** ($CDCl_3$, 400 MHz): $\delta$ 7.28 – 7.05 (m, 10H), 5.29 – 4.91 (m, 1 H), 4.72 – 4.35 (m, 1H), 3.92 – 3.74 (m, 2 H), 3.56 – 3.25 (m, 2 H), 2.55 – 2.29 (m, 3H), 2.22 – 1.85 (m, 3 H), 1.66 – 1.61 (m, 1H), 1.38 – 1.24 (m, 9H).

**Synthesis of $N_3$-Pro-Pro-DPP (UAMC-936-1)**

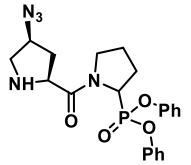


A mixture of Boc-Pro-Pro-DPP (85.0 mg, 0.157 mmol) and 4N HCl in dioxane (2.5 mL) was stirred at room temperature for 2 hours. The solvent was removed under reduced pressure and the resulting crude mixture was purified using reverse phase column chromatography to afford the title compound as a colorless oil.
Column chromatography conditions: 39% ACN /Water; Yield: 67% (46.5 mg, colorless oil).

**UPLC:** R*t* 1.27 and 1.32 minutes; (ESI+) m/z calc'd for $C_{21}H_{24}N_5O_4P$ $[M+H]^+$: 442.42; found: 442.28.
**HRMS:** m/z calc'd for $C_{21}H_{24}N_5O_4P$ $[M+H]^+$: 442.1639; found: 442.1639.
**$^1$H NMR** ($CDCl_3$, 400 MHz): $\delta$ 7.26 –7.06 (m, 10H), 5.01 – 4.92 (m, 1H), 4.10 – 4.03 (m, 1H), 3.90 – 3.76 (m, 1H), 3.67 – 3.43 (m, 2H), 3.16 – 3.07 (m, 2H), 2.92 – 2.83 (m, 1H), 2.53 – 2.28 (m, 3H), 2.20 – 1.94 (m, 2H), 1.79 – 1.67 (m, 1H).

### 6. LC-MS chromatograms

UPLC-MS chromatogram and spectrum of **probe 1:**

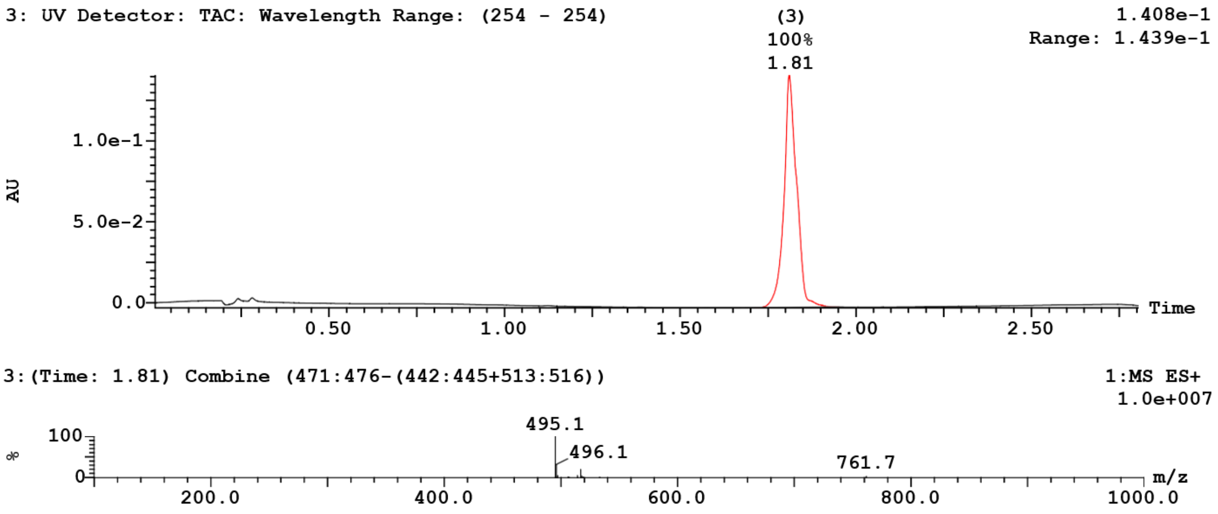


UPLC-MS chromatogram and spectrum of **probe 2:**
(*a split shoulder peak is observed in the chromatogram due to the rotamer effect)*

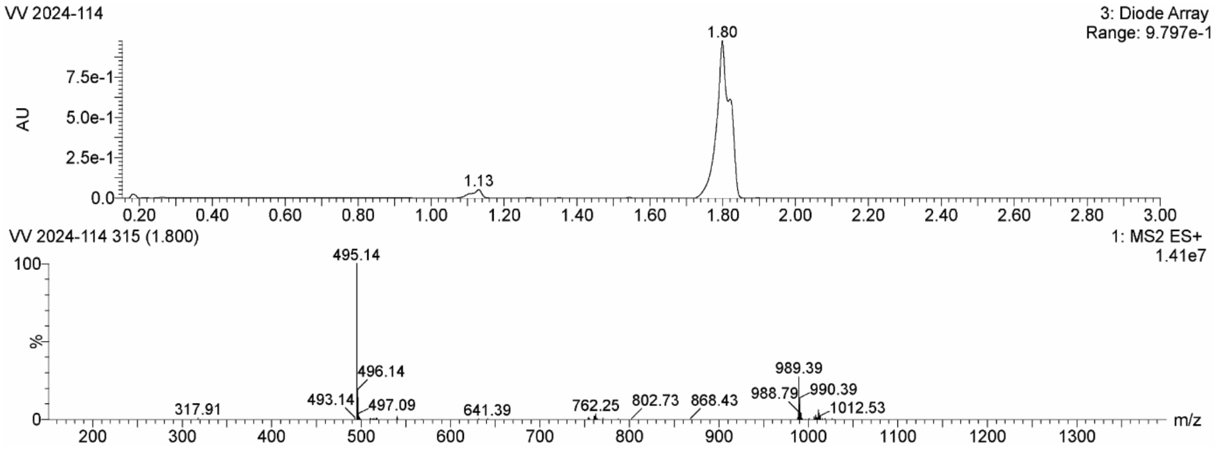

UPLC-MS chromatogram and spectrum of **probe 3**:

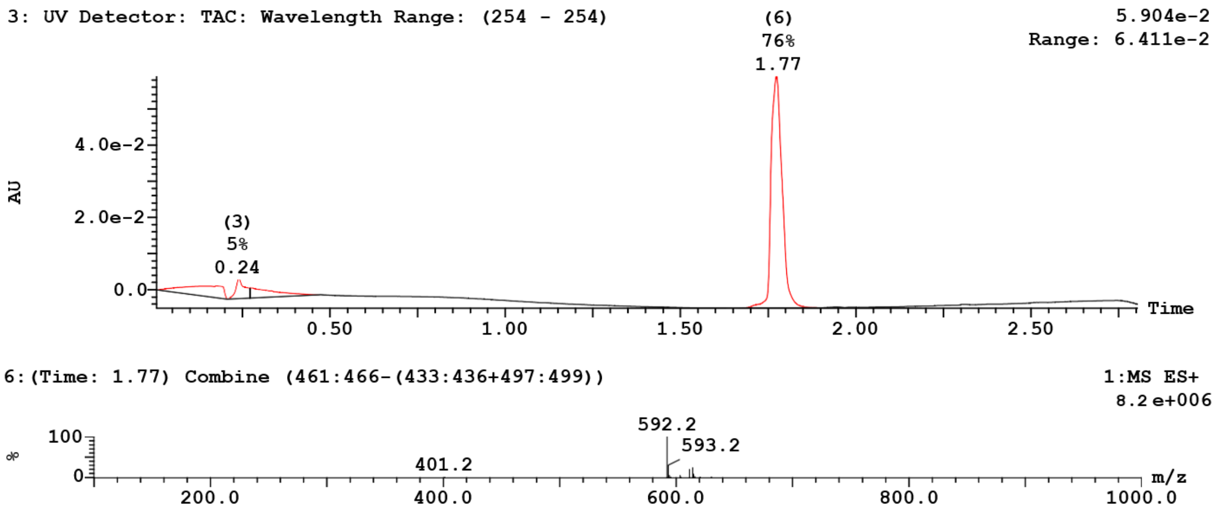


UPLC-MS chromatogram and spectrum of **probe 4**:

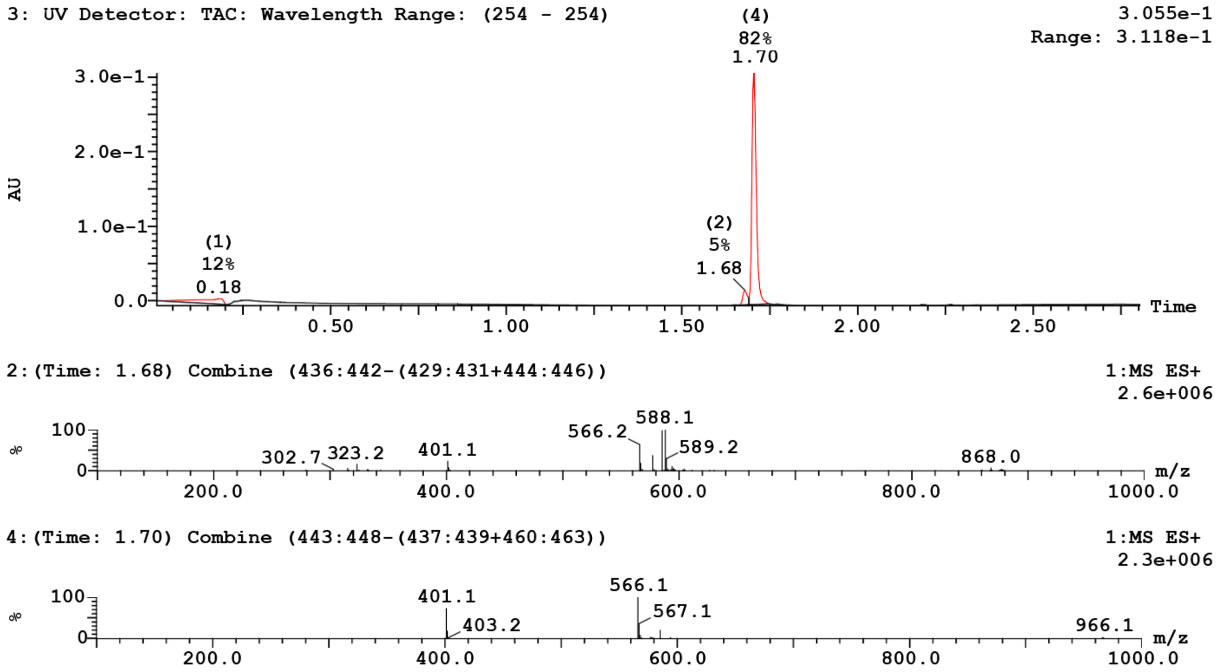

UPLC-MS chromatogram and spectrum of **probe 5**:
*(The split shoulder peak is attributed to slow interconversion of amide rotamers, likely associated with the proline-containing peptide bond)*

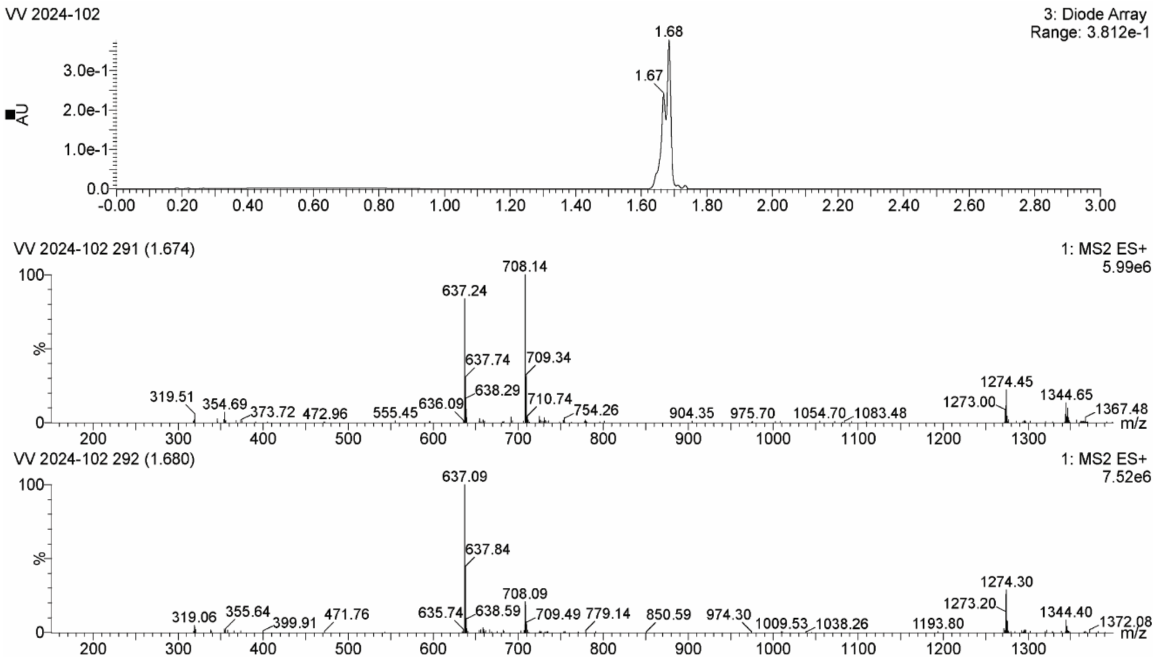


UPLC-MS chromatogram and spectrum of **probe 6**:

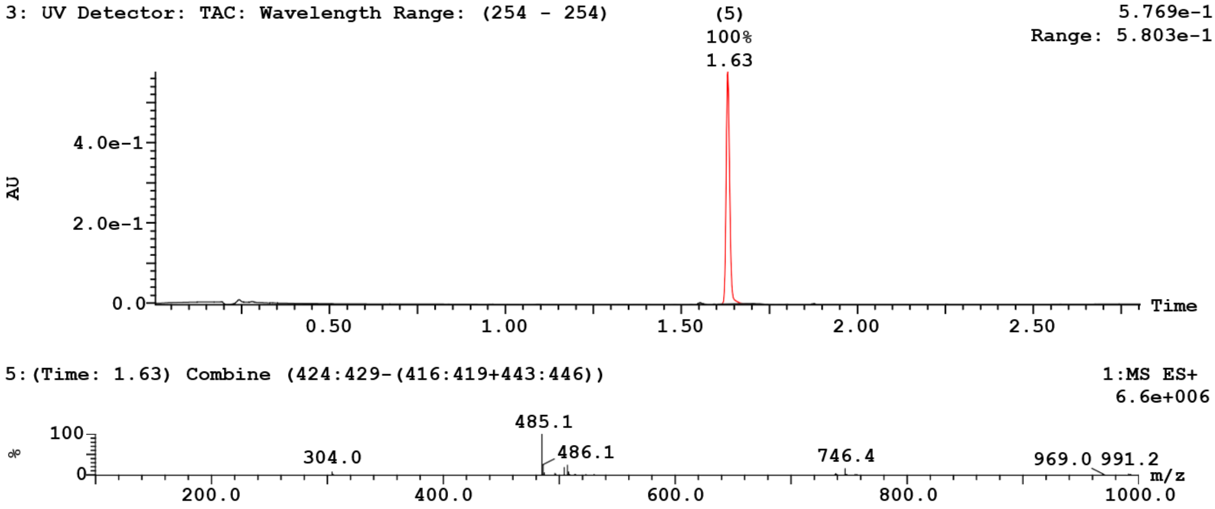

UPLC-MS chromatogram and spectrum of **probe 7**:

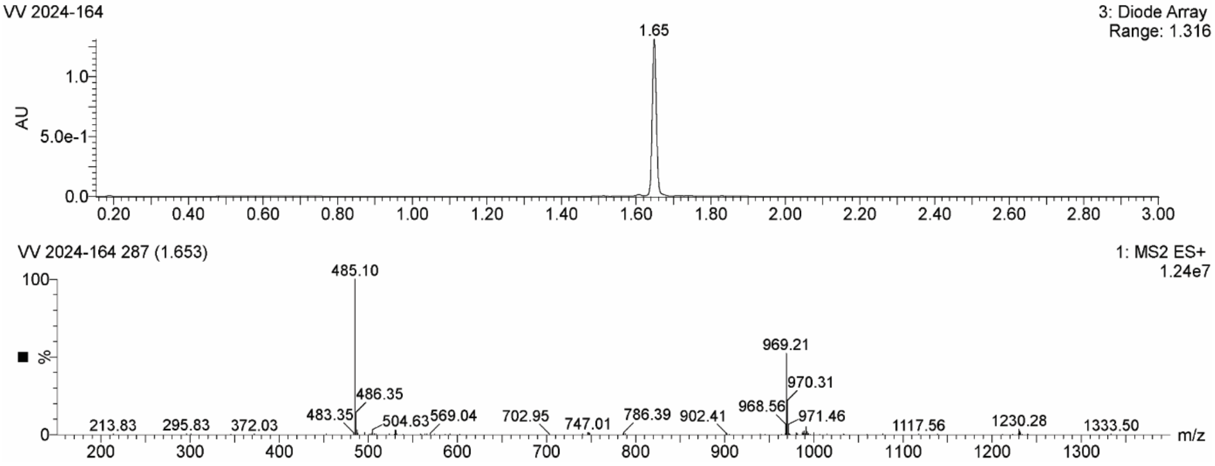


UPLC-MS chromatogram and spectrum of **probe 8**:

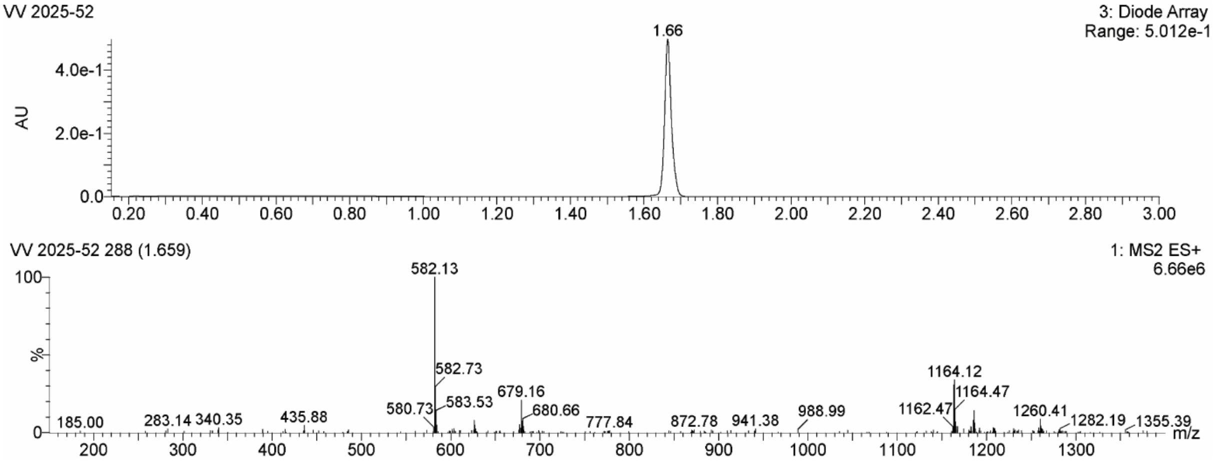

UPLC-MS chromatogram and spectrum of **probe 9**:

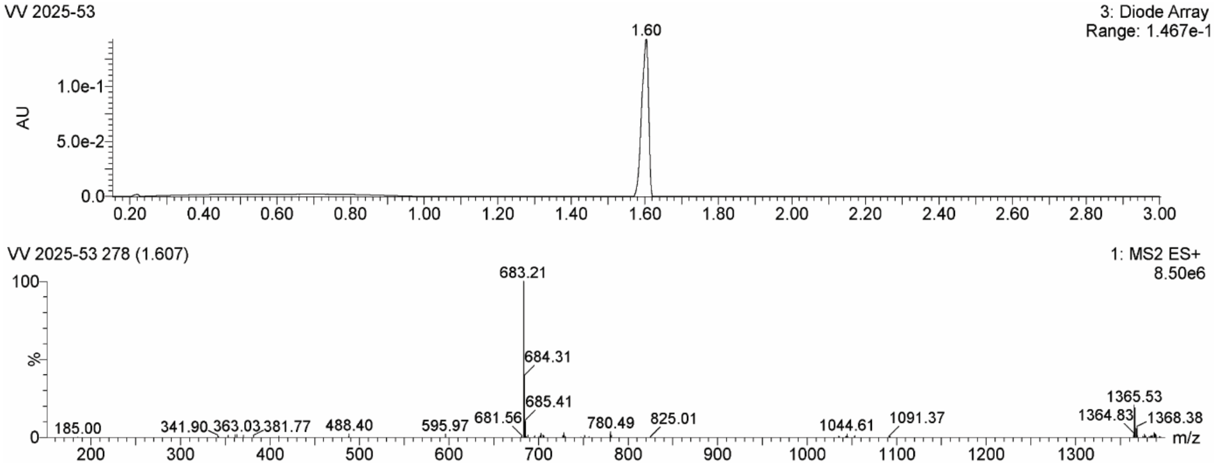


UPLC-MS chromatogram and spectrum of **probe 10**:

*(The split shoulder peak is attributed to slow interconversion of amide rotamers, likely associated with the proline-containing peptide bond)*

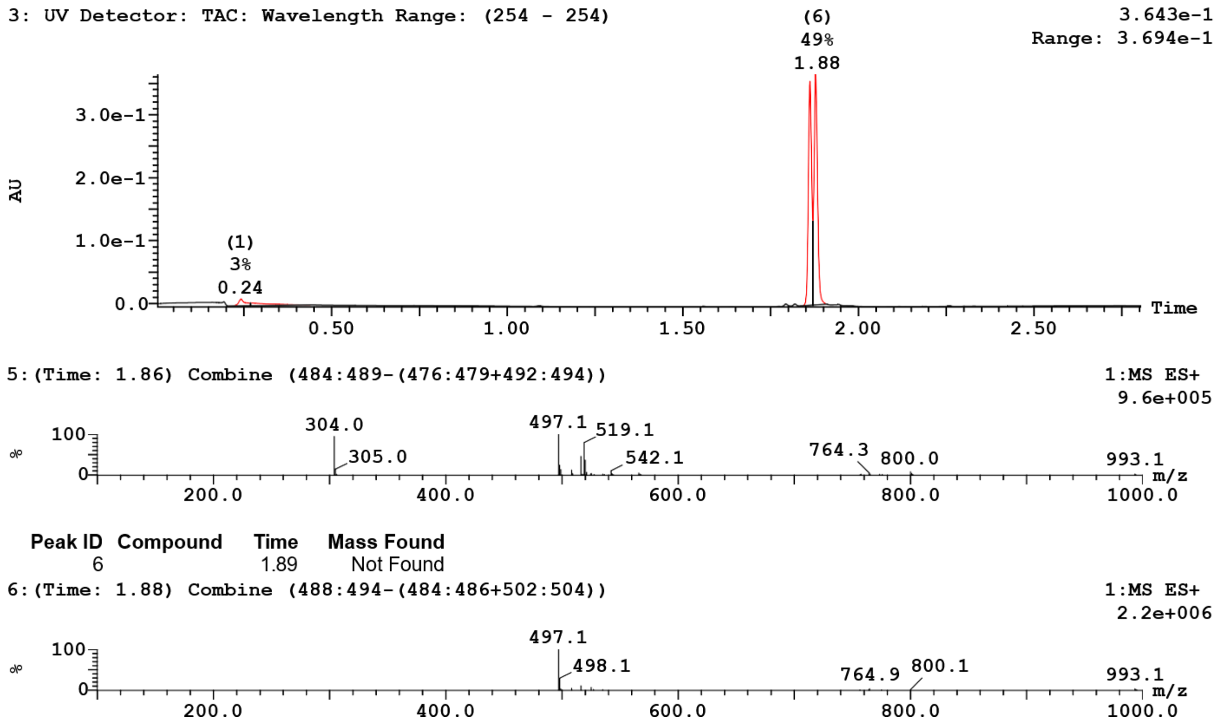

UPLC-MS chromatogram and spectrum of **probe 11**:
(*The split shoulder peak is attributed to slow interconversion of amide rotamers, likely associated with the proline-containing peptide bond)*

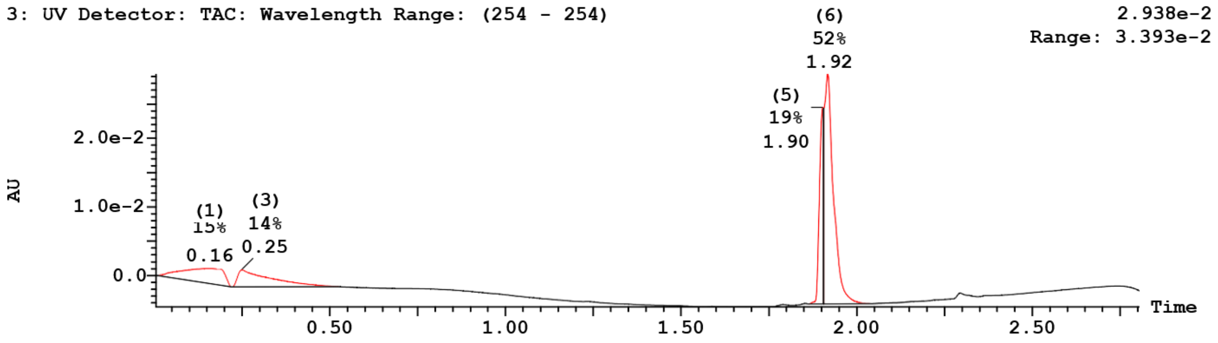


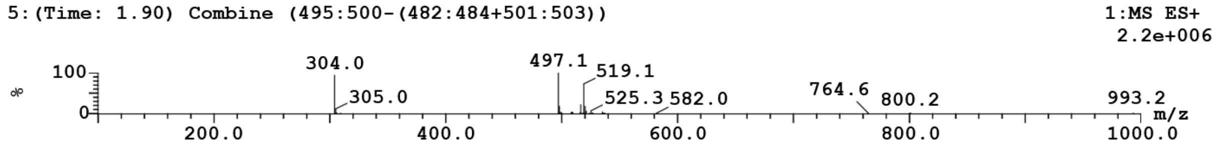


| Peak ID | Compound | Time | Mass Found |
|---|---|---|---|
| 6 | | 1.92 | Not Found |

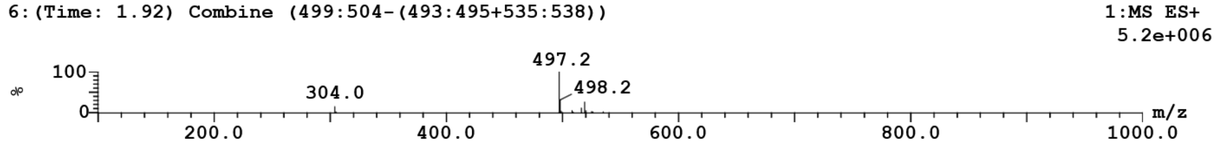


UPLC-MS chromatogram and spectrum of **probe 12**:

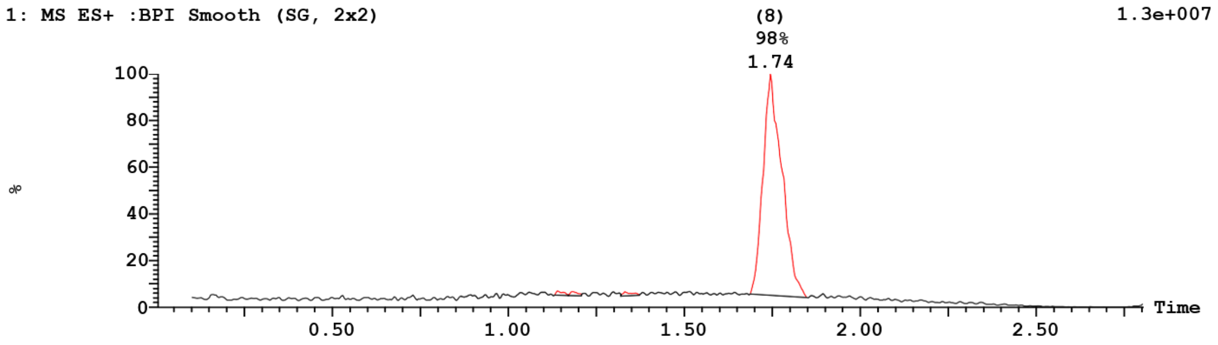


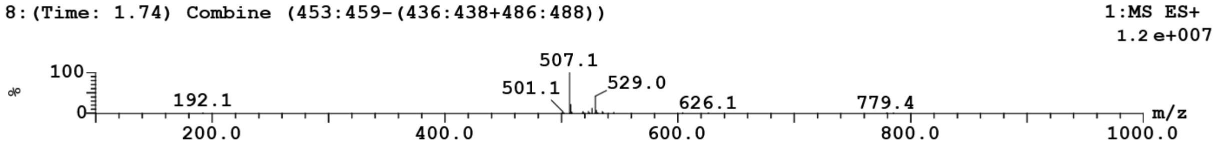

UPLC-MS chromatogram and spectrum of **probe 13**:

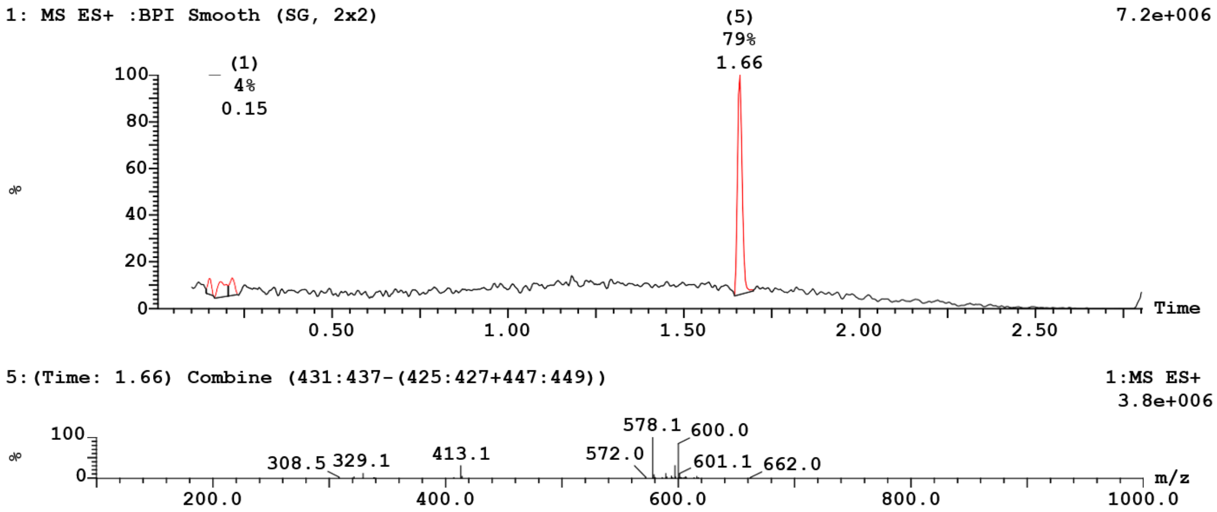


UPLC-MS chromatogram and spectrum of **Alk-Pro-Pro-P(OEt)$_2$**:

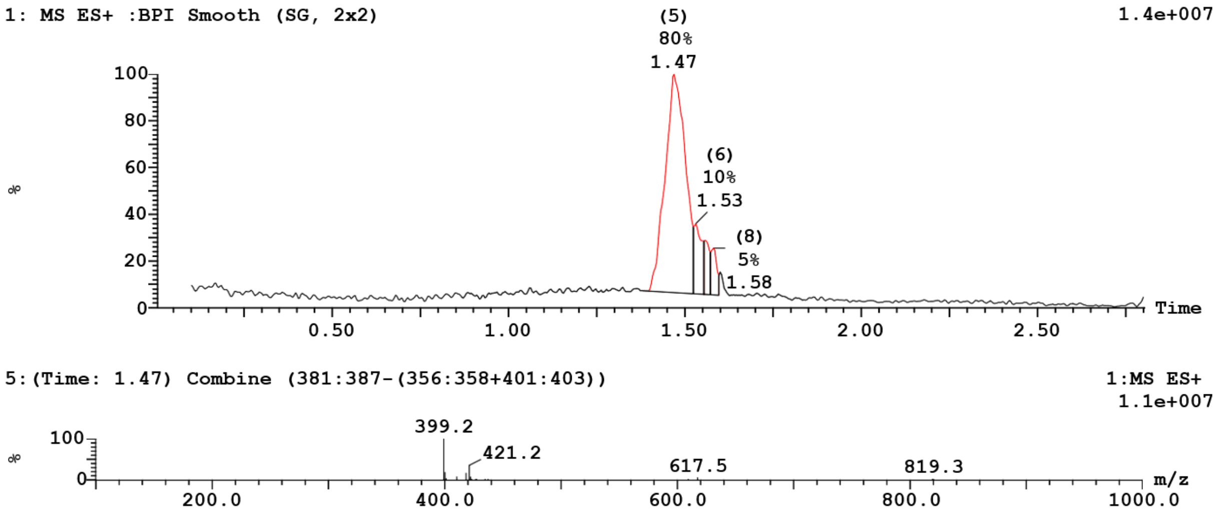

UPLC-MS chromatogram and spectrum of **Alk-Pro-Pro-P(OEt)(OLi)**:

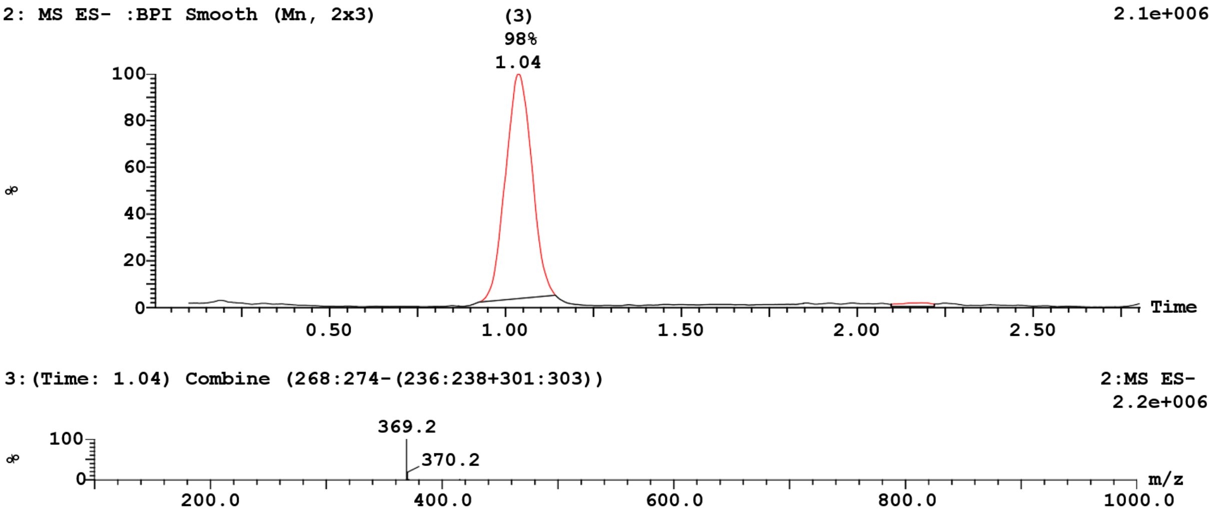


UPLC-MS chromatogram and spectrum of **probe 14**:

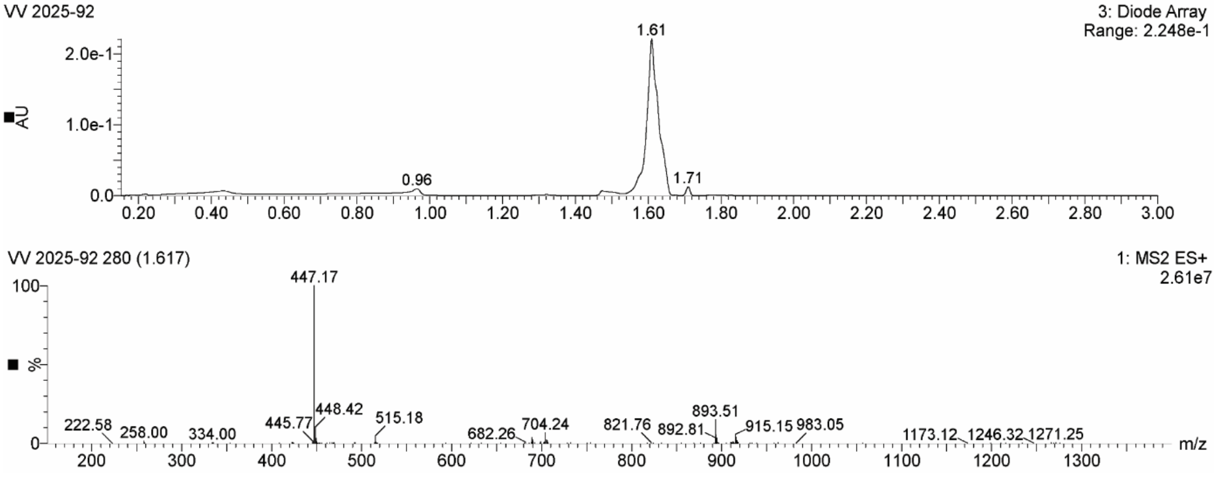

UPLC-MS chromatogram and spectrum of **probe 15**:

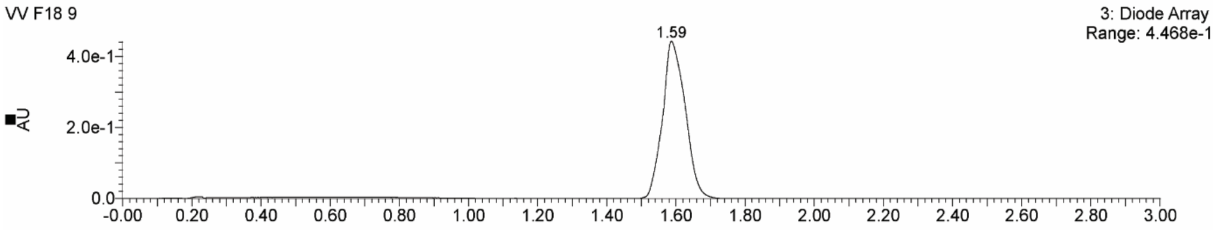


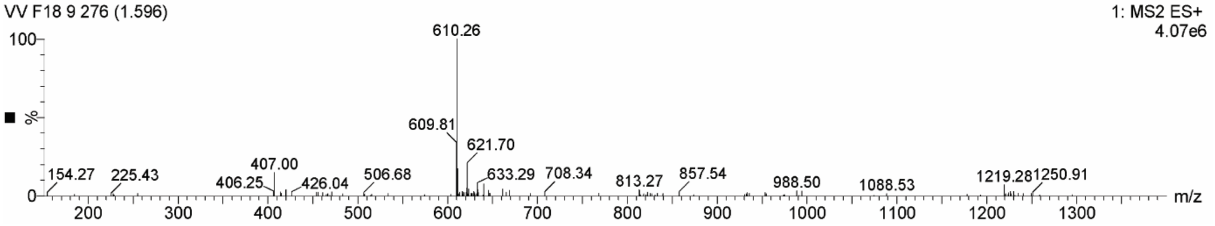


UPLC-MS chromatogram and spectrum of **probe 16:**

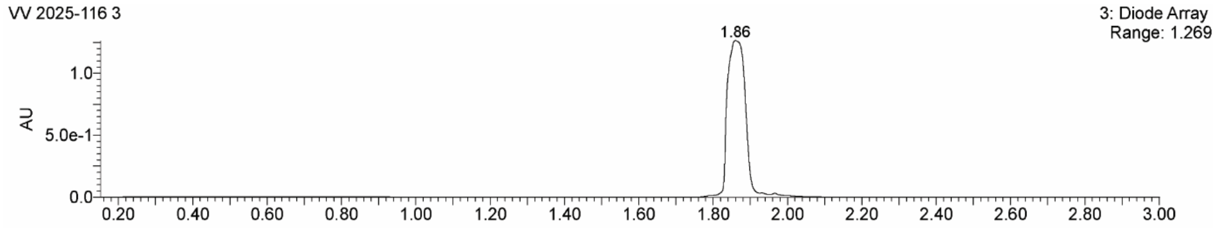


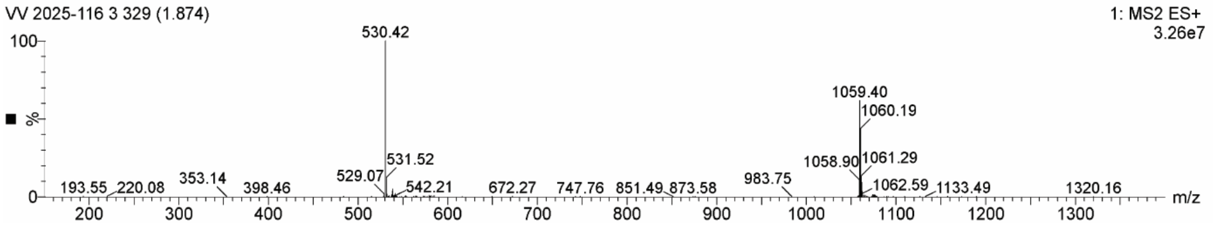

UPLC-MS chromatogram and spectrum of **probe 17**:

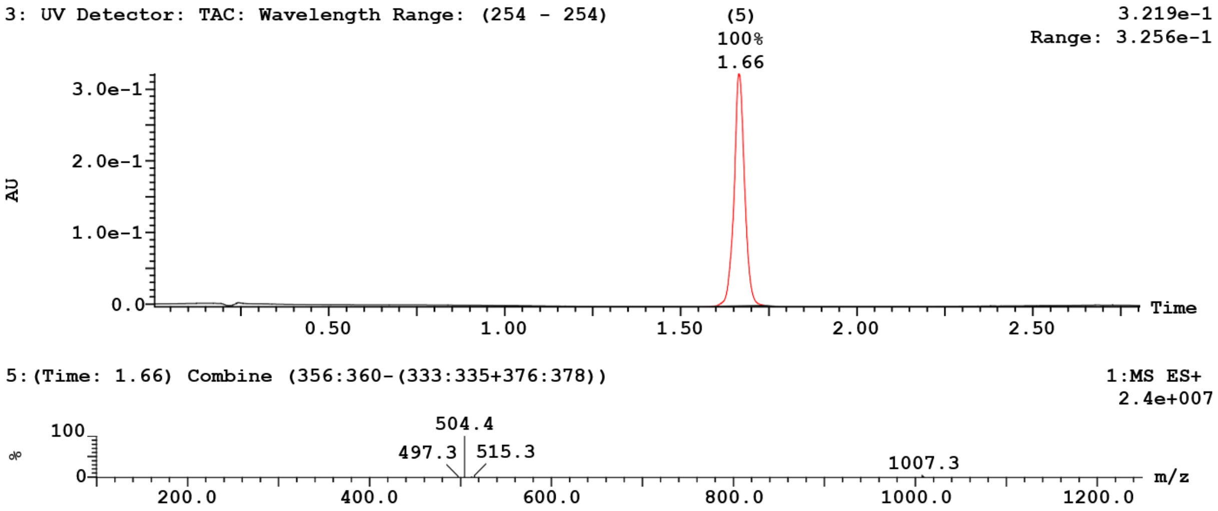


UPLC-MS chromatogram and spectrum of **compound 1**:

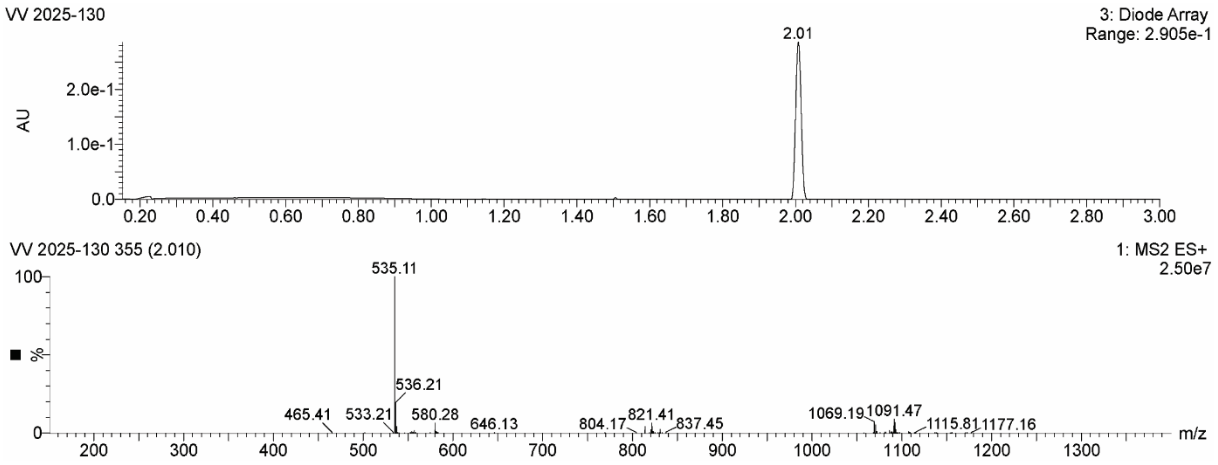

UPLC-MS chromatogram and spectrum of **compound 2**:

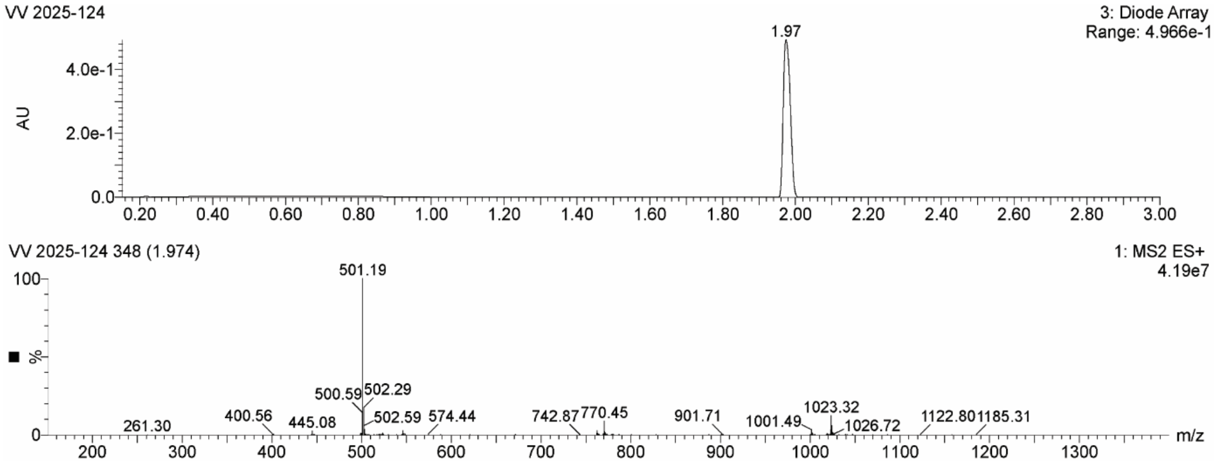


UPLC-MS chromatogram and spectrum of **compound 3**:

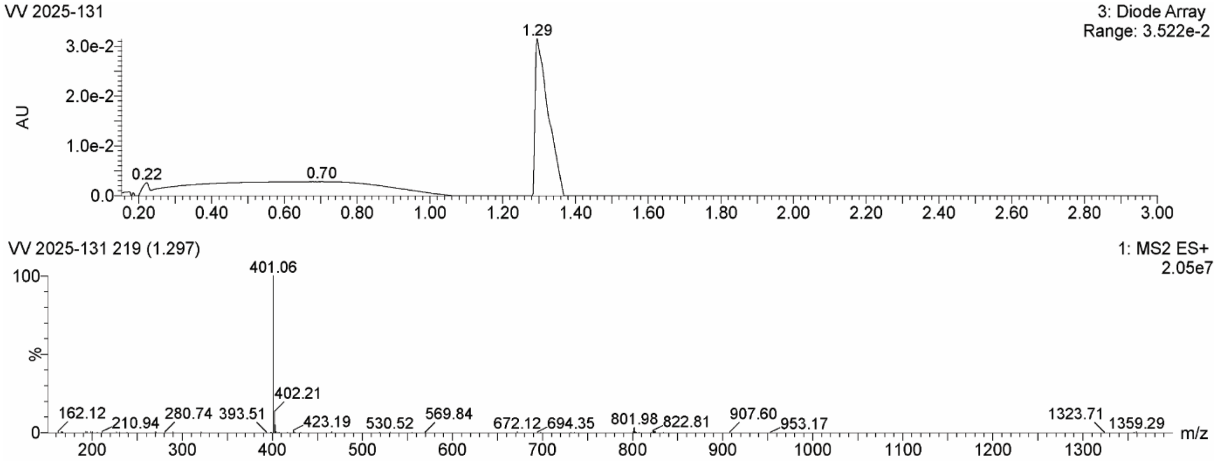

UPLC-MS chromatogram and spectrum of **compound 4**:
(*2 peaks are observed in the chromatogram since the reactions forms 2 diastereomers)*

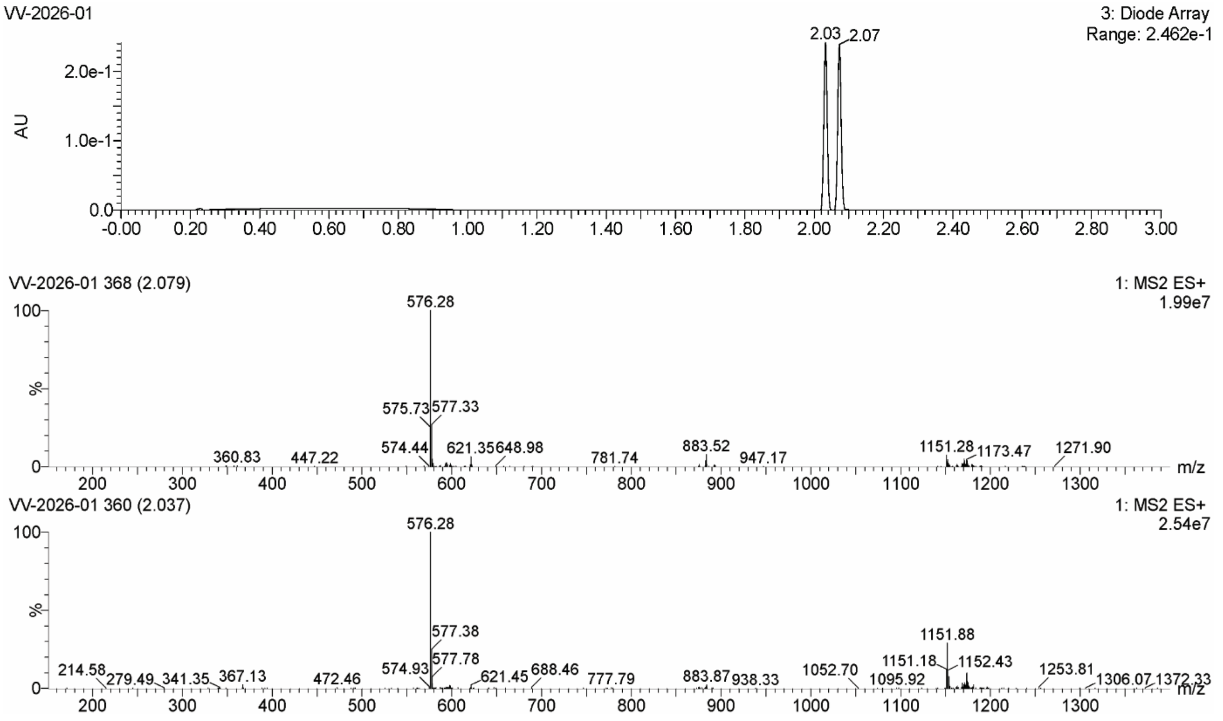


UPLC-MS chromatogram and spectrum of **compound 5**:
(*2 peaks are observed in the chromatogram since the reactions forms 2 diastereomers)*

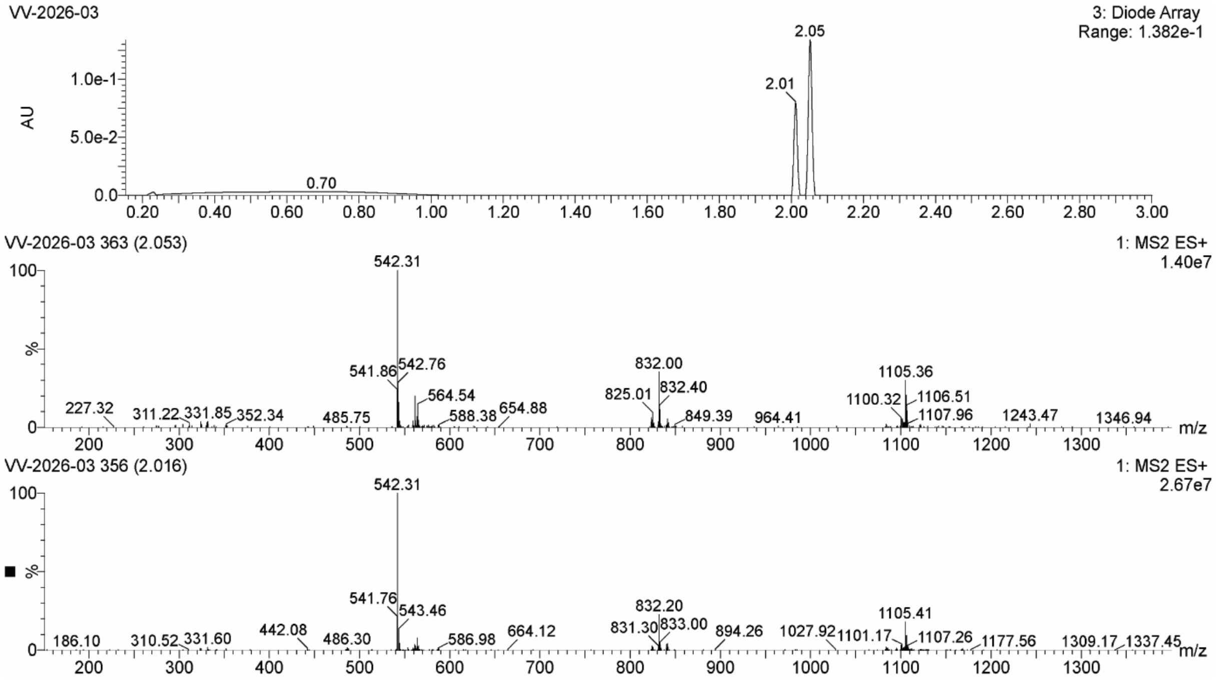

UPLC-MS chromatogram and spectrum of **UAMC-936-1**:
(*2 peaks are observed in the chromatogram since the reactions forms 2 diastereomers)*

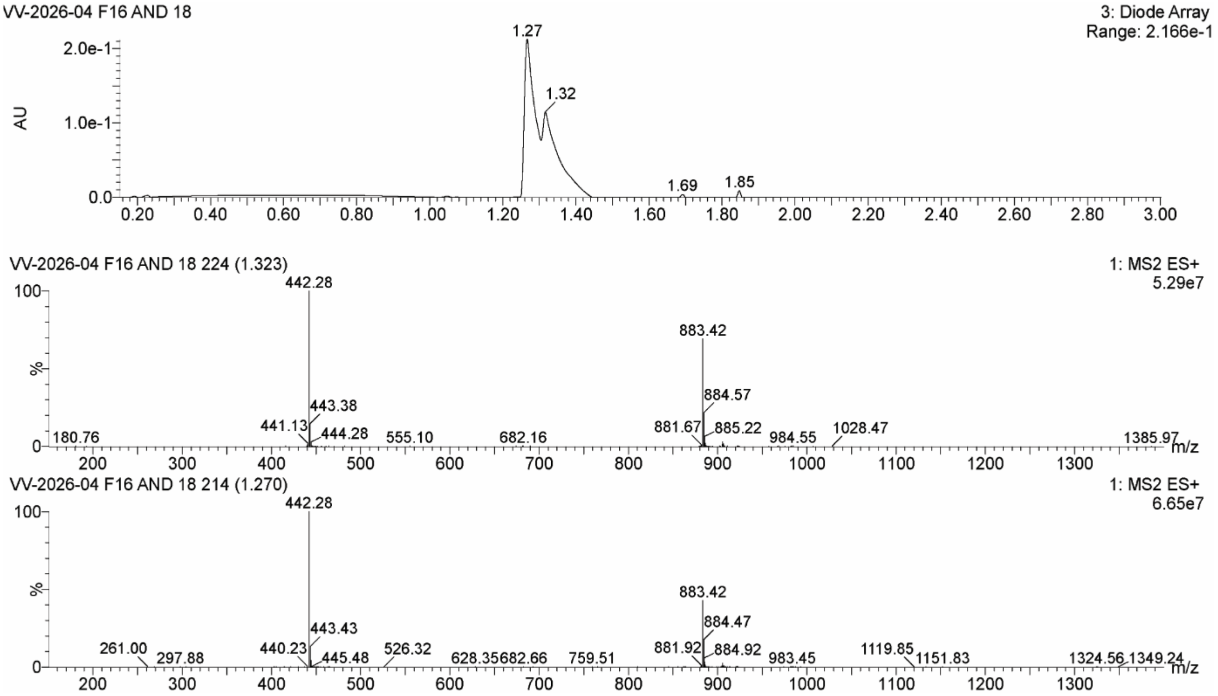

### 7. NMR Spectra

The NMR spectra for compound (1-5) show signal multiplicity consistent with several slowly exchanging conformers. The proline–proline amide bond exists as both cis and trans isomers, and rotation about the carbamate (Cbz) N–CO and phosphoramidate N–P bonds is restricted. Proline ring puckering (endo/exo) further contributes to conformational heterogeneity[89-90].

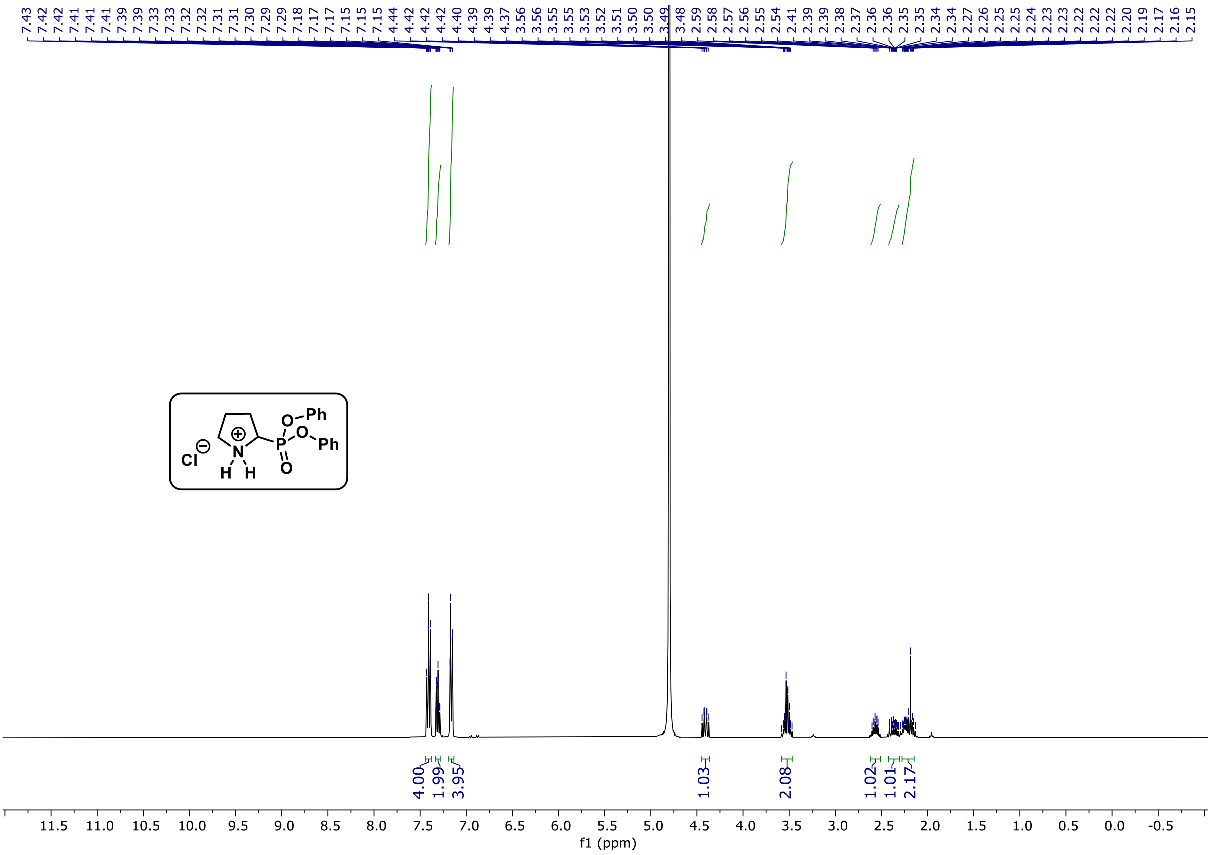


1H NMR spectrum (400 MHz) of **2-(diphenoxyphosphoryl)pyrrolidin-1-ium chloride** at 298 K in $D_2O$.

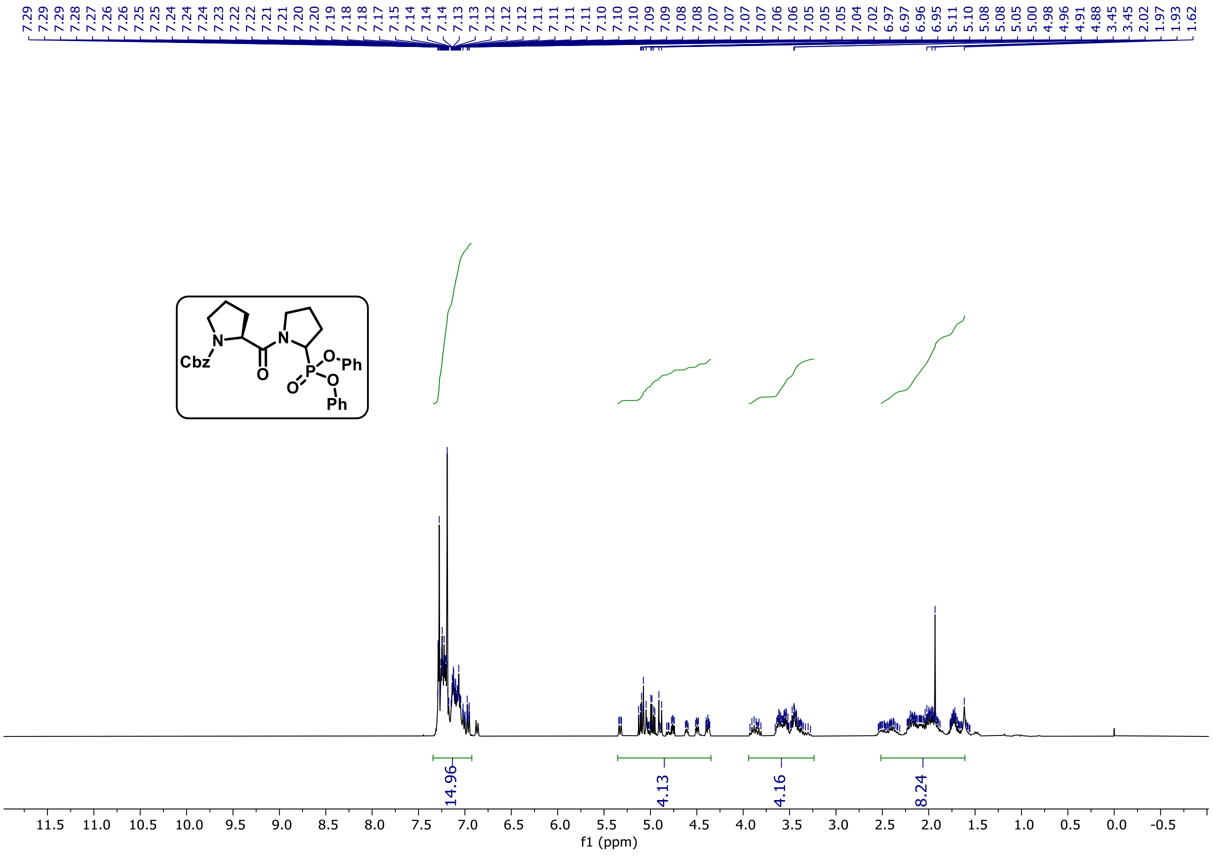

1H NMR spectrum (400 MHz) of **compound 1** (at 298 K in $CDCl_3$)

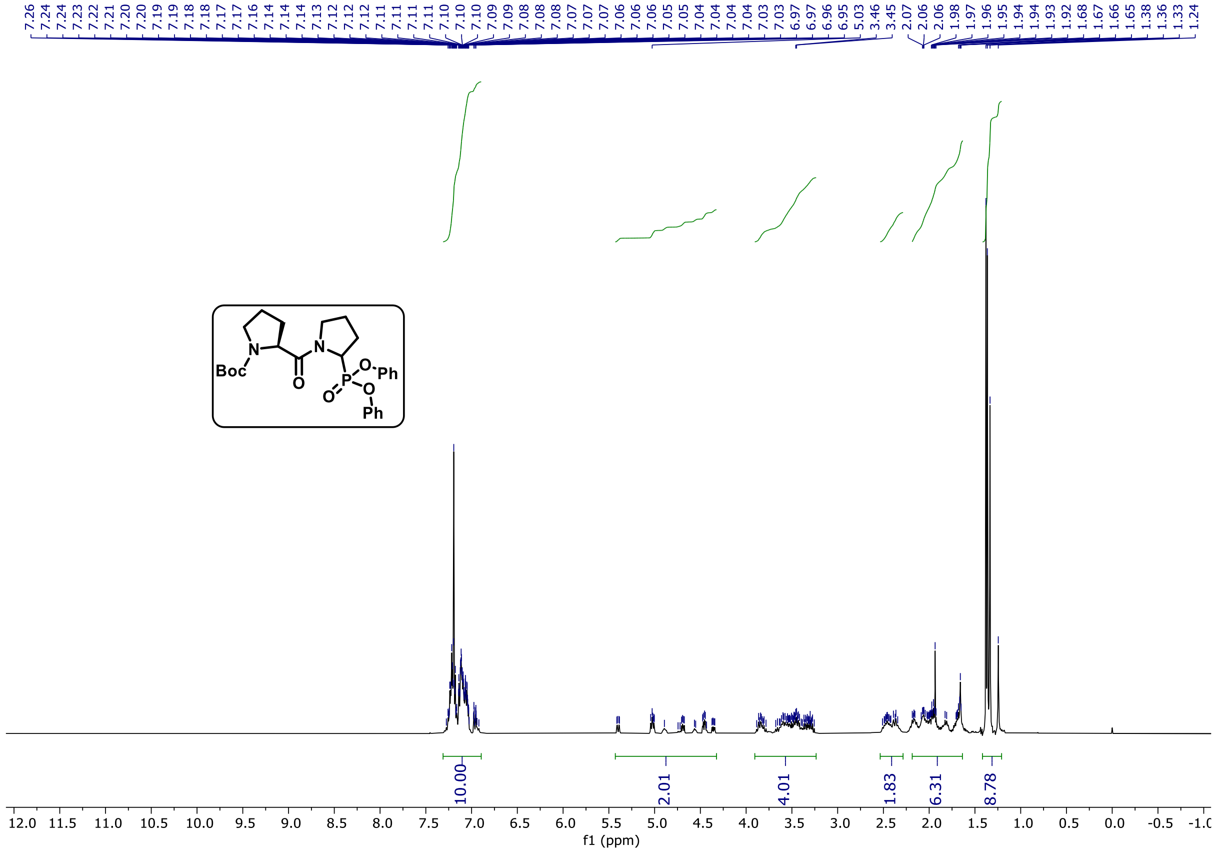

1H NMR spectrum (400 MHz) of **compound 2** (at 298 K in $CDCl_3$)

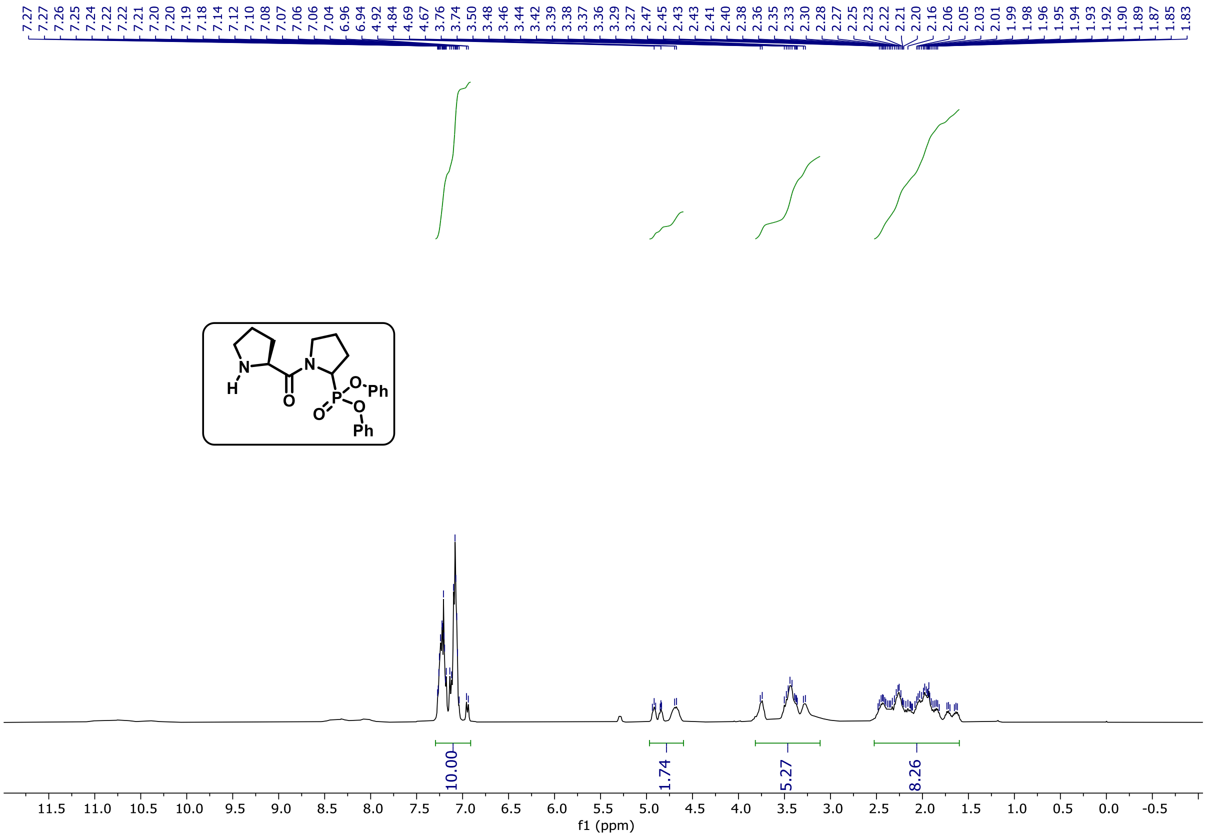

1H NMR spectrum (400 MHz) of **compound 3** (at 298 K in $CDCl_3$)

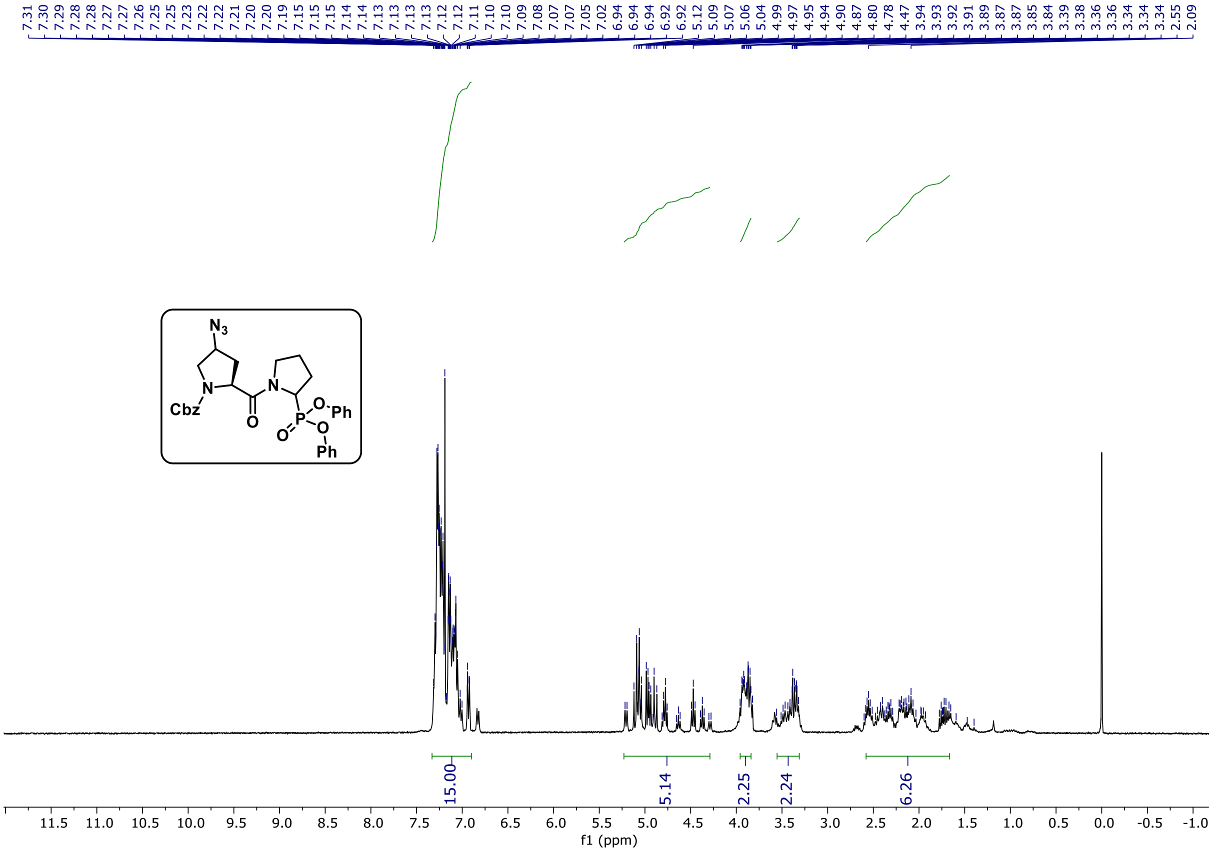

1H NMR spectrum (400 MHz) of **compound 4** (at 298 K in $CDCl_3$)

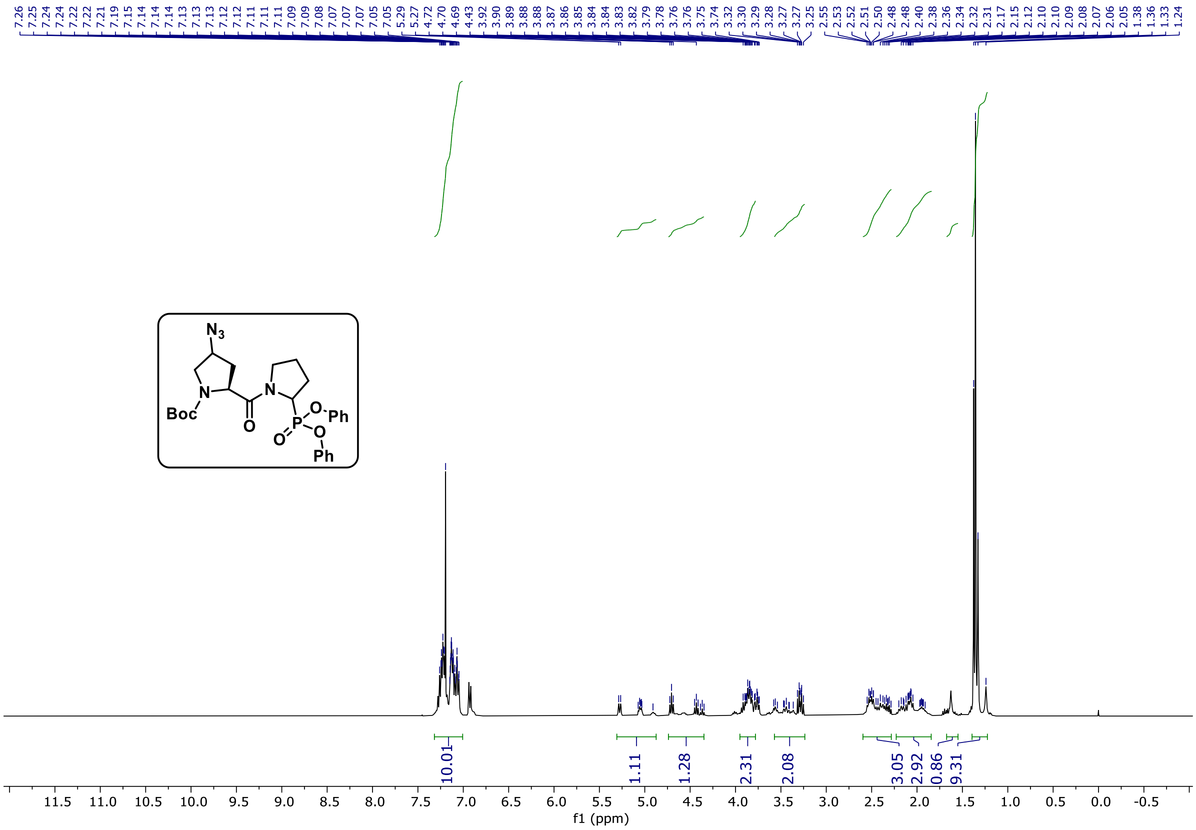


1H NMR spectrum (400 MHz) of **compound 5** (at 298 K in $CDCl_3$)

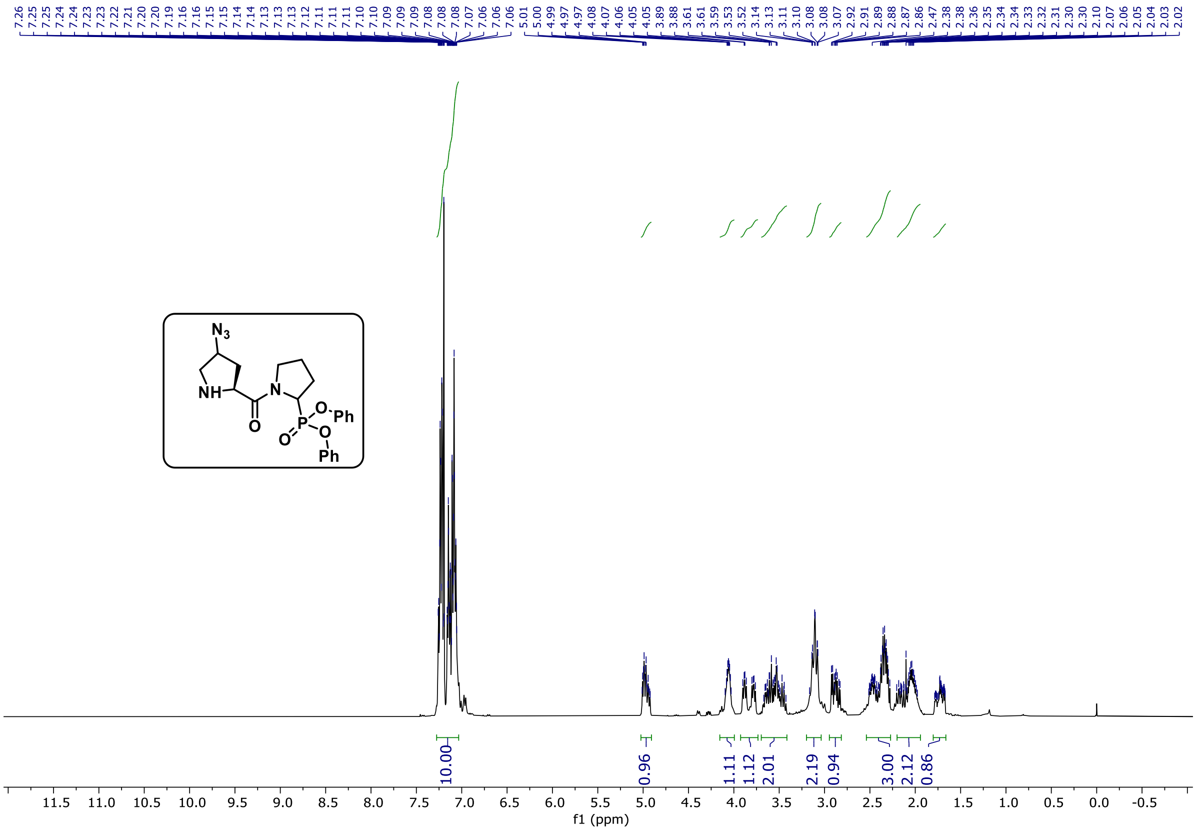


1H NMR spectrum (400 MHz) of **UAMC-936-1** (at 298 K in $CDCl_3$)

### 8. Analytical Data for Unpublished Compounds from the In-House Compound Library

Due to partial degradation of aged library members, compound purity was determined after screening, and only compounds with ≥70% purity were included in the screening.

##### 1) UAMC-105:

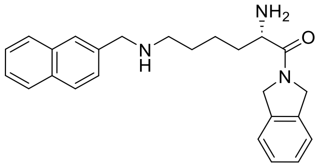

**UPLC:** R*t* 1.10 minutes; (ESI+) m/z calc'd for $C_{25}H_{29}N_3O$ $[M+H]^+$: 388.52; found: 388.20.
**HRMS:** m/z calc'd for $C_{25}H_{29}N_3O$ $[M+H]^+$: 388.2383; found: 388.2388.

UPLC-MS chromatogram and spectrum of **UAMC-105**.

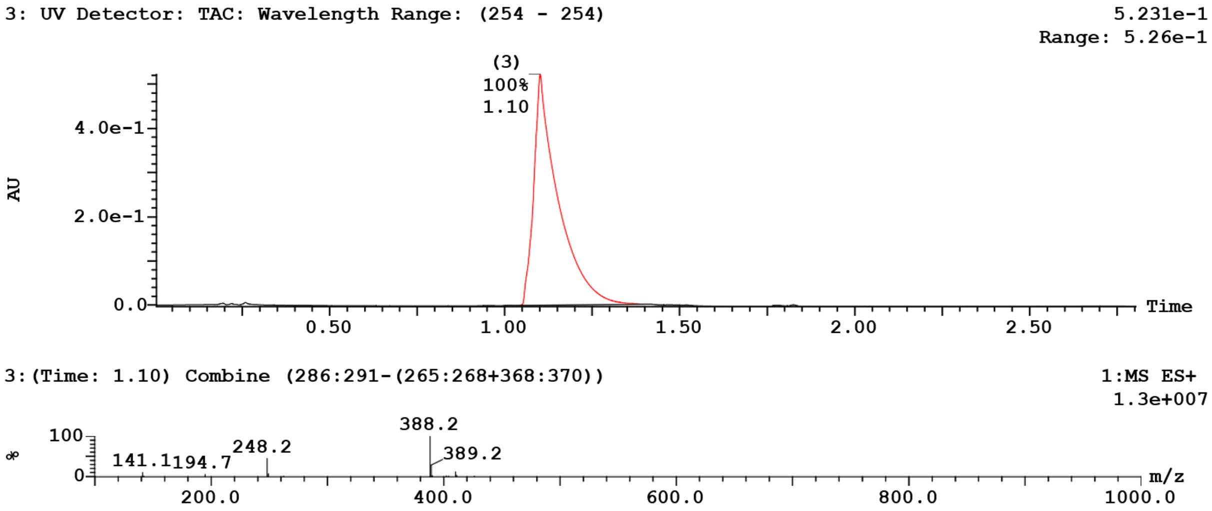


##### 4) UAMC-57:

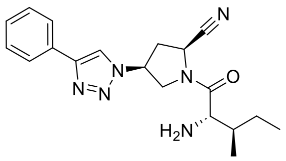

**UPLC:** R*t* 1.27 minutes; (ESI+) m/z calc'd for $C_{19}H_{24}N_6O$ $[M+H]^+$: 353.44; found: 353.20.
**HRMS:** m/z calc'd for $C_{19}H_{24}N_6O$ $[M+H]^+$: 353.2084; found: 353.2080.

UPLC-MS chromatogram and spectrum of **UAMC-57**.

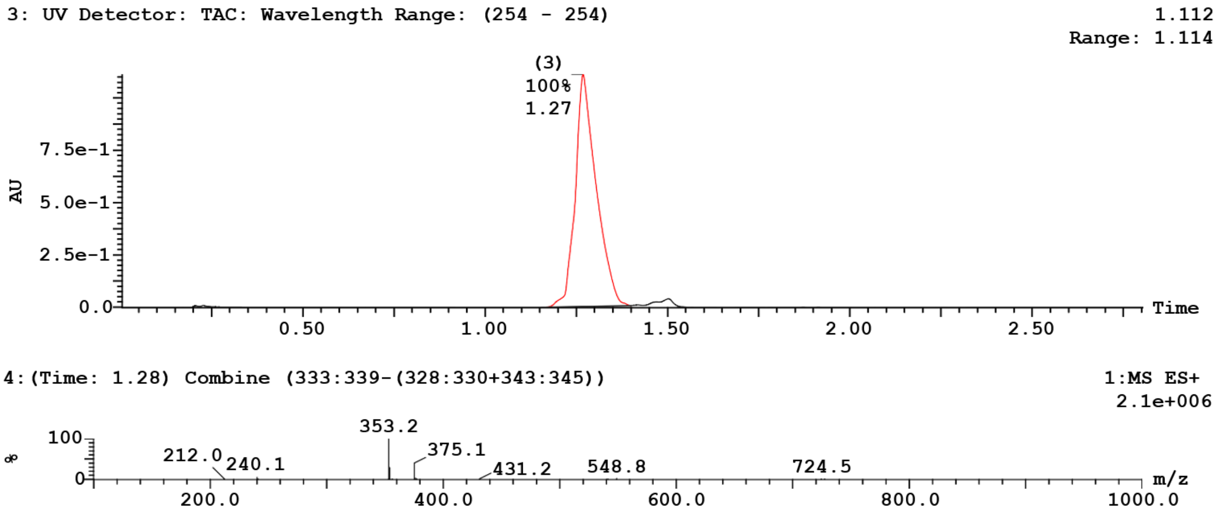


#### 6) UAMC-85:

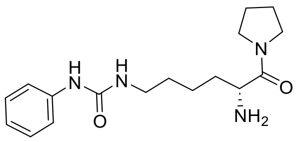


**UPLC:** R*t* 1.12 minutes; (ESI+) m/z calc'd for $C_{17}H_{26}N_4O_2$ [M+H]$^+$: 319.42; found: 319.10.
**HRMS:** m/z calc'd for $C_{17}H_{26}N_4O_2$ [M+H]$^+$: 319.2129; found: 319.2137.

UPLC-MS chromatogram and spectrum of **UAMC-85**.

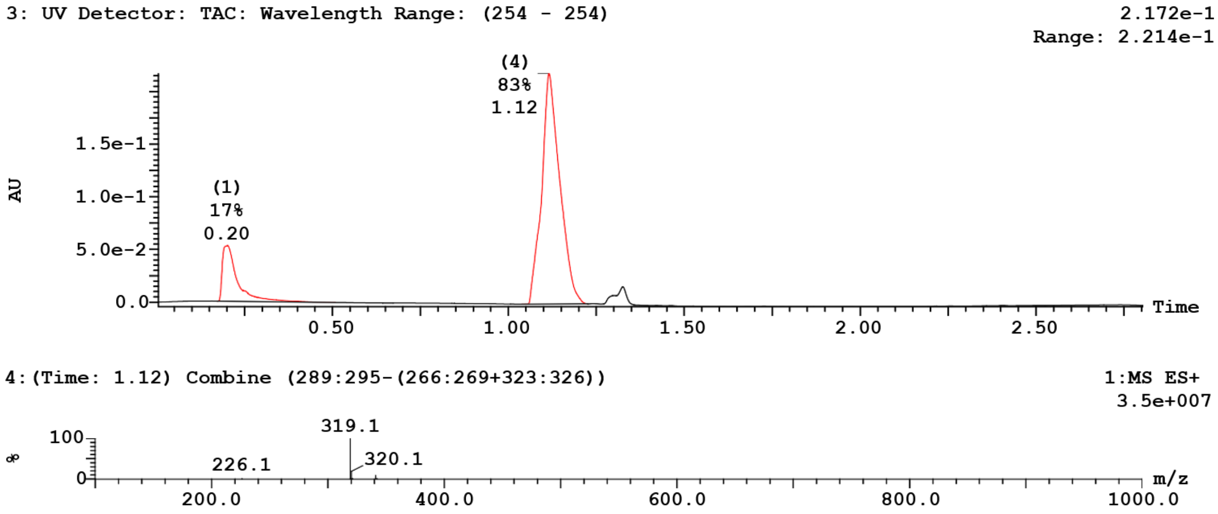


#### 8) UAMC-435:

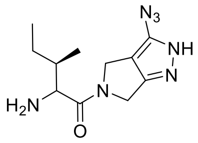

**UPLC:** R*t* 1.07 minutes; (ESI+) m/z calc'd for $C_{11}H_{17}N_7O$ $[M+H]^+$: 264.30; found: 264.10.
**HRMS:** m/z calc'd for $C_{11}H_{17}N_7O$ $[M+H]^+$: 264.1567; found: 264.1573.

UPLC-MS chromatogram and spectrum of **UAMC-435**.

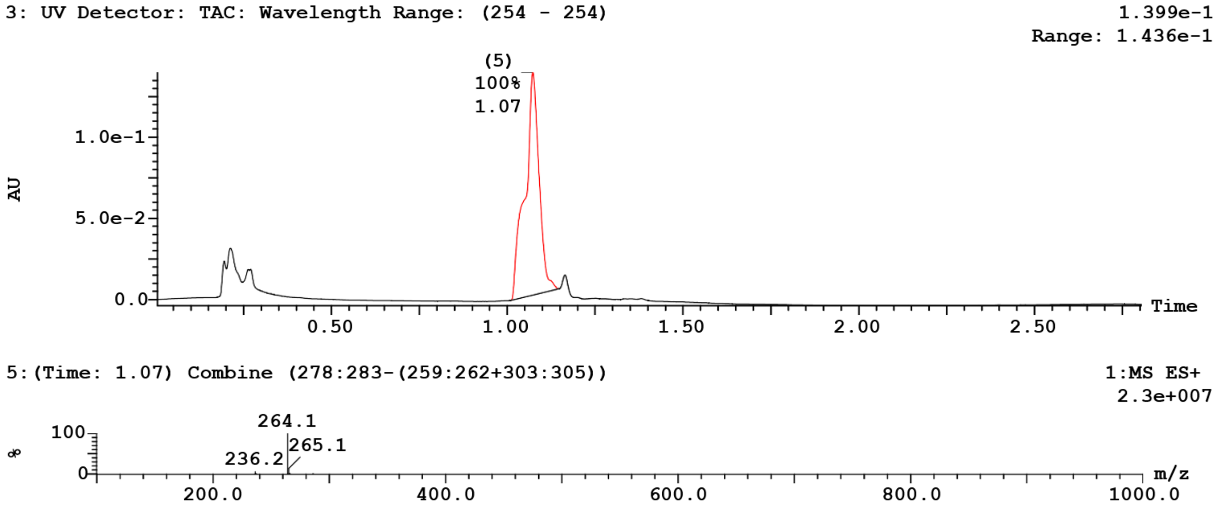


#### 9) UAMC-625:

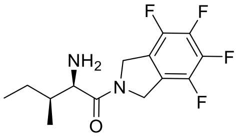


**UPLC:** R*t* 1.29 minutes; (ESI+) m/z calc'd for $C_{14}H_{16}F_4N_2O$ $[M+H]^+$: 305.28; found: 305.00.
**HRMS:** m/z calc'd for $C_{14}H_{16}F_4N_2O$ $[M+H]^+$: 305.1272; found: 305.1272.

UPLC-MS chromatogram and spectrum of **UAMC-625**.

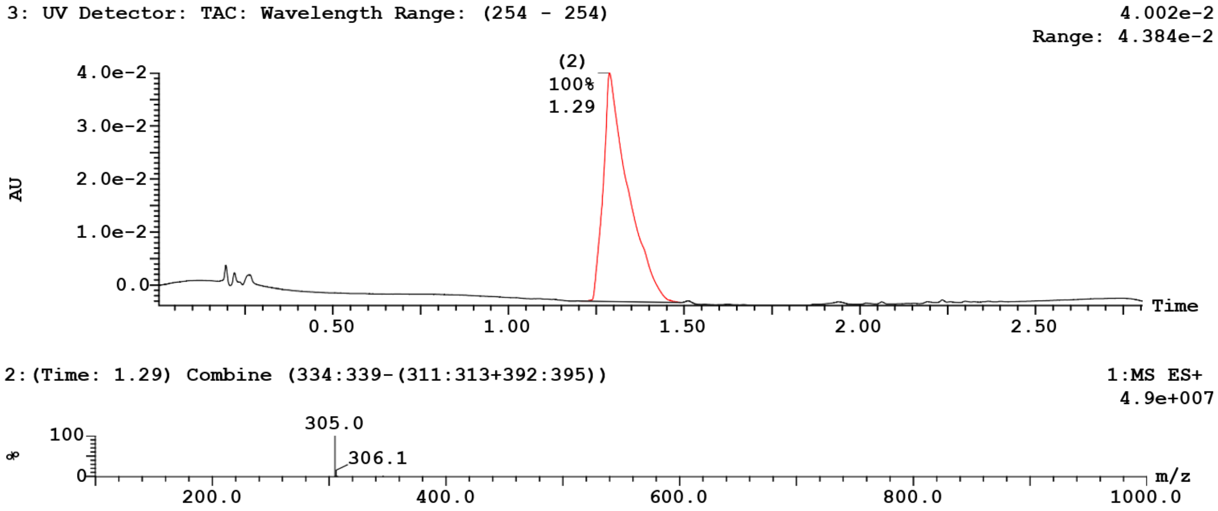

**10) UAMC-649:**

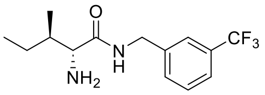


**UPLC:** R*t* 1.27 minutes; (ESI+) m/z calc'd for $C_{14}H_{19}F_3N_2O$ [M+H]$^+$: 289.31; found: 289.10.
**HRMS:** m/z calc'd for $C_{14}H_{19}F_3N_2O$ [M+H]$^+$: 289.1522; found: 289.1525.

UPLC-MS chromatogram and spectrum of **UAMC-649**.

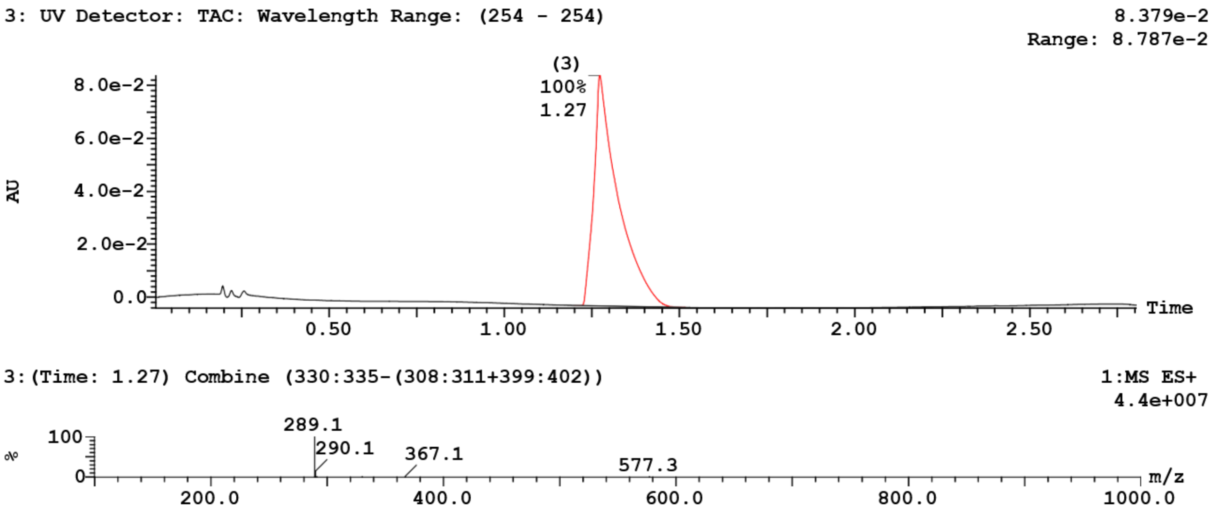


**11) UAMC-651:**

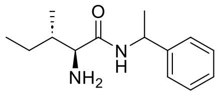


**UPLC:** R*t* 1.16 and 1.18 minutes; (ESI+) m/z calc'd for $C_{14}H_{22}N_2O$ [M+H]$^+$: 235.34; found: 235.20.
**HRMS:** m/z calc'd for $C_{14}H_{22}N_2O$ [M+H]$^+$: 235.1805; found: 235.1810.

UPLC-MS chromatogram and spectrum of **UAMC-651**.

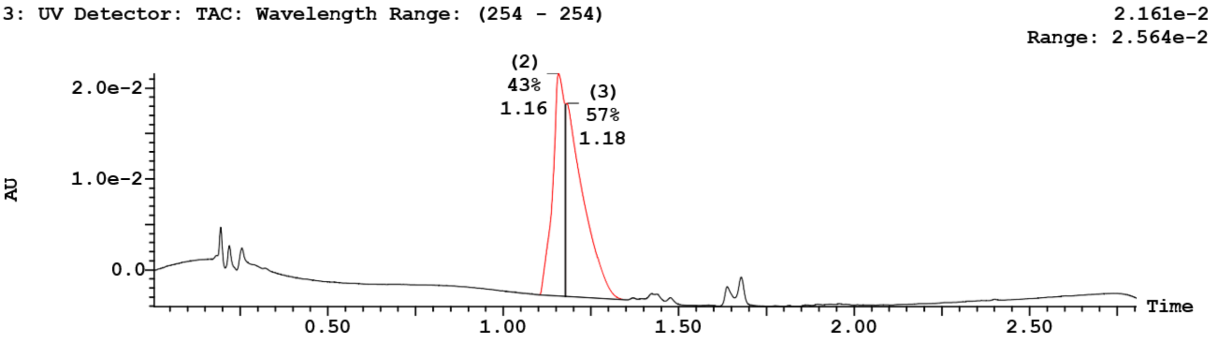

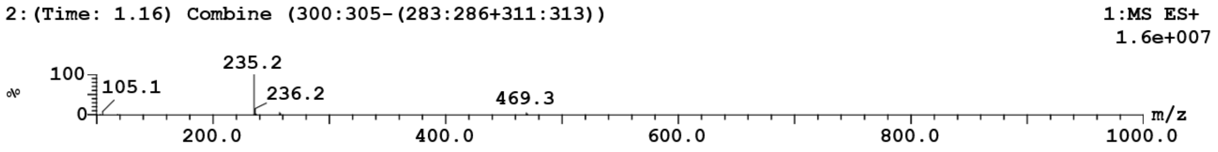


| Peak ID | Compound | Time | Mass Found |
|---|---|---|---|
| 3 | | 1.18 | Not Found |

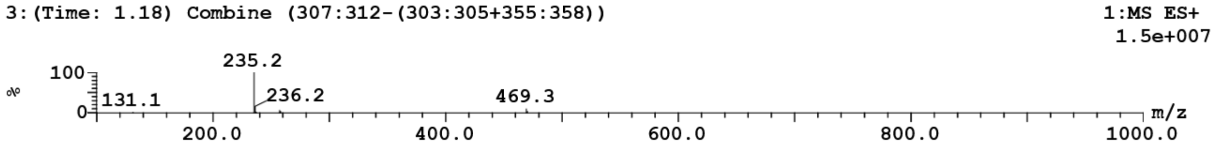


##### 13) UAMC-700:

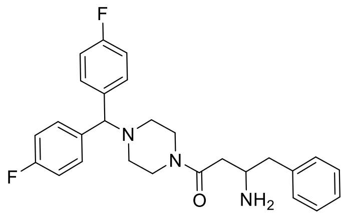


**UPLC:** R*t* 1.56 minutes; (ESI+) m/z calc'd for $C_{27}H_{29}F_2N_3O$ $[M+H]^+$: 450.54; found: 450.20.
**HRMS:** m/z calc'd for $C_{27}H_{29}F_2N_3O$ $[M+H]^+$: 450.2351; found: 450.2357.

UPLC-MS chromatogram and spectrum of **UAMC-700**.

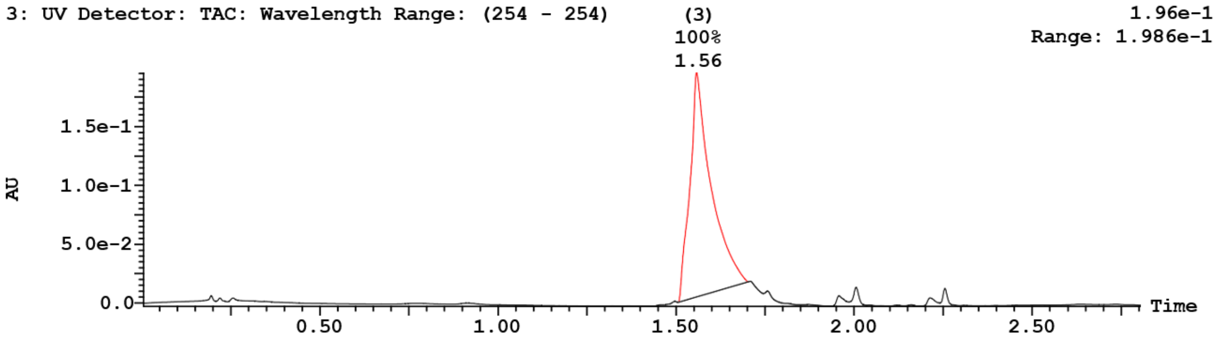


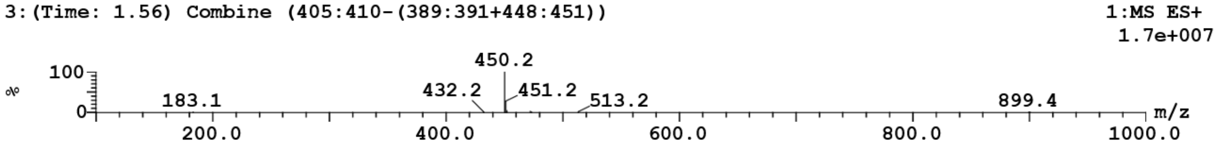


##### 14) UAMC-725:

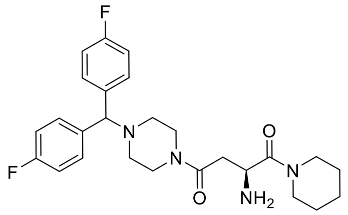


**UPLC:** R*t* 1.51 minutes; (ESI+) m/z calc'd for $C_{26}H_{32}F_2N_4O_2$ $[M+H]^+$: 471.56; found: 471.20.

**HRMS:** m/z calc'd for $C_{26}H_{32}F_2N_4O_2$ $[M+H]^+$: 471.2566; found: 471.2572.

UPLC-MS chromatogram and spectrum of **UAMC-725**.

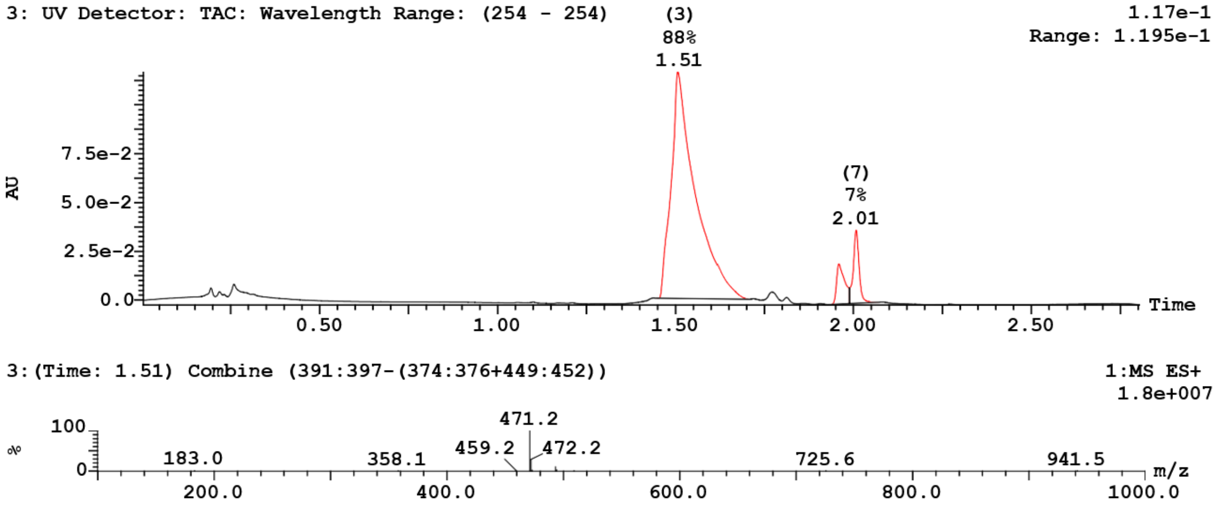


#### 16) UAMC-1431:

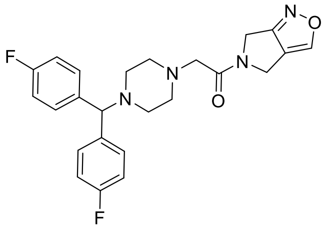


**UPLC:** R*t* 1.40 and 1.44 minutes; (ESI+) m/z calc'd for $C_{24}H_{24}F_2N_4O_2$ $[M+H]^+$: 439.47; found: 439.20.
**HRMS:** m/z calc'd for $C_{24}H_{24}F_2N_4O_2$ $[M+H]^+$: 439.1940; found: 439.1942.

UPLC-MS chromatogram and spectrum of **UAMC-1431**.

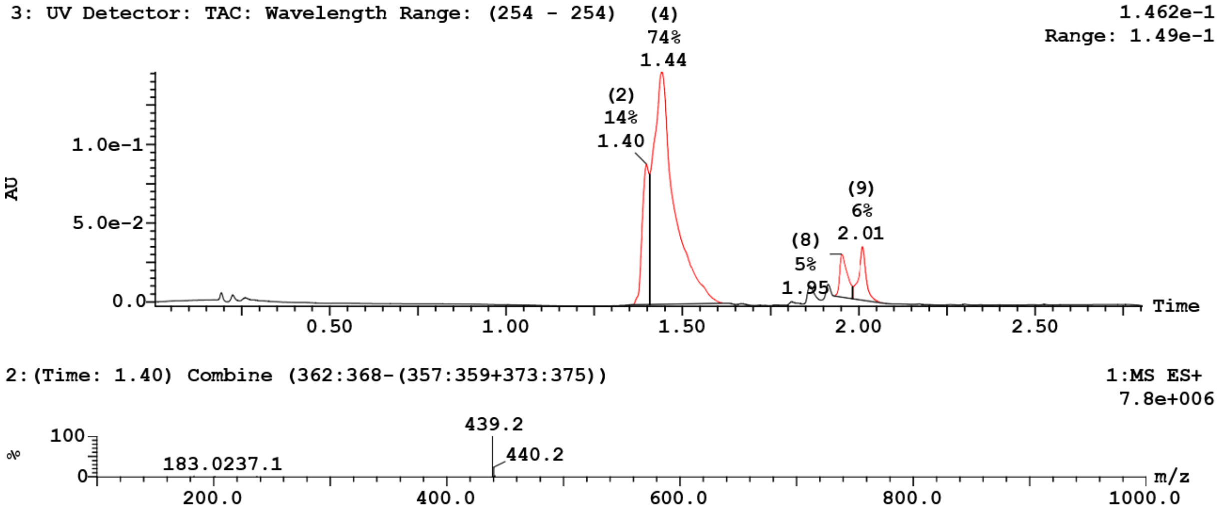

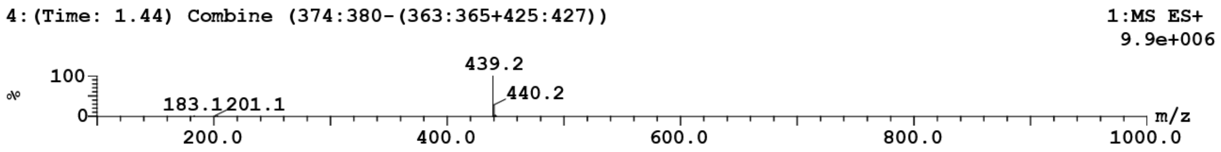


**19) UAMC-1245:**

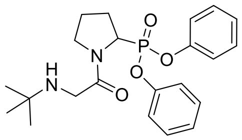

**UPLC:** R*t* 1.38 minutes; (ESI+) m/z calc'd for $C_{22}H_{29}N_2O_4P$ $[M+H]^+$: 417.45; found: 417.20.
**HRMS:** m/z calc'd for $C_{22}H_{29}N_2O_4P$ $[M+H]^+$: 417.1938; found: 417.1941.

UPLC-MS chromatogram and spectrum of **UAMC-1245**.

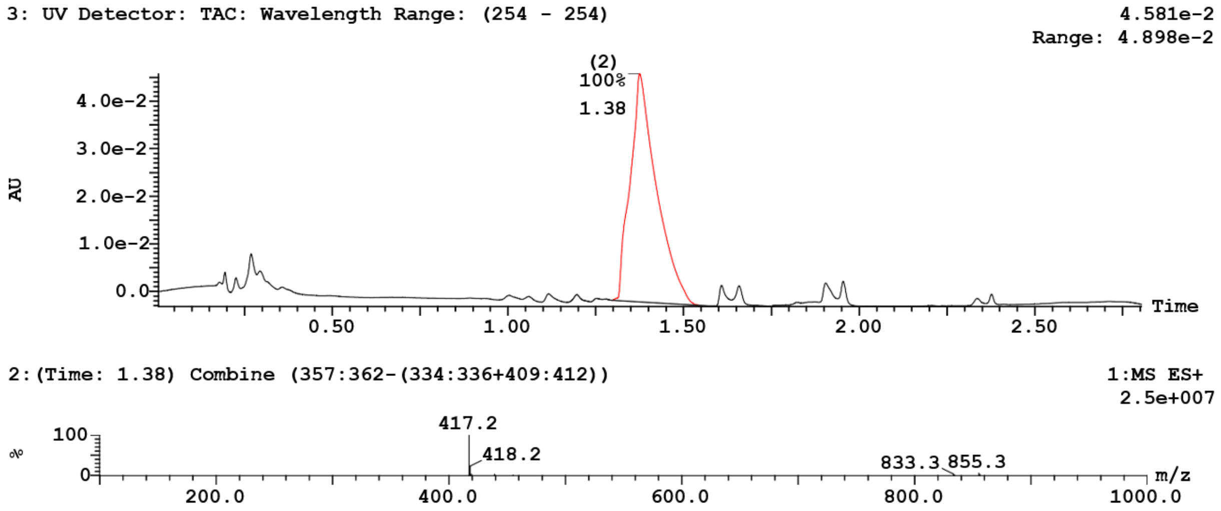


**21) UAMC-3304n:**

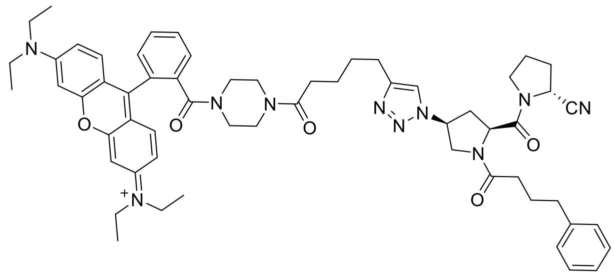

**UPLC:** R*t* 2.04 minutes; (ESI+) m/z calc'd for $C_{59}H_{71}N_{10}O_5$ $[M+2H]^{2+}$: 500.14; found: 500.50.
**HRMS:** m/z calc'd for $C_{59}H_{71}N_{10}O_5$ $[M+H]^+$: 999.5603; found: 999.5598 (1 proton removed to compensate for the positive charge).

UPLC-MS chromatogram and spectrum of **UAMC-3304n**.

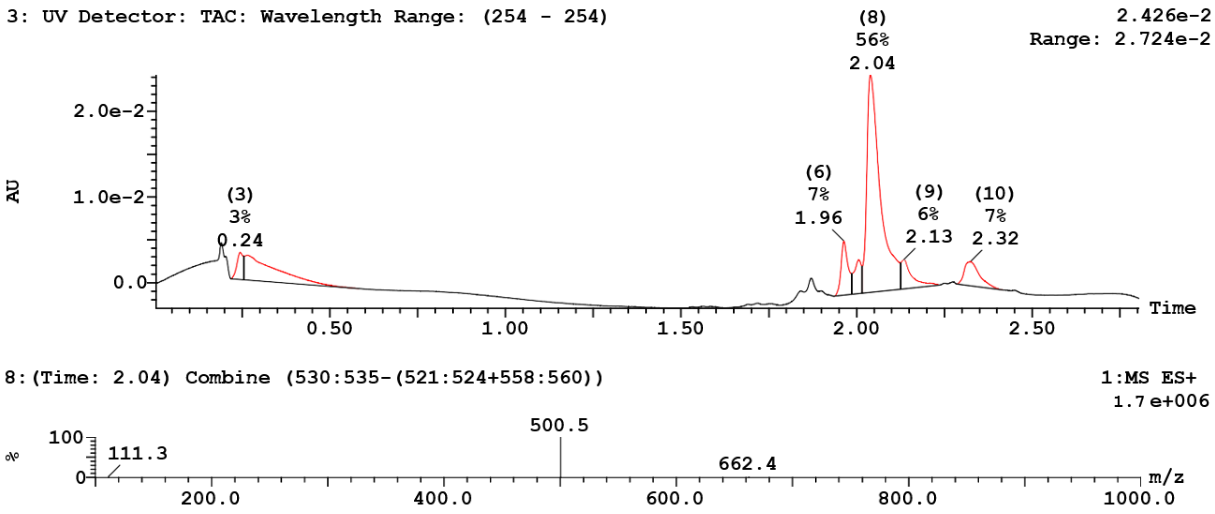
